# Spatial atlas of the ovary identifies molecular events in primordial follicle activation in humans

**DOI:** 10.64898/2026.06.05.730381

**Authors:** Luz Garcia-Alonso, James Ashcroft, Callum Tromans-Coia, Nilay Kuscu, Celeste E Cohen, Pavel V Mazin, Ana Paredes, Hassan Massalha, Cecilia Icoresi Mazzeo, Liz Tuck, Carmen Sancho Serra, Elena Prigmore, Erick Armingol, Gonzalo Martinez, Tong Li, Aljes Binkevich, Alexander V. Predeus, Kevin Troule, Emma Tomlinson, Ana Monteiro, Faranaz Auckburally, Marta Koland, Agnieszka Jaciow, Linlin Wang, Krishnaa T. Mahbubani, Kourosh Saeb-Parsy, Meenakshi Choudhary, Martin Prete, Iva Kelava, Jill Davies, Sheila Lane, Mary Herbert, Virginie Uhlmann, Suzannah A. Williams, Roser Vento-Tormo

## Abstract

The human ovarian reserve is established prenatally, when granulosa cells encapsulate germ cells to form a pool of quiescent primordial follicles, that ultimately determines the female reproductive and endocrine lifespan. From birth to menopause, subsets of these quiescent follicles, located in the thin outer ovarian cortex, are activated and undergo a growth programme with progressive inward migration to the inner cortex, before either undergoing atresia or, during reproductive age, ovulation. Disruption of this process may lead to infertility, metabolic disorders and early menopause, yet early follicle development remains largely poorly understood. Here, we generate the most comprehensive single-cell and spatial multiomics atlas of the human pre- and postnatal ovarian cortex, integrating transcriptomic and chromatin profiles from over four million cells obtained from fetal and newly profiled pediatric and adult donors. We resolve the early granulosa cell trajectory at unprecedented resolution and identify a retinoic acid-associated regulatory switch accompanying follicle activation. We further show that stromal fibroblasts are not homogeneous, but instead form a dynamic scaffold establishing previously unrecognised morphogen and paracrine gradients that organise the cortex into functional niches supporting quiescent, growing, and atretic follicles. Finally, we identify ovarian lipid associated macrophages (oLAMs) that localise around follicles and are likely to support tissue remodelling during folliculogenesis. Together, this atlas provides a foundational blueprint for human ovarian development and homeostasis, and provides a framework for improving strategies in fertility preservation and in vitro follicle maturation.

## Introduction

The ovaries are central to female reproductive capacity and systemic endocrine regulation, orchestrating essential physiological processes across the lifespan^1^. Their fundamental functional units, the follicles, consist of specialised granulosa cells surrounding the oocyte. Follicles are highly dynamic structures that undergo sequential stages of differentiation after activation. In humans, primordial follicles form prenatally beginning at ∼17 post-conceptional weeks (PCW), establishing a finite ovarian reserve before birth^2,3^. Postnatally, these quiescent primordial follicles reside predominantly within the thin (∼1 mm) outer ovarian cortex^4^. From their formation until menopause, subsets of primordial follicles periodically exit dormancy and progress through the primary, secondary (preantral) and early-antral stages in a process termed early, gonadotropin-independent follicle development^2^. Later follicle maturation through the pre-ovulatory, ovulatory and subsequent formation of *corpus luteum,* which are the primary source of estrogen and progesterone, occurs only after puberty in response to cyclic pituitary gonadotropins (FSH and LH)^5^. As follicles grow, they undergo directional migration from the outer towards the inner cortex, acquiring additional somatic support from surrounding cell types, including theca cells, which are essential for follicle maturation and steroidogenesis^6^. The majority of follicles, however, do not reach ovulation but instead undergo atresia, a controlled degenerative process accompanied by immune-mediated clearance and stromal remodeling^7,8^. Despite the central importance of follicles development for women’s health, the molecular mechanisms that maintain the ovarian reserve, regulate early gonadotropin-independent follicle development, and coordinate tissue repair following atresia remain poorly understood in humans^9–11^.

Recent advances in single-cell genomics, including our own work, have begun to chart the cellular composition of the human ovary in development^12–16^ and adulthood^17–21^. However, several critical gaps remain. First, the primordial follicle reserve is under-represented in existing adult datasets owing to limited sampling of the outer cortex and the absence of pediatric stages. This limitation is particularly consequential given the rapid attrition of the primordial pool during childhood, which declines from ∼1 million follicles at birth to ∼300,000 by puberty through early follicle activation and growth^22,23^, rendering primordial follicles exceedingly rare in adult ovaries. Second, existent single-cell studies rely solely on the transcriptome and lack chromatin accessibility data, which is critical for resolving the regulatory programmes that govern cellular transitions. Third, most studies have focused on advanced growing isolated follicles^24^, whereas the surrounding stromal compartment, particularly the outer cortex, has been characterised primarily at histological or biophysical resolution^24–27^. While biomechanical differences and perifollicular zones have been described^28,29^, the underlying cellular heterogeneity, especially among stromal fibroblasts populations, is likely to remain unresolved without comprehensive spatial approaches^20,30^. Spatially resolved single-cell atlases are therefore required to understand how the cortex is functionally organised, where follicles at distinct differentiation stages (from quiescence through growth to atresia) co-exist and respond to their local signalling environments.

Defects in these processes have profound clinical consequences. Approximately 3.5% of women under the age of 40 years experience primary ovarian insufficiency, leading to early menopause, infertility, and accelerated biological aging^11,31^. Although this condition can arise from genetic causes or iatrogenic injury, most commonly gonadotoxic chemotherapies, the majority of cases remain idiopathic^32^. In adults, controlled ovarian stimulation followed by oocyte or embryo cryopreservation provides an effective fertility-preservation strategy. In contrast, for pediatric donors, ovarian tissue cryopreservation (OTC) of the outer cortex with subsequent autotransplantation remains the only clinically feasible option^33–37^. While transplantation can restore endocrine function and delay menopause, fertility outcomes following OTC remain limited compared with oocyte preservation^38,39^, and remain poorly characterised in paediatric donors. Moreover, in certain hematological malignancies, such as leukemia, autotransplantation carries a risk of reintroducing malignant cells residing within the ovarian cortex^40^. Follicle culture with *in vitro* maturation would offer a safer alternative^41^, however optimising this approach requires understanding how early follicles mature within their native cortical niche. A deeper understanding of the cellular and spatial organisation of the outer ovarian cortex can thus guide *in vitro* strategies for pediatric fertility preservation.

Here, we present the most comprehensive, regionally resolved, single-cell multimodal atlas of the human ovarian cortex across the lifespan to date. Our dataset includes 30 newly profiled cortical samples spanning pediatric from birth through puberty, together with 10 adult ovaries. By integrating single-cell transcriptomics, chromatin accessibility, spatial multi-omics, and histomorphology, we generate more than four million high-quality cellular profiles spanning prenatal development through reproductive age. This resource provides a robust reference framework that we use to contextualise and harmonise existing ovarian atlases. Cross-modality integration enables high-resolution reconstruction of granulosa cell trajectories throughout follicle development and identification of molecular regulators underlying this process. Coupling spatial gene expression with tissue histology further reveals higher-order architectural organisation of the ovarian cortex that was unappreciated in previous single-cells studies, including fibroblast strata that spatially transition across niches to establish local signaling hubs. Collectively, this atlas provides the highest resolution reference to date of human ovarian cortical cell states, interactions, and spatial organisation, and forms a foundational resource for understanding ovarian physiology, dissecting the cellular basis of reproductive disorders, and advancing cortical tissue preservation as well as *in vitro* maturation protocols.

## Results

### Spatial single-cell atlas of the human ovarian cortex

To characterise the cellular landscape of the human ovarian cortex across the lifespan, we generated a spatial multimodal atlas from freshly collected ovarian cortical strips from infants, prepubertal and pubertal children (1-15 y/o, n=31; chemotherapy-naïve children undergoing fertility preservation), as well as ovarian biopsies from adult donors (19-46 y/o, n=6), whole ovaries from oophorectomies (37-44 y/o, n=3) and a donor deceased from non gynecological conditions (30y/o, n=1; **Figure 1a-b; Supplementary Table 1**). For additional stages of follicle development, we included visible follicles (0.5-2 mm) manually isolated from pediatric cortical remnants (n=6; **Figure 1a-b; Supplementary Figure 1a**).

**Figure 1.**
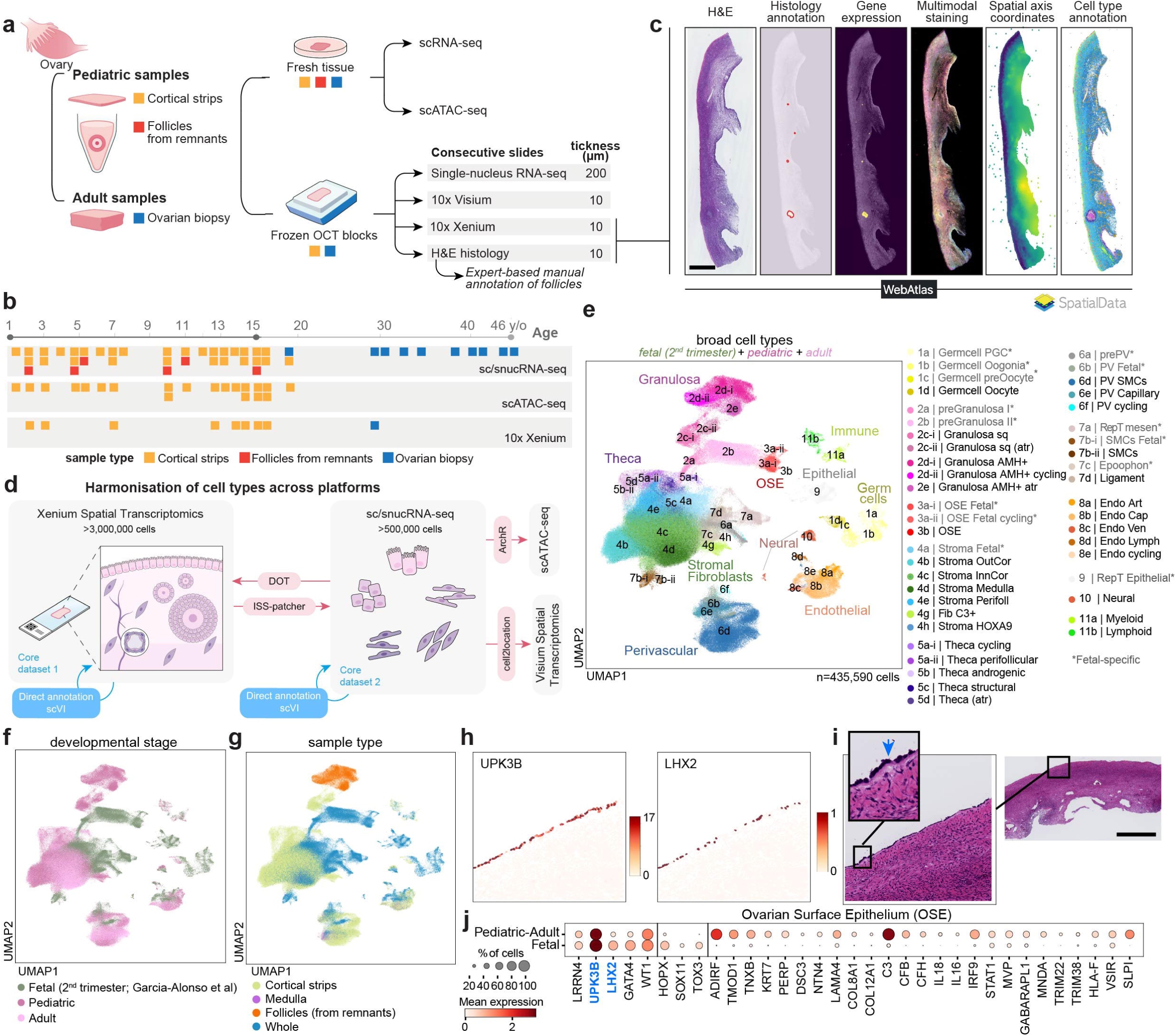
Study overview. **a.** Schematic illustration of samples processing workflow. Color indicates sample type (“orange” for cortical strips, “red” for follicles from the inner cortex remnants, and “blue” for ovarian biopsies). **b.** Diagram summarising the age (x-axis) and sample type composition (color) of our donor cohort along with the technologies used to characterise the donors. **c.** Example Xenium section (AU14; 3 y/o) showing registered cell images and molecular data modalities, available through the WebAtlas web portal and as a SpatialData object. Scale bar = 1mm. **d.** Schematic representation of the pipeline to define and harmonise cell type annotations across data modalities. **e-g.** Batch-corrected Uniform Manifold Approximation and Projection (UMAP) embedding of the scRNA-seq dataset (n = 435,590 cells; n=50 donors) coloured by broad cell type annotation (“e”), developmental stage (“f”), and sample type (“g”). **h.** Measured gene expression of *UPK3B* and *LHX2* in a representative cortical strip sample (AU30; 14y/o) containing Ovarian Surface Epithelium (OSE) cells. **i.** Hematoxylin and eosin (H&E) staining of the section adjacent to panel “h”, shown at both high and low magnification. Scale bar in low magnification = 1,000µm. **j.** Dotplot showing the log-transformed, min-max normalised expression of selected marker and differentially expressed genes (x-axis) between prenatal and pediatric+adult OSE cells (y-axis).

**Figure 2.**
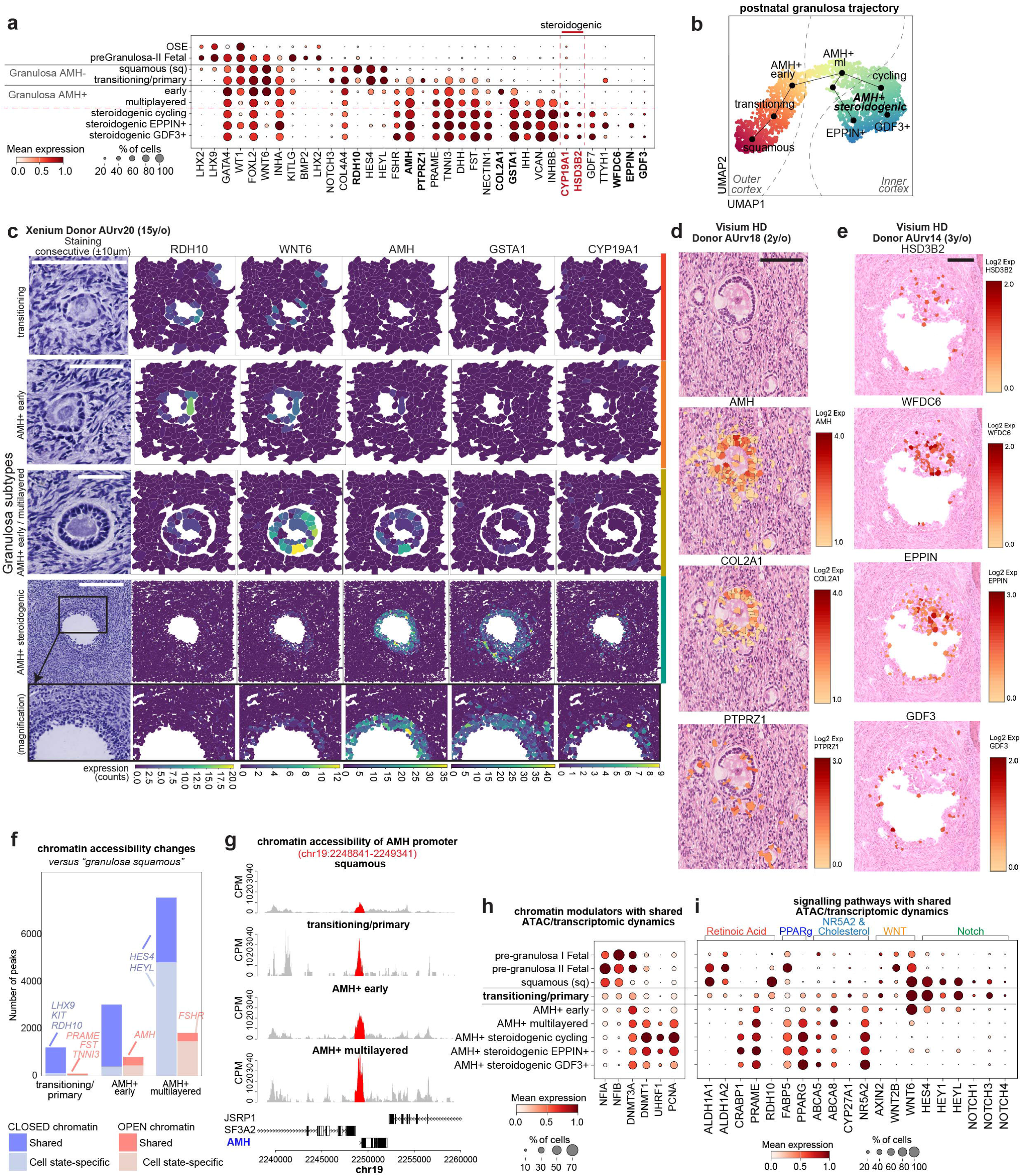
Molecular changes along early granulosa differentiation. **a.** Dotplot showing the log-transformed, min-max normalised expression of selected marker genes (x-axis) for prenatal and pediatric-adult granulosa cell states (y-axis; fine-level annotation). Blue gene names denote literature marker genes. **b.** Batch corrected force directed graph (FDG) visualisation of granulosa cells (downsampled to n=450 cells per category; n=3,134 cells; n=27 donors) from pediatric age donors coloured by pseudotime and labelled with fine-level annotations. Trajectories reconstructed with Slingshot are overlaid on the embedding. Atretic granulosa cells are not included. **c.** Panel showing representative follicle stages captured in Xenium spatial transcriptomic sections, with adjacent histology (first column), multiplex cell staining (second column), and log-transformed expression of selected granulosa state markers (*RDH10, WNT6, AMH, GSTA1, CYP19A1*). The bottom row shows a higher magnification of the antral follicle in the row above. Scale bar = 100µm. **d.**Visualisation of selected marker transcripts in an ovarian cortex section from a 2y/o donor containing primordial, transitioning and multilayered follicles, profiled using Visium HD. Gene panels show log-transformed expression of AMH, COL2A1 and PTPRZ1. Scale bar = 100 µm. **e.** Visualisation of selected marker transcripts in an ovarian cortex section from a 3y/o donor containing one antral follicle, profiled using Visium HD. Gene panels show log-transformed expression of HSD3B2 (steroidogenesis), WFDC6, EPPIN and GDF3. Scale bar = 100 µm. **f.** Barplots showing the number of differentially accessible chromatin peaks (FDR significance < 0.01) in each granulosa cell state relative to quiescent squamous granulosa. Red indicates increased accessibility and blue indicates decreased accessibility; lighter shades reflect state-specific changes, while darker shades represent changes shared across states. **g.** Genome accessibility tracks showing peak accessibility around the AMH locus (chr19: 2,248,950-2,249,450), highlighting increased accessibility in transitioning granulosa cells. **h-i.** Dotplot showing the log-transformed, min-max normalised expression (y-axis) of chromatin remodelers (“g”) and selected components of signalling pathways (“h”), respectively, along prenatal and pediatric-adult granulosa cell states (x-axis; fine-level annotation).

Samples underwent two complementary workflows (**Figure 1a**). Fresh tissue was enzymatically dissociated and profiled using single-cell RNA sequencing (scRNA-seq; **Supplementary Figure 1**) and single-cell chromatin accessibility (scATAC-seq; **Supplementary Figure 2**). In parallel, tissue fragments were snap-frozen in OCT, cryosectioned, and processed on alternating slides for spatial transcriptomics (10x Visium HD and Xenium using the bespoke probe panel in **Supplementary Table 2; Supplementary Figure 3**), H&E histology (**Supplementary Figure 3a,f**), and single-nucleus RNA-seq (snRNA-seq; **Figure 1c, Supplementary Figure 4**). H&E sections were manually and computationally annotated, aligned to their corresponding transcriptomic profiles, and registered into a combined multimodal dataset (**Figure 1c**; **Supplementary Note 1; Supplementary Figure 3f**), enabling large-scale mapping of tissue morphology with spatially resolved transcriptional profiles.

To obtain a comprehensive lifespan-resolved view, we integrated dissociated single-cells and nuclei with second-trimester whole-ovary single-cell data previously generated by our group^12^ (**Figure 1d-g; Supplementary Figure 1b,4a**). Cell lineages and state identities were then assigned across dissociated cells, nuclei, and Xenium-detected cells using canonical markers and spatial context (**Supplementary Figures 1d,3d,4e**). Initial annotations were generated from single-cell transcriptomics data and refined via spatial validations, particularly for granulosa states, which were cross-referenced with histologically defined follicle stages. We harmonised annotations across all platforms (**Figure 1d; Supplementary Note 2**), including scATAC-seq (**Supplementary Figure 2**). Annotations can be visualised and queried at https://www.reproductivecellatlas.org/ovaries/sanger_cells/.

**Figure 3.**
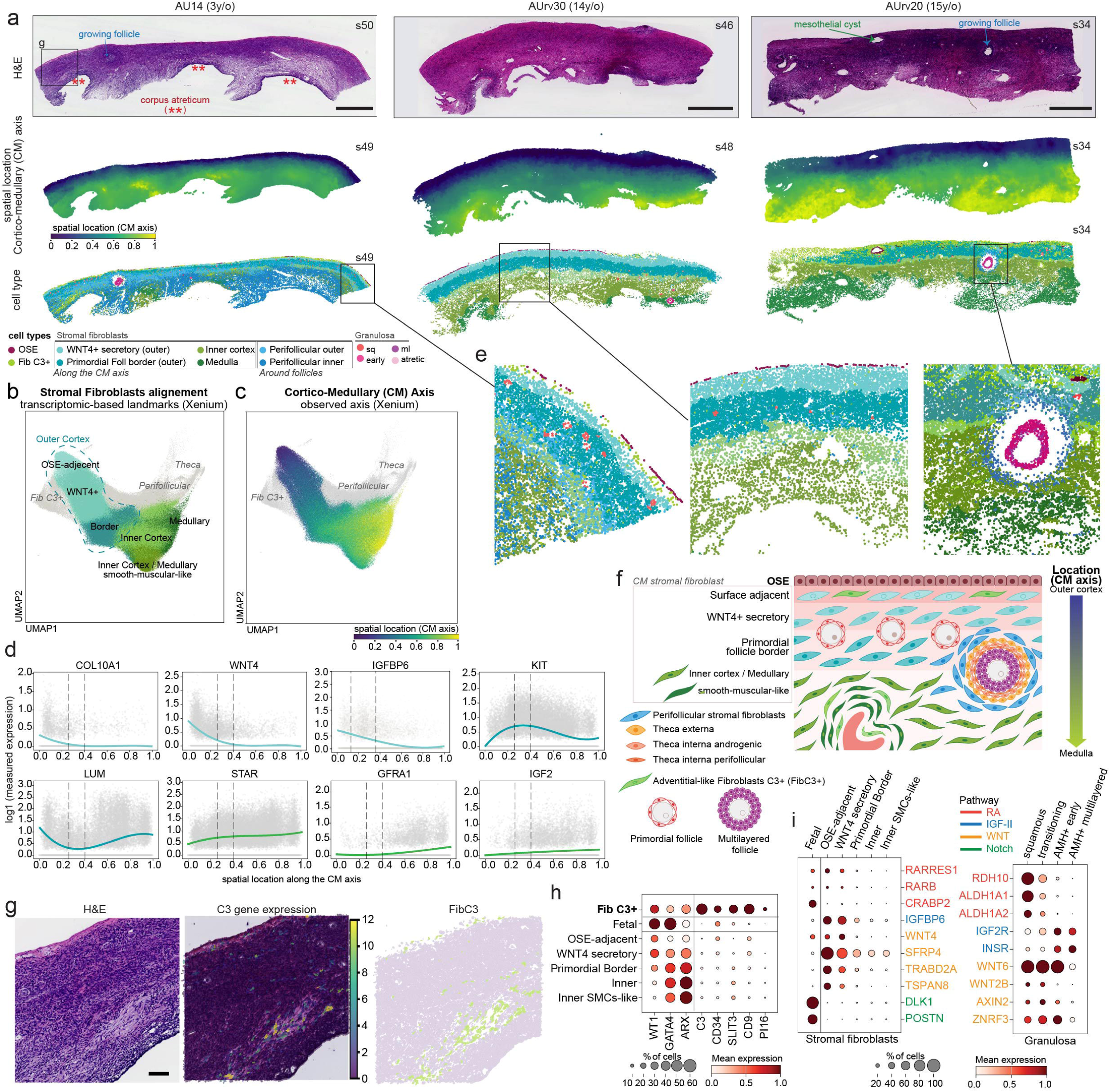
Fibroblast architecture of the ovarian outer cortex. **a.** Three representative ovarian cortical strip sections from pediatric donors (AU14 3y/o; AU30 14y/o and AU20 15y/o) profiled by Xenium, showing adjacent Hematoxylin and eosin (H&E) staining histology (top row), “*Cortico-Medullary axis”* location inferred from the *OrganAxis*-inspired approach (middle row), and cell type annotations for stromal fibroblasts and ovarian surface epithelium (OSE). Scale bar = 1mm **b-c.** Batch-corrected Uniform Manifold Approximation and Projection (UMAP) embedding of the Xenium stromal fibroblast and theca cells coloured by broad cell type annotation, and *“Cortico-medullary Axis”* location inferred from *OrganAxis*-inspired approach (n = cells). **d.** Smoothed splines of Xenium-measured expression (log-transformed) of selected genes spatially dynamic along the “*Cortico-medullary Axis*” in stromal fibroblasts cells. **e.** Higher magnification of selected regions from the sections represented in “a”. **f.** Schematic illustration summarising the location of the different stromal fibroblast subtypes along the “*Cortico-medullary Axis*” together with other ovarian cell types. **g.** Hematoxylin and eosin (H&E) staining (left), gene expression for gene *C3* (middle) and cell type annotation of C3+ Fibroblasts (Fib C3+; right) of a representative pediatric cortical strip section (AU14; 3y/o, low magnification in panel “a” left column sample) profiled by Xenium spatial transcriptomics. Scale bar = 100µm. **h.** Dotplot showing the log-transformed, min-max normalised expression of selected Fib C3+ marker genes (x-axis) in prenatal and pediatric+adult ovarian stromal fibroblasts (y-axis). **i.** Dotplots showing the log-transformed, min-max normalised expression of selected genes (y-axis) in prenatal and pediatric+adult fibroblasts and granulosa cells (x-axis). Genes are colored by their associated signalling pathway.

**Figure 4.**
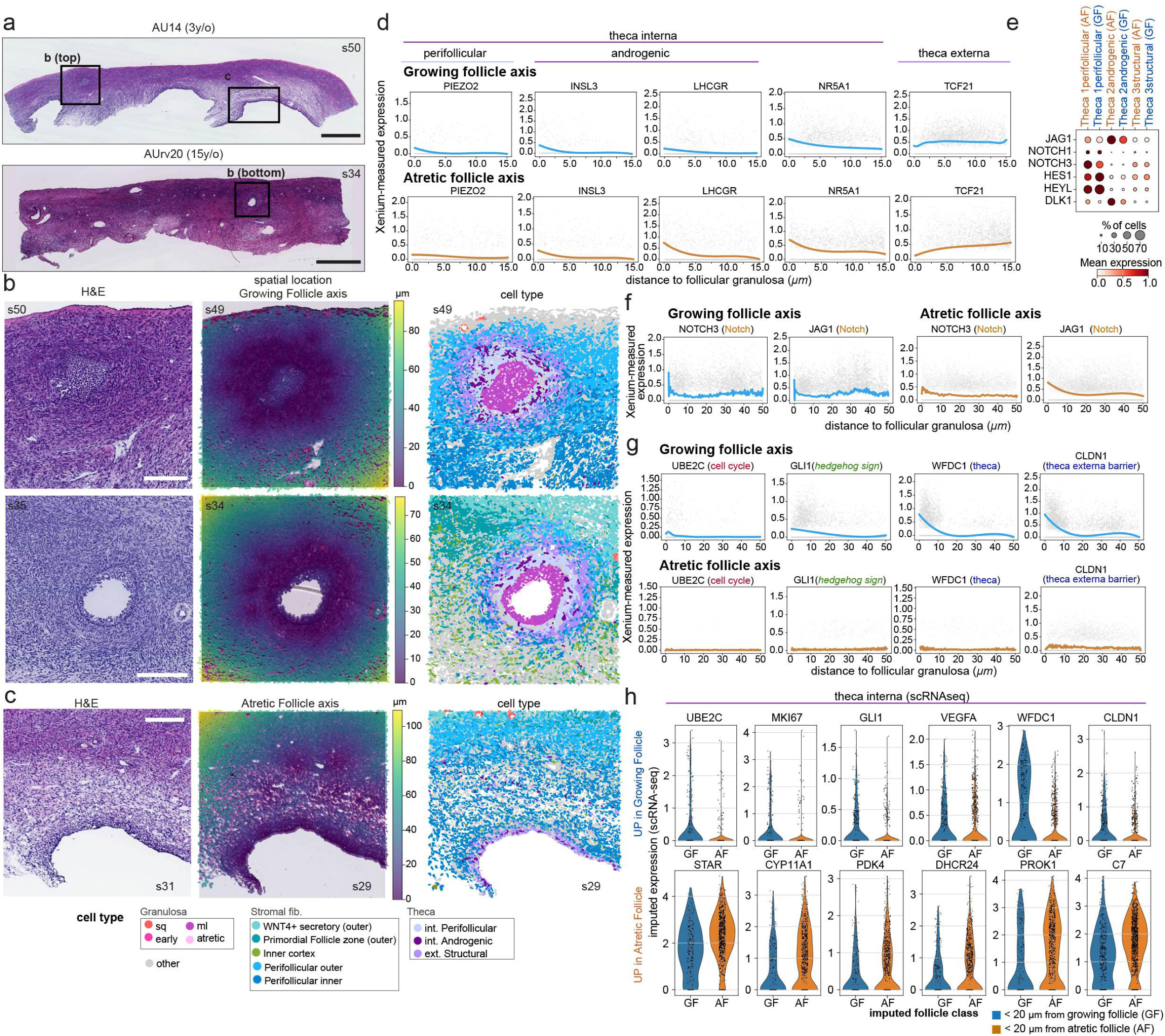
Perifollicular niche characterisation during growth and atresia. **a.** Hematoxylin and eosin (H&E) staining of two representative ovarian cortical strip sections from pediatric donors (AU14 3y/o and AU20 15y/o) containing growing antral and atretic follicles profiled by Xenium. Scale bar = 1mm. **b.** Magnified H&E images (left), inferred growing-follicle axis positions using our common coordinate framework approach (middle), and cell type annotations for fibroblasts, theca and granulosa cells (right) for the two pediatric sections shown in panel “a” (AU14, 3 y/o; AU20, 15 y/o), each containing a growing antral follicle. Scale bar = 200µm. **c.** Magnified H&E image (left), inferred atretic-follicle axis positions using our common coordinate framework approach (middle), and cell type annotations for fibroblasts, theca and granulosa cells (right) for the two pediatric sections shown in panel “a” (AU14, 3 y/o; AU20, 15 y/o), each containing a growing antral follicle. Scale bar = 200µm. **d.** Smoothed splines of Xenium-measured expression (log-transformed) of theca markers. **e.** Dotplot showing the log-transformed, min-max normalised scRNAseq expression of selected NOTCH signalling genes (x-axis) in prenatal and pediatric+adult theca cells assigned to growing or atretic follicles (y-axis). **f-g.** Smoothed splines of Xenium-measured expression (log-transformed) of selected NOTCH signalling genes (“f”) and selected genes with different spatial dynamics between the growing and the atretic follicular axes (“g”). **h.** Boxplots showing the iss-patcher imputed scRNA-seq expression (log-transformed) of differentially expressed genes (y-axis; one-sided t-test; FDR < 0.01) in theca cells classified according to their predicted proximity (< 20 um) to growing or atretic follicles (x-axis).

Integration of Xenium spatial profiles with dissociated single-cell data allows us to resolve over 70 ovarian cell states across lifespan (**Figure 1e, Supplementary Figure 1c)**, each defined by distinctive marker expression and regional localisation (**Supplementary Figure 1d,2c,3d,4e**). We identify several under-explored somatic lineages, including OSE (*LHX2+/UPK3B+*) and glial populations (*PLP1*+), which have been underrepresented in previous datasets. We also identify novel ovarian stromal fibroblast populations, including adventitial C3+ fibroblasts “*Fib C3+*”. Within follicles, we recover oocytes (*ZP3+/GDF9+/DDX4+*) mostly likely from primordial follicles, and the three described theca populations^19^: perifollicular theca interna (*THBD+/PIEZO2+*) in direct contact with granulosa cells, structural theca externa (*PTCH1+/ACTA2^high^*), and androgenic/pre-androgenic theca interna (*CYP17A1+/ANPEP+*). We also identify novel granulosa cells spanning the full developmental continuum, from primordial (*RDH10+/FOXL2+/AMH-*) to preantral/early antral AMH+ granulosa subtypes (*RDH10-/FOXL2+/AMH+*). Overall, our atlas resolves all major lineages in fetal second-trimester and from birth through paediatric-adult stages, including novel paediatric-adult cell states. We provide a detailed description of each population’s spatial location and defining markers in **Supplementary Note 3**.

To contextualise our new large-scale atlas against existing resources, we reanalysed and harmonised cell annotation across published single-cell transcriptomic datasets of adult human ovaries (**Supplementary Figure 5a-c**). Leveraging the increased cellular and regional representation in our dataset, we were able to refine annotations and identify previously unresolved cell states in earlier studies, particularly within granulosa and fibroblast lineages. We find that granulosa cells associated with early follicle development (from primordial to preantral stages), fibroblasts from the outer cortex, and OSE are markedly underrepresented in the existing datasets (**Supplementary Figure 5b**). These discrepancies likely reflect differences in donor age, extent of outer cortex capture, and cell type recovery during tissue digestion, underscoring the importance of our dataset enriched in the outer cortex of pediatric donors for uncovering previously inaccessible biology. In addition, several datasets exhibit increased stress-associated signatures, suggestive of processing-related biases (**Supplementary Figure 5d**). Collectively, our atlas establishes a unified reference for the Human Cell Atlas to map existing and future ovarian single-cell studies.

### Postnatal OSE acquires immunomodulatory potential

Our multimodal atlas identifies the prenatal and postnatal transcriptomes of the OSE (**Figure 1e,h-i; Supplementary Figure 6a**), the fragile mesothelial monolayer encapsulating the ovary and historically difficult to capture in single-cell studies (**Supplementary Figure 6b**). In both prenatal and postnatal samples, OSE cells express mesothelial markers (*UPK3B, LRNN4*), gonadal specification factors (*WT1, GATA4*) and the transcription factor *LHX2* (**Figure 1j; Supplementary Figure 6c**), which is upregulated during human OSE specification around 8 PCW^12^.

Differential expression analysis reveals a marked shift in OSE identity between prenatal and pediatric-adult samples (**Figure 1j; Supplementary Table 3**). Fetal OSE expresses stem-cell associated (*HOPX, SOX11*) and second wave pre-granulosa (*TOX3*) transcription factors, all of which are silenced in the postnatal samples profiled (≥2 y/o). This transcriptional profile is consistent with the developmental role of prenatal OSE in ovarian organogenesis and its progenitor role to the second-wave of granulosa cells that establish the primordial follicle pool^12^. In contrast, pediatric-adult OSE upregulates *ADIRF*, *TMOD1*, and *TNXB*, genes implicated in inhibiting cell growth and migration, and induces epithelial markers (*KRT7*; desmosomal *PERP* and *DSC3*) and extracellular matrix components (*NTN4, LAMA4, COL8A1, COL12A1*). Postnatal OSE also downregulates OSE-identity transcription factors *LHX2* and *GATA4*, yet maintains basal expression. Together, these features are indicative of a more stable epithelial identity, consistent with the barrier function of post-pubertal OSE that undergoes repeated rupture and regeneration during ovulatory cycles.

Postnatal OSE also exhibits a robust immunomodulatory programme. From our youngest pediatric (2 y/o) onward, OSE cells upregulate complement components (*C3, CFB, CFH*), inflammatory cytokines (*IL18*, which recruits innate immune cells; *IL16,* acting on CD4+ T cells), and interferon-associated genes (*IRF9, IRF7, STAT1, MVP, GABARAPL1, MNDA, TRIM22, TRIM38;* **Figure 1j**). OSE also expresses immunoregulatory molecules (*HLA-F*, interacting with inhibitory receptors on innate immune cells; *VSIR*, an immune checkpoint regulator) and the antimicrobial peptide *SLPI* (**Figure 1j**). Together, our data suggest that postnatal OSE establishes an immunoregulatory niche capable of modulating local immune activity. This is consistent with emerging evidence that structural cells participate in immune responses^42,43^, and may thereby contribute to protection of the primordial follicle reserve during repeated ovulation-associated surface injury^44,45^.

### Fine map of early granulosa cells identifies novel states

Follicles are the functional reproductive and endocrine units of the ovaries, each comprising an oocyte surrounded by granulosa cells. These cells undergo marked phenotypic transitions as follicles progress from quiescent squamous granulosa in primordial follicles, established prenatally ∼17 PCW, to preantral, antral, ovulatory, and their corresponding atretic stages^2^. Prior single-cell transcriptomic studies have typically treated granulosa cells as a single broad component^18,20,21^ or focused only on more advanced follicles^17,46,47^. Here we resolve, for the first time, the cortical granulosa cell states spanning early follicle development in humans (**Figure 2a; Supplementary Figure 7a-i**).

We identify seven granulosa subtypes whose differentiation order was inferred using trajectory analysis of scRNA-seq data (**Figure 2b**) and independently validated using spatial mapping of subtype specific markers in Xenium and Visium HD data onto follicles at distinct morphological stages (**Figure 2c-e**). These included: 1) AMH^-^ squamous granulosa in primordial follicles (*RDH10^+^/WNT6^+^/AMH^-^*); 2) transitioning granulosa in activating and primary follicles, co-expressing *AMH* along squamous markers (*RDH10+/WNT6+*) and *PTPRZ1*, a modulator of PI3K/AKT/mTOR^48^, the proposed pathway that governs primordial quiescence^49^; 3) AMH+ early growing granulosa cells upregulating *COL2A1*; 4) AMH+ multilayered granulosa upregulating aromatase *CYP19A1* and *IHH* genes in small preantral follicles; 5) AMH+ steroidogenic granulosa cells upregulating *HSD3B2* (*CYP19A1+*/*IHH+/HSD3B2*+) mapping to preantral or small antral follicles, mostly cycling (*MKI67+)*; 6) *AMH+* steroidogenic granulosa cells upregulating *EPPIN* and enriched in the cumulus (*EPPIN+/WFDC6+*) and 7) *AMH+* steroidogenic upregulating *GDF3* and enriched in the antral follicle wall (*GDF3+*) (**Figure 2a,c-e; Supplementary Figure 7i; Supplementary Table 4a-b**).

In addition, we detect two distinct atretic granulosa populations characterised by broad downregulation of housekeeping genes, one resembling *AMH-* and the other *AMH+* granulosa population. We also noted a subset of *AMH+* granulosa cells downregulating Hedgehog signalling ligands *DHH* and *IHH* that we term AMH+ Hedgdog-low granulosa (**Supplementary Figure 7j**). Comparison with prenatal ovaries shows that the more mature granulosa cells observed during the second trimester ovaries, previously described as second-wave OSE-derived “primordial granulosa”^12^, align transcriptomically with the AMH- granulosa population observed in postnatal samples (**Supplementary Figure 7k**), indicating that these populations represent a conserved developmental state across prenatal and postnatal life.

### Retinoic acid signalling switches at the primordial granulosa transition

Having captured cortical granulosa cell states across early follicle development using single-cell transcriptomics (**Supplementary Figure 7a**) and chromatin accessibility profiling (**Supplementary Figure 7l**), we next examined their regulatory trajectories in detail. As granulosa progresses from squamous to transitioning states, we observe widespread chromatin closure, with regions that close at the initial step remaining closed thereafter (**Figure 2f**). In contrast, opened chromatin regions are highly state-specific and increase in numbers as differentiation proceeds. These newly accessible regions include promoters or transcription starting sites (TSS) of hallmark folliculogenesis genes, such as AMH, FSHR, GSTA1 or IHH, whose accessibility increases in parallel with transcriptional activation (**Figure 2g**; **Supplementary Figure 7m; Supplementary Table 4c**). Regions undergoing closure are enriched for transcription factor motifs associated with second-wave granulosa specification including LHX9, LHX2, NR1H4 (FXR) (**Supplementary Figure 7n; Supplementary Table 4d**).

During the squamous-to-transitioning step, granulosa cells lose accessibility at Nuclear Factor I (NFI) binding sites and downregulate NFIA/NFIB expression (**Figure 2h; Supplementary Figure 7n**). Given the pioneer-like enhancer regulatory roles of NFI in stem-cell contexts^50,51^, the observed silencing of NFI-dependent enhancers suggests a loss of less differentiated characteristics during granulosa transition. Closing chromatin regions are also enriched for DNA methyltransferase (DNMT) binding sites (**Supplementary Figure 7n**) and coincide with increased expression of de novo DNA methyltransferase DNMT3A (**Figure 2h**). Upon reaching the AMH+ multilayered state, granulosa upregulates the DNA methylation maintenance machinery (*DNMT1*, *UHRF1, PCNA;* **Figure 2h**), consistent with replication-coupled propagation of methylation marks. Together, these observations support a unidirectional epigenetic commitment during early granulosa activation, driven by NFI enhancer silencing and DNA methylation-mediated chromatin remodelling.

The transition from squamous to AMH+ granulosa is accompanied by repression of retinoic acid (RA) signaling, a regulatory switch that has not previously been described. Transitioning granulosa cells reduced chromatin accessibility at NR1H4 and RA receptor (RAR) DNA-binding motifs, both of which form heterodimers with RXR (**Supplementary Figure 7n**) together with downregulation of RA-biosynthetic genes (*ALDH1A1/2 and RDH10,* whose promoter also closes; **Supplementary Figure 7m**). Concurrently, these cells upregulate RA-catabolism (*CRABP1*) and the RAR antagonist *PRAME*, whose promoter becomes accessible (**Figure 2i, Supplementary Figure 7m**). In parallel, transitioning granulosa cells activate PPARγ lipid metabolic programmes (*PPARG, FABP5)* and increases expression and chromatin accessibility of *NR5A2* (also known as *LRH-1*), consistent with increased cholesterol uptake in preparation for steroid hormone biosynthesis (**Figure 2i; Supplementary Figure 7m-n**). Accordingly, cholesterol transporters (*ABCA8, ABCA5*) are upregulated, while cholesterol oxidation via *CYP27A1* is reduced (**Figure 2i**). Together, these changes indicate a shift in RXR partner usage from RAR-RXRs and NR1H4-RXRs (FXR-RXR) complexes toward PPARγ-RXRs and NR5A2 driven programmes, redirecting cellular metabolism away from RA signalling and toward lipid utilisation and early steroidogenic priming^52–54^.

Concomitant with this metabolic reprogramming, we observe attenuation of WNT signalling as granulosa enter the AMH+ early state, including downregulation of *WNT6*, *WNT2B* and the canonical target *AXIN2* (**Figure 2i**). In addition, early granulosa cells transiently upregulate components of the NOTCH pathway, including receptors (*NOTCH1/3/4*) and downstream effectors (*HES4, HEY1, HEYL*), followed by decreased expression and closed chromatin accessibility as cells advanced into AMH+ early and multilayered states (**Figure 2i; Supplementary Figure 7m-n**). Our findings are consistent with an antagonistic relationship between PPARγ-RXRs signalling and both WNT and NOTCH pathways^55,56^.

Collectively, these findings reveal that early granulosa differentiation is accompanied by extensive epigenetic and transcriptional rewiring, with suppression of RA signalling emerging as a central regulatory event coordinated with metabolic and signalling pathways modulation, which may guide the transition from primordial to early growing follicle states.

### Stromal fibroblast gradients stratify the ovarian cortex

Follicles are embedded within stromal fibroblasts, the predominant cell type of the human ovary, yet the organisation, heterogeneity and signalling functions of these fibroblasts remain poorly defined. Spatial profiling of pediatric and adult samples using Xenium (**Figure 3a**) uncover a higher-order cortical architecture composed of multiple fibroblast strata, revealing a complexity that extends beyond the canonical cortex-medulla division (**Figure 3b-c; Supplementary Figure 8a-b)**. These strata are less apparent in scRNA-seq data^57^, likely due to dissociation-induced effects that mask gradual transcriptional transitions between fibroblast states (**Supplementary Figure 8c**). Consequently, fibroblast fine annotations were derived exclusively from spatial data.

To capture cortical microanatomy across donors and standardise cortical depth despite differences in ovarian size, sampling region, and section orientation, we mapped stromal fibroblasts from each tissue section (ages 2 to 28 y/o) into a Common Coordinate Framework^58,59^. The resulting “*Cortico-Medullary (CM) axis”* reflects the distance from the outermost cortex toward the medulla (**Figure 3b-c; Supplementary Note 4**). Unlike histologically-based axes^58,60^, which cannot be applied here due to the frequent loss of OSE (the outer cortex histological landmark), our approach aligns conserved fibroblast landmarks defined by shared Xenium expression patterns across donors (**Figure 3b-c**). We projected this CM axis onto the single-cell dataset (**Supplementary Figure 8d**), enabling annotations of fibroblast states that cannot be resolved by clustering scRNA-seq data alone^57^, and enabling quantification of gene expression changes along cortex-to-medulla trajectories using TradeSeq^61^ (**Supplementary Figure 8e; Supplementary Table 5a**).

This analysis reveals a continuous cortico-medullary gradient defined by distinct fibroblast strata that compartmentalises the ovarian cortex into niches with unique paracrine and biomechanical identities (**Figure 3d-e**). Superimposed on this gradient, we identify local deviations surrounding preantral and more differentiated follicles, including atretic follicles (**Figure 3a-b; Supplementary Figure 8g-h**), which we term perifollicular fibroblasts, reflecting folliculogenesis-driven remodelling of the stroma to support follicle growth or atresia.

Along the CM axis, we resolve successive stromal fibroblast strata: OSE-adjacent, WNT4+ secretory, primordial follicle border, inner cortical and medullary fibroblasts, as well as a smooth-muscle-like population (**Figure 3e-f; Supplementary Figure 8e**). OSE-adjacent and WNT4+ secretory fibroblasts above the primordial follicles (**Supplementary Figure 8e-f**) express WNT pathway components (*WNT4, TRABD2A, SFRP4, TSPAN8*), FGF (*FGF10*), IGF modulators (*IGFBP6/IGFBP3*), antagonists of TFG-b superfamily (*FSTL3* and *GREM*, antagonist of activins and BMPs, respectively), the chemoattractant *SEMA3C*, the secretory granule protein *CHGB*, ECM constituents (*LAMA2, HMCN1, CILP2, COL8A1, ATRNL1*), matrix remodeler (*MMP11*), pericellular protease inhibitors (*SERPINA5, LXN, TFPI2*), receptors (*PTGER2, BMPR1B, GRM8, TMEM100*) and retinoic acid response genes (*RARRES1, DHRS3*; **Figure 3d; Supplementary Figure 8e,g-h; Supplementary Table 5a**). The OSE-adjacent fibroblasts share all of these markers with WNT4+ secretory fibroblasts, but also upregulate the collagen *COL10A1* and *HOPX* (**Figure 3d; Supplementary Figure 8g-h**). Spatially, WNT4⁺ fibroblasts coincide with the dense, compact outer cortical region described in histological studies^62^. Re-analysis of published adult ovarian scRNA-seq datasets reveals this population to be present at very low abundance compared to our study (**Supplementary Figure 8i-j**), likely due to the absence of pediatric samples and the underrepresentation of the outer cortex, which may explain why these cells went previously unnoticed in their corresponding studies.

At the inner end of the WNT4+ region, primordial follicle border fibroblasts flank primordial follicles (**Figure 3e-f; Supplementary Figure 8e-f**). These cells downregulate the WNT4+ secretory programme and upregulate the *KIT* receptor while reducing *LUM* expression, indicating a shift in ECM composition and signalling properties (**Figure 3d**). Moving inward, inner cortex and medullary fibroblasts express the WNT antagonist (*SFRP1),* the TGFb growth factor modulator (*LTBP1)* and genes associated with a perivascular-like niche, including the transcription factor *EBF1* and neurovascular support genes (*GFRA1, CXCL12, IGF1/2, PROK1* and *SLC18A2),* reflecting likely roles in supporting vascular, lymphatic and neural structures of the inner cortex and medulla (**Figure 3d; Supplementary Figure 8g-h**). These fibroblasts also upregulate the cholesterol transporter *STAR* which mediates the rate-limiting step of steroidogenesis, consistent with a more pro-steroidogenic niche (**Figure 3d; Supplementary Figure 8g-h**).

At perifollicular niches, stromal fibroblasts diverge from baseline cortico-medullary programmes (**Figure 3a-b; Supplementary Figure 8g-h)**. We identify two major perifollicular subtypes: one resembling the WNT4+ secretory fibroblasts and another resembling inner cortical fibroblasts with a neurovascular signature. Both show theca externa-like features including upregulation of collagen IV (*COL4A1*), laminins (*LAMA4* and *LAMB1*), Biglycan (*BGN*) and the Hedgehog pathway receptor (*PCTH2*), likely in response to DHH/IHH ligands from AMH+ granulosa cells in the near follicles (**Supplementary Figure 8g-h**). However, the absence of theca-specific markers (*PTCH1+*) indicates that perifollicular fibroblasts (*CLDN1-*) and theca externa (*CLDN1+*) represent distinct cell states (**Supplementary Figure 8g-h**). The presence of these perifollicular populations accounts for stromal fibroblast heterogeneity within and between cortical samples; for example, in tissue rich in large atretic follicle remnants, inner cortical fibroblasts predominantly adopt a perifollicular identity (**Figure 3a, left sample**). In our cohort, perifollicular fibroblasts are more abundant in ovaries ≤3y/o, consistent with previously reported higher follicular activity after birth^63,64^ **(Figure 3a left; Supplementary Figure 9a-b)**.

Finally, we identify a previously uncharacterised fibroblast subtype (Fib C3+) located around vasculature and, in some donors, immediately beneath the OSE along the *tunica albuginea* (**Figure 3g; Supplementary Figure 8k**). This population exhibits a transcriptomic profile that is different from stromal fibroblasts, expressing adventitial fibroblast markers (*C3, PI16, CD34, DPT*) together with gonadal transcription factors (*GATA4, WT1, ARX, LHX9*), consistent with a specialised adventitial-like fibroblast of ovarian origin (**Figure 3h**).

### Spatial signals in the primordial follicle reserve niche

Spatial profiling reveals that squamous granulosa in primordial follicles reside adjacent to the WNT4+ secretory fibroblasts in both pediatric and adult outer cortex (**Figure 3e-f; Supplementary Figure 8e-f**), placing them within diffusion range of fibroblast-derived signals. Although the cortical stroma has been proposed to regulate primordial follicle fate^65^, the cellular identity and molecular composition of this niche have remained poorly defined^24,25^. The conserved proximity of squamous granulosa cells and WNT4+ fibroblasts across donors suggests that these fibroblasts may influence primordial follicle quiescence and activation through paracrine signals.

Systematic ligand-receptor analysis predicts a previously unrecognised paracrine network involving retinoic acid, IGF, and WNT signalling between squamous granulosa and neighbouring fibroblast (**Figure 3i**). WNT4+ fibroblasts upregulate the RA-responsive gene *RARRES1*, consistent with exposure to an all-trans RA-rich environment generated by quiescent primordial follicles, which express the RA biosynthetic enzymes *RDH10, ALDH1A1 and ALDH1A2*. Both squamous granulosa and WNT4+ fibroblast express the IGF2 inhibitor, *IGFBP6,* at high levels, suggesting that primordial follicles reside within an IGF2-buffered niche that may mitigate *IGF2* activity from inner cortical fibroblasts and vasculature (**Figure 3d,i; Supplementary Figure 8l**). Primordial border fibroblasts, which embed primordial follicles, show finely tuned WNT signaling: they still express *WNT4*, although at lower levels, alongside antagonists (*SFRP4, TRABD2A, TSPAN8*), while squamous granulosa express *WNT6, WNT2B* and downstream effectors (*AXIN2, ZNRF3*), indicative of low but sustained canonical WNT activity in the primordial follicle across the lifespan within primordial granulosa (**Figure 3i**).

Despite their shared proximity to primordial follicles, the postnatal WNT4+ population and the fetal ovarian fibroblasts are assigned as two distinct clusters (**Figure 1e-f**), suggesting a significant developmental shift in their transcriptomic profiles. To resolve these differences, we compared pediatric WNT4⁺ secretory fibroblasts (1-15 years, when there is a significant pool of quiescent primordial follicles) with fibroblasts adjacent to primordial follicles in second-trimester fetal ovaries (17-21 PCW), when the primordial follicle pool is being established (**Figure 1e**)^12^. Fetal second-trimester stromal fibroblasts express WNT4, but lack postnatal WNT modulators (*TRABD2A, TSPAN8),* as well as additional postnatal markers such as the IGF-2 modulator *IGFBP6* and *COL10A1*. Instead, fetal fibroblasts upregulate the RA-binding protein *CRABP2*, consistent with increased RA intake, and express the NOTCH antagonist *DLK1* (**Figure 3i**) which in postnatal tissue is restricted to theca. These differences indicate a developmental shift from a NOTCH-inhibited, WNT-active niche in the prenatal ovary to a NOTCH-tolerant, WNT-buffered niche postnatally. Given that WNT and NOTCH are active in squamous and transitioning granulosa, but downregulated upon entry into AMH+ stages (**Figure 3i**), these observations suggest that balanced NOTCH/WNT signals may modulate primordial follicle quiescence.

Together, our analyses provide a detailed molecular map of the fetal and postnatal niches supporting the human primordial follicle reserve and uncover previously unrecognised regulatory interactions that shape granulosa differentiation and follicle development.

### Differences in perifollicular niches of growing and late atretic follicles

Following follicle activation, AMH⁺ granulosa cells proliferate to form multilayered preantral and antral follicles. As follicles grow, AMH⁺ granulosa cells extend into deeper inner-cortex fibroblast strata, where they encounter distinct microenvironments (**Figure 3e-f**) and emerging supporting cell types, such as theca. Most growing follicles ultimately undergo atresia; large follicles leave behind remnant granulosa and theca cells that persist within the inner cortical stroma before eventual clearance^66^. Our previous results showed that early antral growing and atretic remnant follicles are both surrounded by “perifollicular fibroblasts”, which are distinct from the baseline cortico-medullary programme (**Figure 3a-b**), consistent with local stromal remodelling associated with follicle growth and degeneration.

To systematically compare local stromal environments during follicle growth and atresia, we constructed a shared spatial coordinate framework that maps fibroblast and theca cells relative to their nearest AMH+ granulosa cells in either growing early antral follicles or atretic remnants (**Figure 4a**). Using the granulosa-theca boundary as a spatial reference, we define a “*Growing Follicular axis*” and an “*Atretic Remnant Follicular axis*”, this last one around follicles in later stages of atresia (**Figure 4b-c; Supplementary Note 4; Supplementary Table 5b-c**). Both axes recover the expected spatial sequence of theca states, perifollicular interna, closest to the follicle, followed by androgenic interna and externa, and then perifollicular fibroblasts (**Figure 4b-c right**). Consistently, expression of canonical theca marker genes correlates with distance from the follicular granulosa layer (**Figure 4d**). A full list of spatially variable and differentially expressed genes is provided in **Supplementary Tables 5b-d**.

Despite these shared organisational features, we identify compartmentalised NOTCH signalling across theca subsets. NOTCH receptors and downstream effectors (*NOTCH1/3, HEYL, HES1*) are enriched in perifollicular theca interna and externa, whereas androgenic theca instead upregulate the ligands *JAG1* and the non-canonical modulator *DLK1*, with *DLK1* further increased in atretic theca (**Figure 4e; Supplementary Tables 5d**). NOTCH signalling has previously been described in the *corpus luteum* of other mammals^67–71^, but not in theca cells during early follicle differentiation. Altogether, our integrated analysis across lineages and developmental stages reveals a highly dynamic NOTCH activity spanning folliculogenesis: receptor activity is elevated in squamous granulosa cells of fetal primordial follicles, downregulated as granulosa enter the early AMH⁺ state in primary and preantral follicles (**Figure 2g**), and re-established in emerging theca (**Figure 4e**). *DLK1*, characteristic of fetal ovarian fibroblasts but not postnatal fibroblasts (**Figure 3i**), reappears in androgenic and atretic theca (**Figure 4e**).

Beyond NOTCH signalling, growing and late atretic follicles display distinct molecular niche signatures. Along the *Growing Follicular axis,* but not in the *Atretic Remnant Follicular axis*, theca cells adjacent to AMH⁺ granulosa upregulate cell-cycle genes (*UBE2C*) and the Hedgehog pathway ligand *GLI1* (**Figure 4g-h**), consistent with theca proliferation driven by paracrine DHH/IHH signalling from AMH⁺ granulosa cells, the primary drivers of theca specification and differentiation^72,73^. Theca interna cells in the *Growing Follicular axis* also uniquely upregulate *VEGFA*, indicative of pro-angiogenic niche, together with genes associated with stromal blood-barrier cells, including ECM regulators (*CCN1/2*), tight-junction components (*CLDN1*)^74,75^ and immune-modulatory factors *LGALS1* and *WFDC1*^76,77^ (**Figure 4g-h; Supplementary Table 5d**). To our knowledge, *WFDC1* and *CLDN1* have not previously been described in theca cells, yet are among the most distinctive markers in our cohort.

In contrast, theca cells along the *Atretic Remnant Follicular axis* show reduced expression of proliferative, angiogenic, and barrier-associated genes, consistent with decreased proliferation and breakdown of the blood-follicle-barrier described in late atresia^66^ (**Figure 4g-h**). Instead, atretic theca upregulate lipid and cholesterol metabolism genes associated with steroidogenic activity (*PDK4*, *STAR*, *CYP11A1, DHCR24* and *PSAP),* as well as the complement factor *C7,* consistent with prior reports of complement gene expression within follicular compartments^17^. Atretic theca cells also upregulate monocyte recruitment factors like *APOE* and the chemokine *PROK1,* which had been implicated in follicular atresia in bovines^7866^. Together, these findings indicate that atretic theca cells acquire a dual metabolic and immune-recruiting phenotype, promoting monocyte infiltration and differentiation into macrophages, likely to aid clearance of apoptotic cells and active tissue remodeling.

### Lipid-associated macrophages in the ovary

Tissue-resident macrophages are functionally specialised to their local microenvironments through distinct transcriptional programmes^79,80^. We jointly analyse the fetal and pediatric ovaries, and identify three tissue-resident macrophages **(Figure 5a, Supplementary Figure 10a, Supplementary Table 6a)**. These populations correspond to macrophage subsets previously described in adult ovary and assigned based on the expression of canonical markers^81^ (**Supplementary Note 3.6**), and our cross-lifespan analysis provides insights into their molecular identity. We adjusted their name to reflect those features.

**Figure 5.**
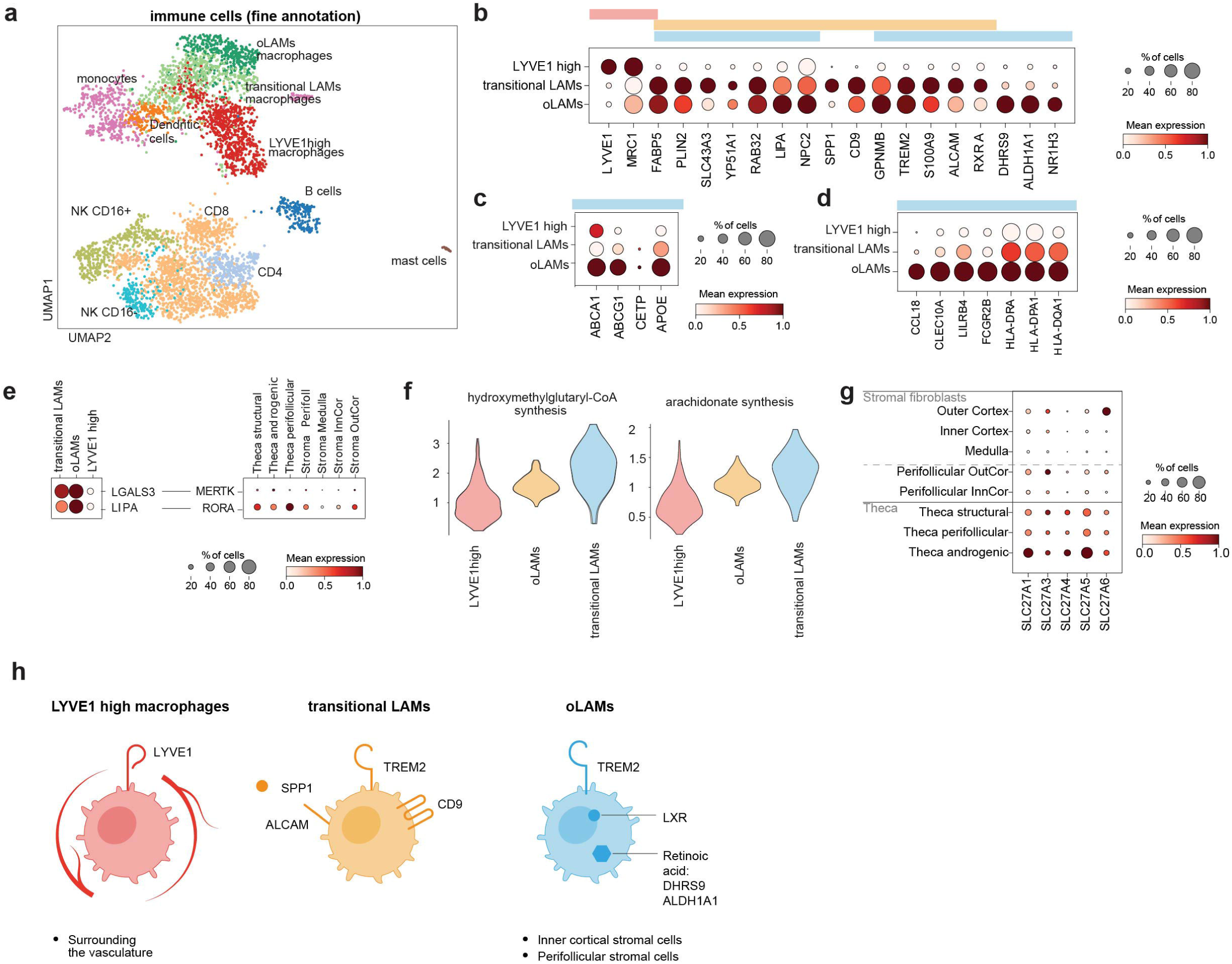
Ovarian macrophages. **a.** Batch-corrected Uniform Manifold Approximation and Projection (UMAP) embedding of the scRNA-seq immune cells from the pediatric-adult ovaries (n = 4,819). **b.** Dotplot showing the log-transformed, min-max normalised expression of macrophage marker genes (x-axis) across the three ovarian macrophage subtypes (y-axis). **c.** Dotplot showing log-transformed, min-max normalised expression of LXR target cholesterol efflux transporters and lipid transport genes (x-axis) across the three ovarian macrophage subtypes (y-axis). **d.** Dotplot showing log-transformed, min-max normalised expression of anti-inflammatory genes and antigen-presentation genes (x-axis) across the three ovarian macrophage subtypes (y-axis)**. e.** Dotplot showing the log-transformed, min-max normalised expression of selected macrophage genes and stromal- and thecal-associated signalling genes (x-axis) across macrophage subsets (y-axis). **f.** Violin plots showing the distribution of scCellFie metabolic scores for key lipid-associated pathways across ovarian macrophage subsets. **g.** Dotplot showing the log-transformed, min-max normalised expression of fatty-acid transporters of the SLC27A family genes (x-axis) across stromal fibroblast and theca cells (y-axis). **h.** Diagram of macrophage subtypes.

The first population, termed *LYVE1* high macrophages, is defined by high expression levels of *LYVE1+* and the canonical tissue-resident marker *MRC1* **(Figure 5b)**. These macrophages resemble interstitial, vessel-associated macrophages described in multiple murine tissues, where they support tissue repair and homeostasis^82^. The second population, termed transitional lipid-associated macrophages (LAMs), is defined by upregulation of monocyte markers (*S100A9*) as well as upregulation of lipid-metabolism genes (*FABP5*, *PLIN2*, *SLC43A3*, *CYP51A1, RAB32*), the lysosomal lipase *LIPA*, and the cholesterol transporter *NPC2*. These macrophages also express canonical markers of LAMs^83^ (*TREM2*, *CD9*, *GPNMB, SPP1)* **(Figure 5b, Supplementary Table 6a)**. The third population, termed oLAMs, share the lipid-metabolic genes and LAM-associated transcriptional programme of transitional LAMs (apart from *SPP1*) but is distinguished by upregulation of retinoic acid synthesis genes (*DHRS9, ALDH1A1*) and the lipid-activated transcription factor *LXRa* (*NR1H3*). LXRa heterodimerises with RXR, which is activated by 9-cis RA, enabling transcriptional regulation control of lipid-metabolic programmes **(Figure 5b).** Consistently, oLAMs-LXR upregulate established LXR target genes, including the cholesterol efflux transporters (*ABCA1*, *ABCG1)*, *APOE*, and *CETP* **(Figure 5c)**. oLAMs also express anti-inflammatory (*CCL18*, *CLEC10A*, *LILRB4*, *FCGR2B)* and antigen-presentation genes (*HLA-DRA, HLA-DPA1, HLA-DQA1*; **Figure 5d)**.

Spatial analysis using k-nearest-neighbour enrichment reveals distinct niche associations (**Supplementary Figure 10b, Supplementary Table 6b)**. oLAMs were enriched near perifollicular fibroblasts and inner cortical stromal cells, suggesting these are the macrophages that infiltrate atretic follicles^66^. Ligand-receptor analysis identifies multiple potential interaction modules between oLAMs, theca cells and fibroblasts (**Supplementary Figure 10c, Supplementary Table 6c).** oLAMs upregulate *LGALS3*, an opsonising ligand predicted to activate MERTK on neighbouring theca cells, potentially facilitating macrophage-mediated tissue-remodelling during follicular atresia **(Figure 5e)**.

Consistent with their transcriptional profile, metabolic inference using scCellFie^84^ predicts increased activity of hydroxymethylglutaryl-CoA synthesis, a precursor to cholesterol biosynthesis, as well as arachidonate synthesis in oLAMs **(Figure 5f, Supplementary Table 6d)**. These findings suggest that oLAMs generate lipid metabolites that may be transferred to neighbouring cells. Supporting this model, theca cells and fibroblasts express *RORA*, a nuclear receptor that promotes steroidogenesis, and theca cells express fatty acid transporters of the *SLC27A* family, consistent with potential uptake of macrophage-derived lipids **(Figure 5e,g).**

Together, these data identify ovarian lipid-associated macrophages as spatially organised, metabolically specialised immune cells that support extensive follicular remodelling by phagocytosing dying cells while maintaining an immunoregulatory environment that limits inflammation (**Figure 5h)**.

## Discussion

Despite the pivotal role of the ovary in women’s health and fertility, surprisingly little is known about the cellular mechanisms controlling primordial follicle quiescence and growth in humans. Here, we present a high-resolution, multimodal single-cell and spatial map of the human ovarian cortex from pediatric and adult donors, integrated with publicly available prenatal datasets^12^. This resource resolves the cellular events along the establishment and activation of the primordial follicle reserve across the human lifespan. The resulting ovarian atlas constitutes a fundamental reference for human ovarian biology by (1) profiling the largest number of cells and donors to date, including the pediatric window; (2) providing harmonised, high-resolution cell-state annotations across studies; (3) capturing large numbers of primordial and transitional follicles that enable inference of mechanisms underlying follicle activation; (4) integrating multimodal measurements, including chromatin accessibility, to resolve the regulatory programmes underpinning cellular identity; and (5) providing spatial and temporal contexts essential for understanding the role of the niche environment in follicle development.

This study represents the first iteration of the ovarian atlas within the Human Cell Atlas (HCA) initiative^85^ that aims to generate comprehensive reference maps of the human body. We mitigate batch effects and clinical confounders by using a large cohort spanning diverse non-ovarian clinical backgrounds and relying on spatial mapping to overcome any potential artefacts associated with enzymatic dissociation that can influence follicular activation^86^. The atlas enables rapid annotation and harmonisation of ovarian transcriptomic studies, as shown by transferring labels to previous ovarian cellular maps; and identification of potential off-target effects of fertility-compromising therapies^87^. We define clear transcriptomic differences between pre- and postnatal ovaries, including the establishment of OSE as an immunomodulatory protective barrier, and shifts in cell-type proportions across postnatal stages, such as within perifollicular fibroblasts. Future iterations of the ovarian atlas will expand cell population coverage to complete the full trajectory of follicle maturation and luteinisation (like for pre-ovulatory, ovulatory and post-ovulatory structures such as the corpus luteum and the corpus albicans), and will pair single-cell profiling with histological assessment of individual follicles to refine granulosa cell states according to follicle health status, which will be required to fully define the exquisite regulation of this organ.

Beyond serving as a reference resource, this atlas provides new biological insights into human ovarian physiology. Existing ovarian single-cell atlases have largely focused on adult tissues^17–21^, where primordial follicles are rare and frequently underrepresented. By including pediatric samples enriched for the primordial follicle pool and specifically targeting ovarian cortex, we resolve, for the first time, a continuum of cortical granulosa cell states underlying primordial follicle activation and early follicle development. Our analysis, including paired open chromatin and transcriptomics measurements, reveals coordinated regulatory and signalling changes accompanying the transition from follicle dormancy to growth within the native cortical niche. Among these, RA signalling emerges as the most extensively remodelled pathway during primordial activation, implicating it in early follicle differentiation. This observation is consistent with clinical evidence linking systemic retinoid exposure (with isotretinoin) to reduced ovarian reserve markers (serum AMH, antral follicle count and estradiol) despite preserved gonadotropin levels^88^. Beyond epigenetic and transcriptional modulation of RA signalling, we observe coordinated changes in WNT and NOTCH pathway activity, although future mechanistic studies will be required to establish causal relationships among these signalling axes.

While decades of research have been follicle-centric, focusing primarily on oocytes and granulosa cells, our study underscores the importance of the microenvironment by revealing the highly organised cellular architecture of cortical stromal fibroblasts and the presence of specialised follicular macrophages. Our single-cell-resolved spatial approach identifies that cortical fibroblasts are not a homogeneous population but are organised into distinct layered structures that generate morphogen and signaling gradients. This architecture was previously unresolvable by scRNA-seq alone^18,20^ and required integration of spatial transcriptomics with histopathological image registration to establish a Common Coordinate Framework (CCF)^57–59^ across pediatric and adult donors. This donor-consistent spatial alignment enabled precise localisation of WNT4+ secretory fibroblasts to the dense, rigid stroma of the outer cortex where the primordial follicle reserve resides. Moreover, it allowed systematic identification of a previously unrecognised paracrine network regulating WNT, IGF, NOTCH and RA signaling pathways, which we predict to be relevant for granulosa activation and subsequent follicular development. These niche-derived signals highlight candidate upstream pathways that may converge on the PI3K/AKT/mTOR axis, a post-translationally regulated axis linked to primordial quiescence and activation that remains incompletely characterised in humans^86,89–^91.

Our spatial framework further supports a view of the ovarian stroma as a dynamic scaffold that is continuously reshaped by follicle growth and atresia, moving beyond simplified models in which follicles merely migrate from the dense outer cortex to the vascularised medulla as they mature^6^. In our updated model, follicle-adjacent perifollicular fibroblasts display transcriptional profiles distinct from follicle-distant fibroblasts and spatially co-localise with theca and specialised immune populations. This dynamic remodelling implies a reciprocal relationship whereby growing and atretic follicles reconfigure the local architecture and signalling milieu, which may in turn influence neighbouring, less differentiated follicles, reminiscent of a domino effect. Variation in the local abundance of growing or atretic follicle niches contributes substantially to donor-to-donor sample heterogeneity, posing major challenges for comparative analysis based on ovarian biopsies. Spatially registered reference frameworks such as the one presented here are therefore essential for accurate sample alignment and identification of true biological signals.

Our data-driven analysis also assigns a central role to immune cells in regulating follicle homeostasis. We define oLAMs that localise surrounding the follicles and express lipid metabolism genes, immunoregulatory genes, and phagocytosis related genes, suggesting a role in clearing lipid-rich cellular debris generated during folliculogenesis while maintaining an immunoregulatory profile that dampens inflammation. oLAM expresses the machinery required for RA production and lipid response – with RXRa previously shown to be expressed by several tissue resident macrophages^92,93^ – in the ovary, this population may additionally contribute to local steroidogenic processes. More broadly, LAMs with transcriptomic profiles similar to oLAMs are found in other tissues, including adipose tissue^94^, and liver^95,96^, where they have been implicated in disease progression^95^. The ovary is affected by multiple gynecological disorders, such as endometriosis and ovarian cancer, in which RXR-regulated macrophages have been implicated in tumour progression^93^. We therefore speculate oLAMs may contribute both to pathological states and overall fertility by supporting proper follicle development.

Overall, this atlas has direct and far-reaching translational relevance for fertility preservation. For girls undergoing gonadotoxic treatments such as chemotherapy, cryopreservation of cortical strips is standard care, yet clinical outcomes remain variable and mechanistically poorly understood. By resolving the cellular, spatial and signalling architecture of the human ovarian cortex, our atlas provides a rational framework to refine cryopreservation and tissue-handling strategies, including approaches to suppress premature follicle activation and preserve niche integrity. More broadly, it lays the foundation for physiologically informed *ex vivo* culture systems and *in vitro* follicle maturation protocols, with the potential to substantially advance fertility preservation and reproductive medicine.

## Methods

### Patient samples

Ovarian cortical tissues from chemonaive pediatric donors were obtained via laparoscopy for the Oxford Cell and Tissue Biobank (OCTB), which stores cryopreserved ovarian cortex collected from paediatric patients undergoing fertility preservation prior to gonadotoxic therapy. The study (REC: 19/YH/0441) was approved by the Yorkshire & The Humber - Leeds East Research Ethics Committee for the Sanger Institute, and the study (REC: Ethics REC: 19/NW/0526) by the North West - Greater Manchester West Research Ethics Committee for the Williams team. Ovarian tissues from adult donors of reproductive age were obtained from the Newcastle upon Tyne Hospitals via either laparoscopic cortical biopsy or oopherectomy (whole ovaries, REC:16/NE/003). The study (REC:16/NE/0003) was approved by the North East - Newcastle & North Tyneside 1 Research Ethics Committee. In all instances, written, informed consent was provided by study participants before obtaining tissue samples and phenotypic data.

One whole ovary sample from an adult of reproductive age was obtained from deceased transplant organ donor within 1 h of circulatory arrest. The study (REC: 15/EE/0152) was approved by the East of England-Cambridge South Research Ethics Committee. Written and informed consent was provided by the donor families.

### Tissue processing

Pediatric ovarian cortical biopsies were collected via laparoscopy and subsequently dissected by the OCTB biobank into ∼1 mm cortical strips prior to cryopreservation. Adult ovarian samples were also collected via laparoscopy and placed directly into the transport medium without further processing. In both cases, samples allocated for research were kept and transported in HypoThermosol FRS solution (Sigma Aldrich, cat. no. H4416) at 4 °C. This approach preserved tissue integrity for downstream single-cell and spatial multi-omic profiling. Fresh samples were subjected to single-cell dissociation for scRNAseq, scATACseq and, when possible if sample size allowed, a small piece of fresh tissue was also embedded in OCT compound (ThermoFisher Scientific, cat. no. 23730571) inside a cryomold and rapidly frozen using an isopentane bath cooled on dry ice (−75 °C).

For some pediatric samples, inner cortex remnants from cutting the cortex into strips were also retained and transported in HypoThermosol FRS solution. Remnant pieces were examined by light microscopy, and individual whole visible follicles (0.2–2 mm in diameter) were manually isolated and processed for single-cell isolation.

### Single-cell isolation from fresh tissue

Tissue dissociation from fresh ovarian samples was conducted within 48hr of tissue retrieval. Upon arrival, both pediatric and adult fresh ovarian samples were washed with PBS and finely minced into smaller pieces in a petri dish on ice in order to increase the efficiency of the digestion. Tissue fragments were then transferred to a 15mL falcon tube containing 10mL of collagenase mix (RPMI-10%FBS containing 1mg/mL Collagenase IA (Sigma Aldrich, #C2674), 50microg/mL LiberaseTM ( Sigma Aldrich, #5401119001 ) and 0.1 mg/mL DNaseI (Sigma Aldrich, #4716728001)) at 37 °C for 45 minutes with rotation. Sample was then filtered (100 micron) and centrifuged 5min at 450g. Pellet was then resuspended in PBS-0.04%BSA and sorted for live/dead using ReadyDrop 7-AAD cell viability dye (Bio-Rad #1351102) in a Sony SH800 (https://benchling.com/s/prt-RVpSRQ8HG5rHg2ivx1C3 ).

Follicles isolated from the inner cortex remnants, were digested in Trypsin-EDTA 0.25% phenol red (Thermo Fisher Scientific #25200072) and 0.1 mg/mL DNaseI for 5 minutes at 37 °C then 5mL of PBS-0.04%BSA was added to stop the digestion, filtered (100 micron) and centrifuged 5min at 450g. Pellet was then resuspended in PBS-0.04%BSA and viability assessed by Trypan blue.

### Serial Cryosectioning for Paired Multi-omic Profiling

Ovarian cortex tissue from paediatric donors was processed for multimodal profiling by generating serial cryosections from each frozen OCT sample block. Cryosectioning was performed on a Leica CM3050S cryostat, with blocks trimmed until the full tissue surface was exposed.

Initial 10 μm sections were used to assess RNA integrity and tissue morphology. RNA was extracted using the Qiagen EZ2 RNA/miRNA Tissue Kit and quality was assessed via Agilent TapeStation RNA ScreenTape, with only samples achieving a RIN >7 retained for downstream molecular analysis. Additional sections were stained with Hematoxylin and Eosin (H&E) for histological validation.

Following quality control, each block was cryosectioned serially for parallel application to multiple spatial and single-cell profiling modalities, including 10x Visium spatial transcriptomics, Xenium In Situ, single-nucleus RNA-seq, and histopathological imaging. This approach enabled paired multi-omic measurements from near-adjacent sections of the same ovarian cortical regions.

### H&E staining and imaging

Slides holding fresh frozen sections were removed from -80 °C storage and air dried before being fixed in 10% neutral buffered formalin for 5 min. After being rinsed with deionized water, slides were dipped in Mayer’s haematoxylin solution for 90 s. Slides were completely rinsed in 4-5 washes of deionized water, which also served to blue the haematoxylin. Aqueous eosin (1%) was manually applied onto sections with a pipette and rinsed with deionized water after 1-3 s. Slides were dehydrated through an ethanol series (70, 70, 100, 100%) and cleared twice in 100% xylene. Slides were coverslipped and allowed to air dry before being imaged on a Hamamatsu NanoZoomer 2.0HT digital slide scanner.

### Single-nuclei extraction

Nuclei were isolated from ovarian OCT block cryosections (∼100 µm thickness) by homogenising tissue in a glass Dounce homogeniser (Sigma). Homogenisation was performed in cold nuclei isolation buffer supplemented with Protector RNase Inhibitor (Roche).

The suspension was filtered, centrifuged, and resuspended. An aliquot was stained with Trypan Blue, to assess nuclear integrity and concentration. Nuclei were then purified using a Percoll gradient, followed by a final Trypan Blue staining and count (https://dx.doi.org/10.17504/protocols.io.n2bvjxqewlk5/v1).

### Libraries preparation and sequencing

#### 10x Genomics Chromium GEX and ATAC

Single-cell and single-nucleus libraries were prepared using the 10x Genomics Chromium Single Cell Next GEM 3’ v3.1 or Next GEM 5’ v2 kits for transcriptomics, and Single Cell Next GEM ATAC v1.1 or v2 kits for ATAC, both according to the manufacturer’s instructions and with the aim to attain between 2,000 and 10,000 cells/nuclei per reaction. All libraries were sequenced on an Illumina-HTP NovaSeq 6000 platform with a S4 or SP flow cells, using paired-end sequencing, and to target a minimum of 50K reads per cell/nucleus. The sequencing format for scRNA-seq and snucRNA-seq was Read 1: [28] cycles; i7 index: [10] cycles; i5 index: [10] cycles; Read 2: [90] cycles. For scATAC-seq, the sequencing format was: Read 1: 50 cycles; i7 index: 8 cycles; i5 index: 16 cycles; Read 2: 50 cycles.

#### 10x Genomics Xenium

Ovarian tissue cryosections (∼10 µm thickness) from the OCT blocks were processed using the 10X Genomics Xenium Spatial Transcriptomics platform, following the manufacturer’s protocols and using a bespoke gene panel (named *hOvary_480g;* design id *EK2ZRP*). Briefly, samples were fixed with formaldehyde, permeabilised using SDS and methanol. Next, the tissue was incubated with target-specific probes for hybridisation, followed by ligation and rolling circle amplification for in situ signal enhancement. Prior to on-instrument analysis, autofluorescence quenching was performed. The slides were then stained with the Xenium Multi-Tissue Stain Mix as part of the Multimodal Cell Segmentation workflow. The mix combines antibodies targeting plasma membrane (ATP1A1/E-cadherin), intracellular proteins (Vimentin) and an 18S ribosomal RNA probe to delineate cytoplasmic compartment, complementing DAPI nuclear staining. Finally, slides were then loaded onto the Xenium Analyzer v2.0.1.0 together with decoding reagents and consumables for iterative imaging, barcode decoding and multimodal cell segmentation. This multimodal staining strategy enhanced the identification of cell boundaries for segmentation.

The custom probe panel was designed to target 480 genes that capture the diversity of cell states in the ovarian cortex across development (**Supplementary Table 2**). Genes were selected from a preliminary analysis of the scRNA-seq pediatric dataset and from our previously published fetal ovarian study, and were complemented by markers previously implicated in ovarian biology and ovarian conditions from the literature.

#### 10x Genomics Visium CytAssist

Ovarian tissue cryosections (10 µm) were cut from the OCT blocks onto 10x Genomics Visium slides and processed according to the manufacturer’s protocol. Briefly, sections were methanol-fixed, stained with haematoxylin and eosin (H&E), and imaged using a Hamamatsu NanoZoomer 2.0-HT slide scanner. Following imaging, sections were permeabilised, and spatially barcoded cDNA was generated by on-slide reverse transcription using template switching, followed by second-strand cDNA synthesis and PCR amplification. Resulting material was eluted from the slide and prepared into dual-indexed sequencing libraries according to the 10x Genomics protocol. Libraries were sequenced on an Illumina NovaSeq 6000 platform using SP flow cells, targeting a minimum of ∼25,000 read pairs per spot, with the sequencing format: Read 1, 28 cycles; i7 index, 10 cycles; i5 index, 10 cycles; Read 2, 90 cycles.

### Analysis of scRNA-seq and snucRNA-seq data

#### Preprocessing

For each library, sequencing reads were aligned to the GRCh38-2020-A reference genome, mRNA counts were quantified and initial quality control was performed using Cell Ranger v6.1.2 with default parameters. Called cells were scored for “doublet” likelihood using Scrublet, and classified to a cell cycle phase (that is, G1, G2/M or S) using the Scanpy function ‘score_genes_cell_cycle()’. Cell cycle and doublet scores were used to aid interpretation of downstream clustering but were not used to filter cells at this stage.

Genes were excluded if they were expressed in fewer than 3 cells in the dataset. Cells were excluded if they expressed fewer than 500 genes or had >20% mitochondrial content.

Each sample was analysed independently for quality control assessment. Briefly, after transforming expression values to log(CPM/100 + 1), we performed principal components analysis (PCA), nearest-neighbour graph construction, and Uniform Manifold Approximation and Projection (UMAP) and Leiden clustering to identify major cell type lineages.

#### Integration, clustering

To mitigate batch effects across donors and obtain a corrected low-dimensional representation of the gene expression, we applied the autoencoder-based method *scVI* (single-cell variational inference). Models were trained on the 2,000-5,000 most highly-variable genes, using a 64-dimensional space and one or two hidden layers, depending on whether the integration was performed: i) across all lineages jointly or ii) on a per-lineage basis (**Supplementary Figure 1a**). For the germ cell-specific analysis across the lifespan in the scRNA-seq modality, given the heterogeneous donor contributions to the number and developmental stage of germ cells and oocytes, we applied Harmony (theta = 0) to mitigate donor-associated batch effects. For the per-lineage integration within paediatric samples only in the snucRNA-seq modality, where donor-related batch effects were relatively modest, donor batch effects were likewise modelled using Harmony (theta = 0). In all cases, the resulting latent spaces were evaluated using the per-sample annotations, confirming the preservation of lineage/broad cell type identity while effectively reducing donor-specific batch effects.

The scVi batch-corrected low-dimensional representation was then used to construct the nearest-neighbour graph, UMAP and perform Leiden clustering (resolution adjusted manually to match well-known cell types) to identify: i) major cell lineages and cell types in the joint all-lineages analysis (referred to as “broad_annotation”) and ii) finer granularity cell state identities in the per-lineage analysis (referred to as “fine_annotation”).

#### Cell type annotation

To assign each cluster to the corresponding cell identity, we first identified their most characteristic genes by calculating the Term Frequency-Inverse Document Frequency (TF-IDF), as implemented in the R library SoupX. The approach binarises gene expression into 0/1 (representing not expressed or expressed genes in a cell, respectively) and quantifies how specifically a gene is associated with a cluster within the dataset. Cluster-specific genes were contrasted with literature marker genes for the matched lineage, which were used to guide cell type annotation (both “broad_annotation” and “fine_annotation”). Cell annotations and marker specificity was validated by analysing spatially-resolved transcriptomics data (both 10x Visium and In Situ Sequencing). Marker genes and the rationale underlying each annotation are provided in **Supplementary Note 3**.

Clusters that were likely driven by technical artefacts (that is, mostly composed of low-quality cells or doublets) were discarded. Briefly, we flagged as low-quality clusters those that (1) did not express any distinctive gene in the TF-IDF approach (and thus are not representing any independent biological entity) and, additionally, (2) expressed an overall lower number of genes, or (3) expressed an overall lower number of counts, or (4) displayed elevated mitochondrial or nuclear RNA content. Doublet clusters were flagged as those that met the following criteria: (1) did not express any distinctive gene in the TF-IDF approach and, additionally, (2) exhibited higher Scrublet doublet scores or (3) expressed marker genes from multiple lineages (for example, granulosa and immune markers).

#### Granulosa and macrophage trajectory inference

The granulosa and macrophage differentiation trajectory was inferred using Slingshot, with squamous granulosa cells (primordial follicles) or monocytes defined as the starting point, respectively. The resulting pseudotime values and cell-to-lineage weights were provided to the TradeSeq framework. The associationTest() function was then used to extract genes exhibiting significant differential expression along the inferred trajectory.

### Analysis of scATAC-seq data

#### Preprocessing

scATAC data were processed with Cell Ranger-Atac v2.0.0 with default parameters. Downstream analysis was performed in ArchR (v1.0.3) following standard pipeline.

#### Integration, clustering

First, fragments were quantified in genome wide tiles, cells with less than 500 fragments or with TSS enrichment score below 10 were removed. Then we performed interactive LSI for dimension reduction and clustering at resolution of 1.

#### Cell type annotation

Cell type annotation was performed using addGeneIntegrationMatrix function with GeneScoreMatrix in two steps. First, labels were transferred from the scRNA-seq modality using the full dataset, to assign lineage-level annotation. Lineage-level labels were refined using majority voting at the cluster level. OSE cells were picked manually as a tight cluster of cells (at resolution of 10) that present the highest gene score for the corresponding marker genes *UPK3B, LHX2, LRRN4*. Second, label transfer was repeated in a per-lineage manner using the “broad_annotation” from the scRNA-seq modality (except for OSE) again applying the majority voting across clusters obtained at a clustering resolution of 2. The resulting cell type annotations were used for peak calling by MACS2 via the ArcR wrapper. Called peaks were annotated with transcription binding motifs from CIS-BP and JASPAR-v2020 databases using the ArchR::addMotifAnnotations function.

#### Identification of DARs and TF-binding motifs along early granulosa differentiation

For the granulosa cell-specific analysis, granulosa cells were integrated using Harmony with donor modelled as batch, followed by reclustering at resolution of 2. Clusters in which more than half of the cells had low coverage (<10,000 fragments) or were identified as doublets. As result, 3,437 cells out of 3,984 originally annotated as granulosa cells were kept. Cluster labels were then re-transferred using only cortical granulosa reference labels (Granulosa_sq, Granulosa_transitioning, Granulosa_AMH_early and Granulosa_AMH_multilayered), given that the scATAC-seq libraries were exclusively derived from cortical tissue rather than remnant material. To ensure balanced comparisons between granulosa states, each granulosa population was downsampled to an equal number of cells (n=278 cells). Differentially Accessible Regions/Peaks (DARs) were identified using ArchR::getMarkerFeatures function comparing: i) Granulosa_sq (i.e. the quiescent granulosa state) versus all other granulosa states combined; and ii) each granulosa state vs Granulosa_sq. Finally, a hypergeometric test was applied to quantify transcriptional factor binding motifs enriched in the DARs.

### Analysis of Xenium In Situ data

Default on-instrument multimodal cell segmentation was performed on Xenium datasets using DAPI and multistaining signals with Xenium Analyzer v2.0.1.0.

#### Preprocessing

Cell-assigned Xenium gene expression data were pre-processed using standard Scanpy workflows. Only cells with a segmented area ≥ 25, total transcript counts > 50, and a nucleus count ≤ 1 were retained, thereby excluding very small, low-coverage or multinucleated segments. Some sections were analysed independently for quality control assessment. Briefly, after transforming expression values to log(CPM/100 + 1), PCA was performed followed by nearest-neighbour graph construction, UMAP and Leiden clustering to identify major cell lineages.

#### Integration, clustering

To correct batch effects across donors and obtain a corrected low-dimensional representation of Xenium gene expression, we applied ResolVI, using donor as the batch variable. Models were trained on the 400 most highly variable genes with the following parameters: 40 latent dimensions, 2 hidden layers with 128 units each, negative binomial gene likelihood (gene_likelihood = "nb"), gene-wise dispersion (dispersion = "gene"), dropout rate of 0.1, and a maximum of 100 training epochs. Integration was performed either (i) across all lineages jointly or (ii) on a per-structure/lineage basis (run 1-vessels: endothelial and perivascular cells; run 2-stroma and follicles: theca, granulosa, oocyte and stromal cells; and run 3-immune: myeloid and lymphoid cells). In all cases, the resulting latent spaces were evaluated using per-sample annotations, confirming preservation of lineage and broad cell type identity while reducing donor-specific batch effects.

The ResolVI batch-corrected latent representation was then used to construct the nearest-neighbour graph, compute UMAP embeddings, and perform Leiden clustering, with the clustering resolution adjusted manually to align with well-established cell types identified in the scRNA-seq/snucRNA-seq datasets. This analysis was used to identify (i) major cell lineages and cell types in the joint all-lineage integration (referred to as “broad_annotation”) and (ii) higher-resolution cell-state identities (referred to as “fine_annotation”) in three per-lineage analyses.

#### Cell type annotation

Assignment of cell identities to Xenium-derived clusters followed an approach analogous to that used for the scRNA-seq and snucRNA-seq datasets. Here, differential gene expression analysis was performed with the Scanpy.rank_genes_groups function to identify the most strongly upregulated genes in each cluster. These differentially expressed genes were then compared to marker genes identified in the scRNA-seq/snucRNA-seq analysis and used to guide cell type annotation of Xenium-segmented cells at both the “broad_annotation” and “fine_annotation” levels.

Clusters exhibiting a clear doublet-like signature, likely corresponding to improperly segmented cells, and characterised by the co-expression of marker genes from multiple lineages (for example, granulosa and immune markers) and absence of a distinctive marker profile were annotated as “Mixed” populations.

#### Spatial cell type mapping and tissue niche identification

Spatial locations of the cell types defined in the scRNA-seq dataset were validated by projecting these annotations onto the Xenium-profiled cells using DOT^97^. The approach reconstructs the in situ spatial organisation of cells by leveraging spatial proximity, local gene expression similarity, and modality-specific features to accurately align the single-cell profiles with the Xenium spatial coordinates.

#### Spatial cell type Neighbourhood Enrichment Analysis

To quantify spatial enrichment between DOT annotated cell types, we performed a spatial cell type neighbourhood enrichment analysis. For every Xenium slide, a k-nearest-neighbour graph (kNN = 3) was constructed using Euclidean distances between cell centroids. Using this graph we can list every cell pair consisting of a query cell type (A) and a candidate neighbouring cell type (B), and subsequently generate a 2×2 contingency table comparing the number of A-B edges to the number of A-non-B edges. A Fisher’s exact test was then applied to determine whether cells of type B occurred as statistically enriched or depleted neighbours of query cell type A. P-values were corrected using the Benjamini-Hochberg method and effect sizes were reported as odds ratios.

#### Common coordinate framework for spatial axis inference

We construct a common coordinate framework for ovarian tissue using spatial_axis (github.com/uhlmanngroup/spatial_axis), which is conceptually similar to that described in^57,58^. A notable distinction in our method is that we derive spatial annotations from transcriptomic cell type annotation, rather than annotations created manually or from a pixel classifier. Additionally, our method supports any number of classes, allowing us to construct a continuous spatial axis across multiple morphological landmarks. We provide an extended description in the **Supplementary Note 4**. Briefly, spatial_axis defines for each cell a value in respect to morphological landmarks (for example, stromal layers). For each cell, we calculate the distance to K=15 nearest neighbours for each morphological landmark. Although the number of nearest neighbours is a tuneable hyperparameter, we selected 15 as it captured a smooth gradient across stromal layers for the donors tested. These distances are then aggregated. We then compute the normalised distance for each cell, considering only morphological landmarks that are spatially adjacent.

#### Label transfer from Xenium to scRNA-seq with ISS-Patcher

To enable spatial analysis of the scRNA-seq data, we mapped the cell type annotations and spatial coordinates from the Xenium cells onto dissociated cells in the scRNA-seq dataset. Using a modified k-nearest neighbours (kNN) approach implemented in the iss_patcher library (github.com/Teichlab/iss_patcher), we imputed the spatial coordinates for scRNA-seq cells by mapping their transcriptomic profiles to their nearest Xenium-profiled neighbours. This procedure assigned each scRNA-seq cell a position along both the cortico-medullary and follicular spatial axes.

To identify genes with expression patterns varying with the cortico-medullary spatial position of the stromal fibroblasts (spatially variable genes), we utilized the TradeSeq associationTest() function, which models gene expression along the imputed spatial axes as pseudotime-like trajectories in the scRNA-seq stromal fibroblasts. For modeling gen expression dynamics along the follicular axes, the analysis was extended to include theca cells and perifollicular fibroblasts as well.

#### Expert Annotation of Oocytes

Expert researchers in oocyte biology annotated digitised H&E-stained ovarian sections to identify histologically distinct follicular structures. Annotations focused on the presence, developmental stage, and localisation of oocytes within the ovarian cortex. Morphological criteria for staging were guided by established ovarian histology references to ensure consistency and biological relevance.

### Analysis of Visium cytassist

Visium spatial gene expression preprocessing was performed using Space Ranger v2.0.0 (10x Genomics), aligned to the GRCh38-2020-A reference genome to match the reference used for the single-cell RNA-seq dataset.

#### Spatial cell type mapping

To infer the spatial distribution of cell types across the Visium sections, we applied cell2location (v0.8), a Bayesian model that integrates single-cell RNA-seq data as reference to estimate the abundance of each cell type within individual spatial transcriptomics spots.^98^ Expression signatures were derived from the scRNA-seq dataset and used to decompose Visium gene expression profiles into spatial maps of cell type abundance. Each tissue section was analysed independently. We visualised the absolute contribution of each cell type to each spot, focusing on confidently assigned mRNA signatures.

### Cell-cell communication analysis with CellPhoneDB

To study potential cell-cell interactions, we first grouped cell types into spatial neighbourhoods (or microenvironments) where they coexist. Spatial location was assigned based on histological inspection (for example, follicular cells) or spatial neighbourhood enrichment analysis from label-transfer cell type predictions on Xenium cells from scRNAseq signatures using DOT (for example, for macrophages).

We then used the DEGs-based method in CellPhoneDB v4.0.0 to identify putative paracrine and juxtacrine interactions between co-localising cell types in the ovary. CellPhoneDB retrieved the ligand-receptor pairs that met the following criteria: (1) all interacting partners were expressed in at least 10% of cells in the relevant cell type; (2) the interacting cell type pairs co-localised; and (3) at least one partner in the pair (ligand or receptor) was significantly upregulated in the corresponding cell state compared with other cells within the same lineage (Wilcoxon test; adjusted P < 0.01 and log2 fold change > 0.75).

Differential expression analysis was performed on a per-lineage basis to identify genes specifically upregulated in a given cell state relative to other states within that lineage (for example, oLAMs versus all other macrophage states).

### Metabolic activity predictions with scCellFie

To reconstruct metabolic pathway activity from our scRNAseq data, we used scCellFie^84^ to map gene expression profiles onto a curated set of metabolic tasks. scCellFie computes task activity by applying gene protein reaction rules to generate a quantitative representation of metabolic function for each cell. Differential metabolic activity analysis was performed between macrophage cell types using the scCellFie differential activity module, which performs group-wise comparisons of task activity scores and applies Benjamini Hochberg FDR correction to identify significantly altered metabolic programmes. Plotting and visualisation of task activity profiles were performed using the scCellFie plotting module.

### Compositional analysis of cellular abundances

We evaluated cellular composition changes between major physiological transitions at postnatal (2-3 years), pre-puberty (4-10 years), and post-puberty (10-19 years) lifespan stages using tascCODA^99^, a Bayesian method that accounts for both the compositional nature of the data and hierarchical relationships between features (e.g. families of cell types such as all stromal cell types together). We used an aggregation bias parameter ϕ = 5.0, as indicated in the tascCODA methodology^99^, to favor detection of leaf-level effects (i.e. cell type specific changes). As a Bayesian method, tascCODA does not produce p-values; instead, it finds credible effects, which are determined using posterior median effect sizes that exceed node-specific thresholds of practical significance, as described by Ostner et al. ^99^.

### Donor’s genetic ancestry inference

All samples were subjected to genotyping analysis to determine genetic ancestry, as defined by the National Academies of Sciences, Engineering, and Medicine (NASEM)^100^ and inspired by the gnomAD approach^101^. For this, we employed variant calling using freeBayes, followed by the random forest classifier trained on gnomAD data (as described in https://gnomad.broadinstitute.org/help/ancestry) to assign the ancestry labels^100,102,103^. The reference ancestries include African (AFR), Amish (AMI), Amerindian Latino (AMR), Ashkenazi Jewish ancestry (ASJ), East Asian (EAS), Finnish (FIN), Middle Eastern (MID), Non-Finnish European (NFE), South Asian (SAS) and other (OTH).

## Data and code availability

We provide user-friendly access to our annotated scRNA-seq resource via cellxgene at our https://www.reproductivecellatlas.org/ovaries/sanger_cells/ and to our Xenium-registered multimodal dataset at https://webatlas.sanger.ac.uk. All the raw and processed sequencing data generated in this study are currently being deposited to EGA (scRNA-seq, scATAC-seq), ArrayExpress (scRNA-seq, scATAC-seq, 10x Visium) and BioImageArchive (10x Xenium). The code used to perform the analyses presented in the manuscript can be found at <u>github.com/uhlmanngroup/spatial_axis</u> and <u>github.com/ventolab/Human-Ovarian-Atlas/</u>.

## Acknowledgements

We wish to acknowledge the donors, patients and relatives who provided ovarian tissues for research. We also acknowledge members of staff at the Oxford Cell and Tissue Biobank (OCTB) for their assistance in obtaining the pediatric ovarian tissue samples; the Newcastle upon Tyne hospital‘s clinical research and gynaecology team for their support in donor recruitment, informed consent procedures and obtaining the adult ovarian tissue samples; the transplant organ donors and their families for their generous tissue donations through the Cambridge Biorepository for Translational Medicine; Dr. Felix Day and Prof. John R.B. Perry for providing access to PCOS summary statistics from their preprint^104^Dr. Valentina Lorenzi and Dr. Nadav Yayon for insightful discussions on the spatial transcriptomics strategy and analysis; B. Cakir for his help on the reproductivecellatlas.org web portal; Tarryn Porter, Heather Stanley, Steven Leonard and Yong Gu for support with the sequencing pipeline and the Spatial Genomics Platform (SGP) and Sanger Core Sequencing pipeline for support with sample processing and sequencing library preparation; A. García from Bio-Graphics for scientific illustrations; and A. Maartens for his help on manuscript writing. Bio-Graphics for scientific illustrations; and A. Maartens for his help on manuscript writing. This publication is part of the Human Cell Atlas – www.humancellatlas.org/publications/.

## Author contributions

L.G-A and R.V-T conceived and designed the study, with contributions from S.A.W and V.U. M.H, K.T.M, K.S-P, M.C, J.D and S.L contributed to ovarian sample acquisition. C.I.M, L.T, E.T, F.A, M.K, A.J and L.W performed sample dissections. C.S.S and E.P performed sample processing for single-cell profiling, with contributions from J.A, H.M, G.M and A.M, supervised by I.K. N.K, L.T and C.I.M performed sample processing for the imaging experiments, with contributions from J.A. L.G-A coordinated data analysis, which was performed by L.G-A, J.A, C.T-C, C.C and P.M, with contributions from E.A, T.L, A.B, A-V.P, K.T and M.P. C.T-C performed histopathological imaging data analysis, with contributions from T.L and M.P. C.T-C and N.K performed histological annotation of oocytes and follicles, with contributions from L.G-A and J.A. L.G-A, J.A, C.C and R.V-T interpreted the data, with contributions from A.P. L.G-A and R.V-T wrote the manuscript with contributions from J.A, C.T-C, N.K, C.C, P.M, M.H, V.U and S.A.W. L.G-A and R.V-T supervised the work; S.A.W supervised pediatric tissue management, and V.U supervised computational imaging analyses. All authors read and approved the manuscript.

## Supplementary Figures Legends

**Supplementary Figure 1.**
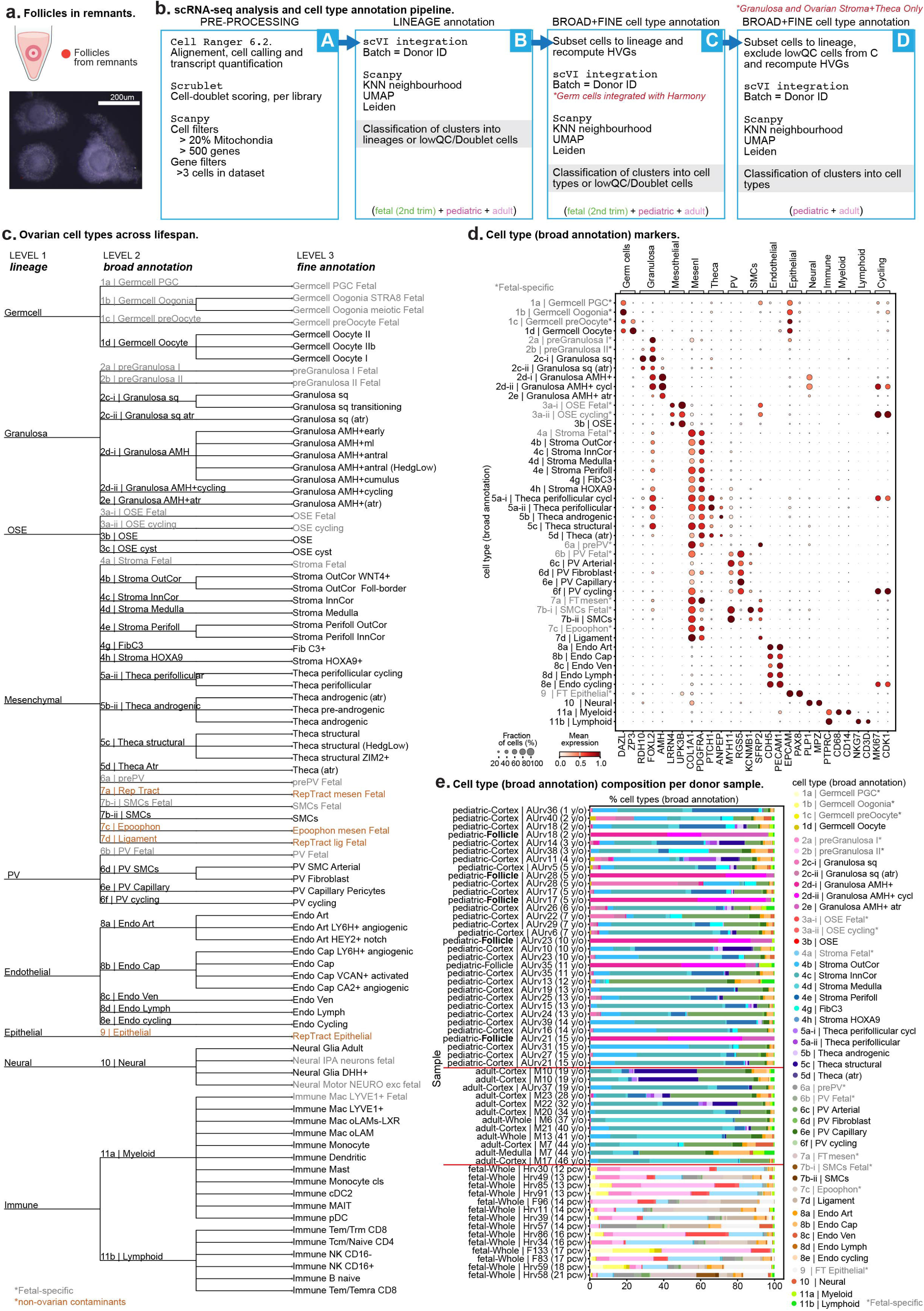
scRNAseq analysis overview and ovarian cell type nomenclature. **a.** Representative light microscopy images of intact follicles isolated from remnant fragments from the cortical strip dissection. Scale bar = 200µm. **b.** Schematic representation of the computational workflow used to analyse scRNA-seq data. **c.** Schematic representation of ovarian cell type hierarchy and nomenclature. The tree illustrates the taxonomic organization across three levels of increasing granularity from major lineage to broad and fine-resolution annotations. Tree built with *itol.embl.de*. **d.** Dot plot showing the variance-scaled, log-transformed expression of marker genes (x-axis) characteristic of the annotated cell types (y-axis; broad annotation). Top-layer groups marker genes by major lineage. **e.** Bar plot showing the proportion of cell types per scRNA-seq libraries, grouped by donor and sampling type, and colored by their broad annotation.

**Supplementary Figure 2.**
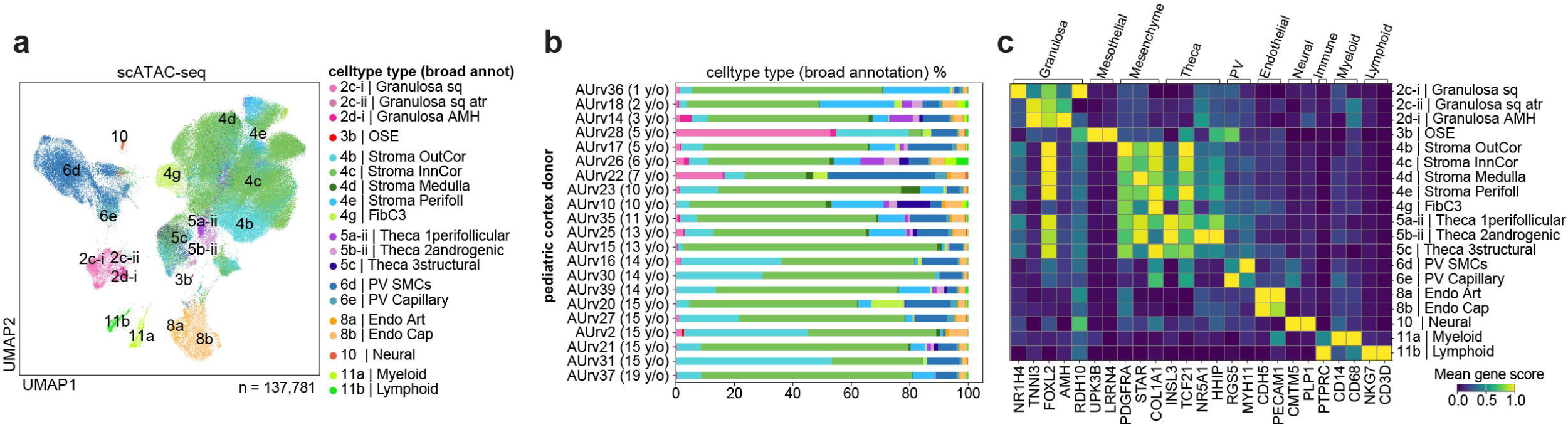
scATACseq analysis overview. **a.** Batch-corrected Uniform Manifold Approximation and Projection (UMAP) embedding of the scATAC-seq dataset (n = 137,781 cells) coloured by broad cell type annotation, transferred from scRNA-seq dataset with ArchR. **b.** Bar plot showing the proportion of cell types in the scATAC-seq libraries per donor, colored by their broad annotation. **c.** Heatmap plot showing the variance-scaled, accessibility scores of marker genes (x-axis) characteristic of the annotated cell types (y-axis; broad annotation). Top-layer groups marker genes by major lineage.

**Supplementary Figure 3.**
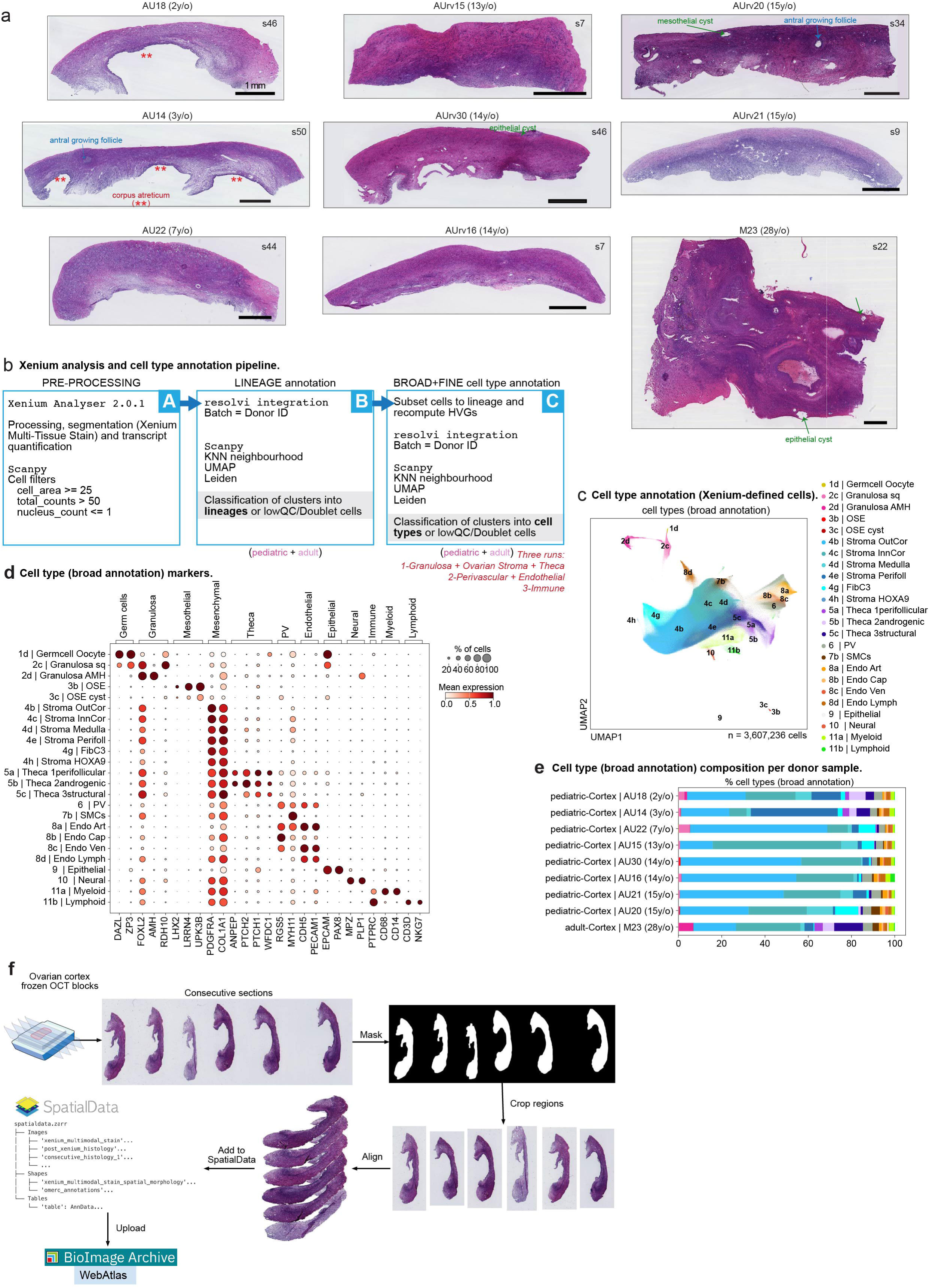
Xenium and imaging analysis overview. **a.** Representative Hematoxylin and eosin (H&E) stained sections from each of the eight ovarian cortical strips from pediatric donors and the cortical biopsy from an adult donor profiled by 10x Xenium In Situ Spatial Transcriptomics; donor ids and age indicated at the title. Scale bar = 1mm. **b.** Schematic representation of the computational workflow used to analyse the transcriptomic profiles from Xenium segmented cells for direct cell type annotation. **c.** Batch-corrected Uniform Manifold Approximation and Projection (UMAP) embedding of the Xenium segmented cells (n = 3,607,236 cells) coloured by broad cell type annotation, annotated based on marker genes. **d.** Dot plot showing the variance-scaled, log-transformed expression of marker genes (x-axis) characteristic of the annotated cell types (y-axis; broad annotation). Top-layer groups marker genes by major lineage. **e.** Bar plot showing the proportion of cell types per donor cortical biopsy, colored by their broad annotation. **f.** Schematic representation of the pipeline for aligning H&E images from consecutive sections and registering multiple spatial modalities (H&E, histological annotation, gene expression, multimodal staining, spatial axis coordinates and cell type annotation) into a SpatialData object, made available through WebAtlas and the BioImage Archive.

**Supplementary Figure 4.**
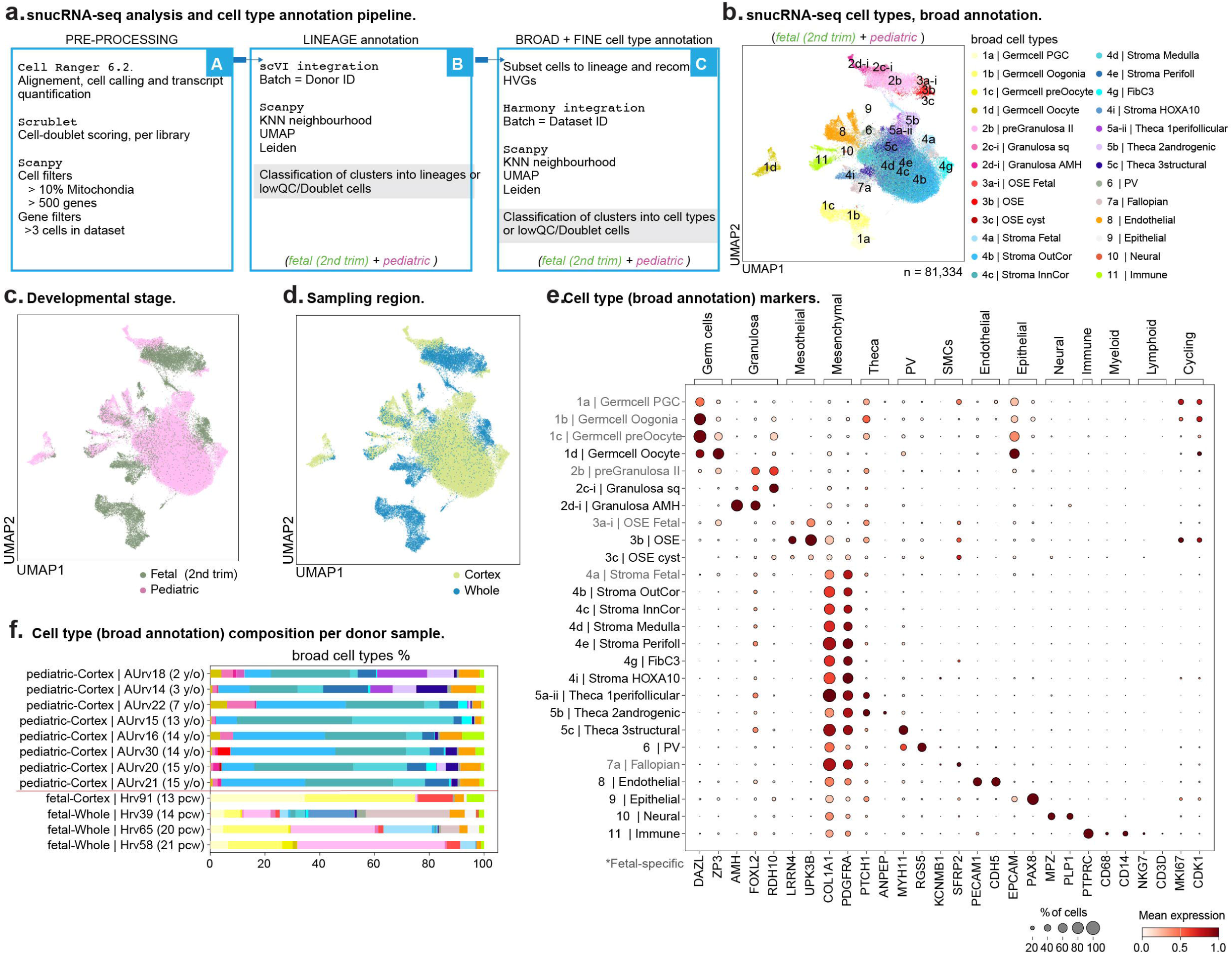
snucRNAseq analysis overview. **a.** Schematic representation of the computational workflow used to analyse snucRNA-seq data. **b-d.** Batch-corrected Uniform Manifold Approximation and Projection (UMAP) embedding of the snucRNA-seq dataset (n = 81,334 cells) coloured by broad cell type annotation, annotated based on marker genes (“b”), developmental stage (“c”) and “sampling region” (“d”). **e.** Dot plot showing the variance-scaled, log-transformed expression of marker genes (x-axis) characteristic of the annotated cell types (y-axis; broad annotation). Top-layer groups marker genes by major lineage. **f.** Bar plot showing the proportion of cell types in the snucRNA-seq libraries per donor, colored by their broad annotation.

**Supplementary Figure 5.**
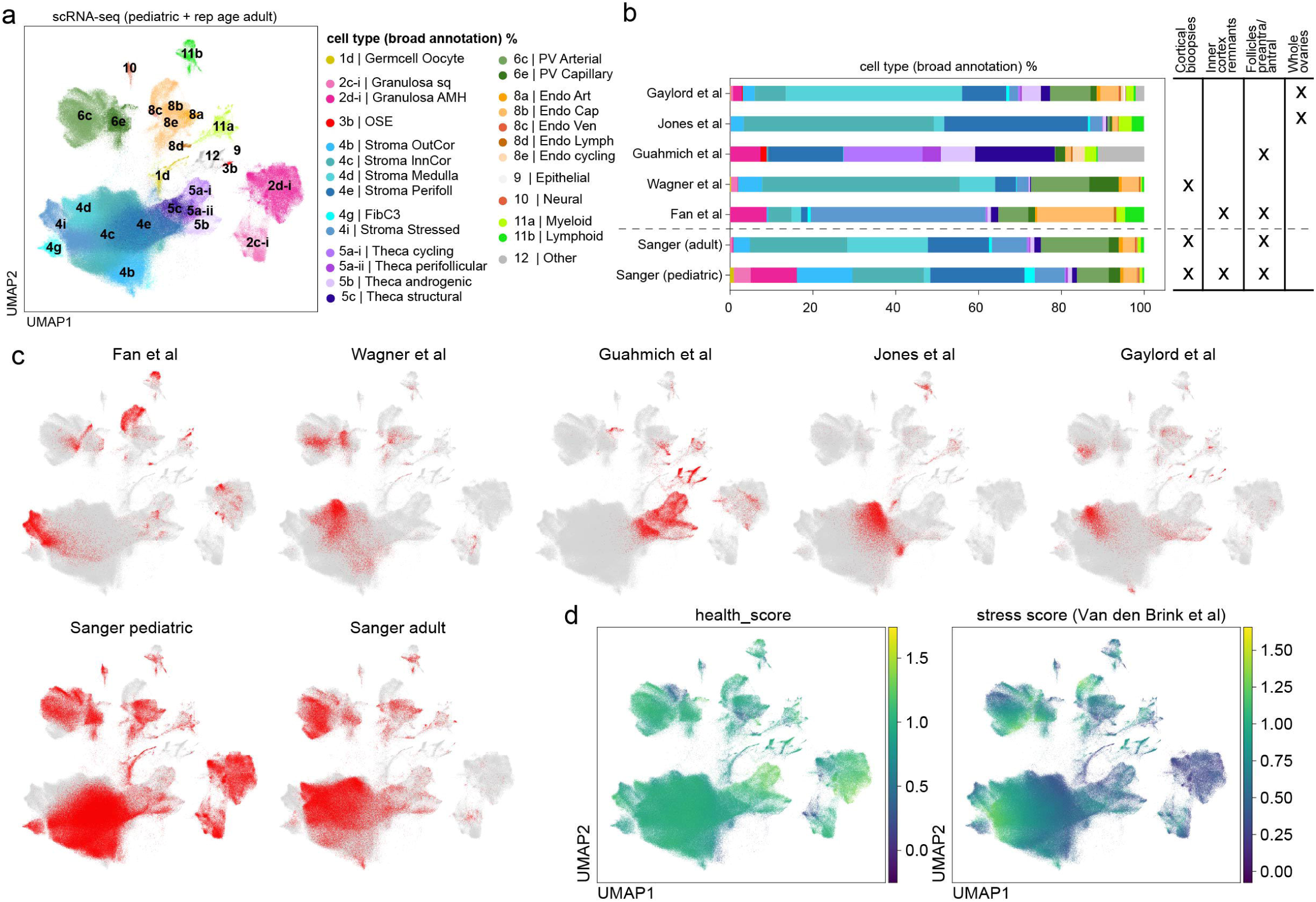
Integration and comparison with existing scRNAseq data from human adult ovaries. **a.** Batch-corrected Uniform Manifold Approximation and Projection (UMAP) embedding of our and existing scRNA-seq datasets combined (n = 675,119 cells) coloured by broad cell type annotation, annotated based on marker genes. **b.** Bar plot showing the proportion of cell types in each scRNA-seq dataset, colored by broad annotation (left); and table indicating the biopsy type profiled by each dataset (right). **c.** UMAPs, as in (a), coloured by each of the datasets used in the integration. **d.** UMAPs, as in (“a”), coloured by health score (left) and stress score (right).

**Supplementary Figure 6.**
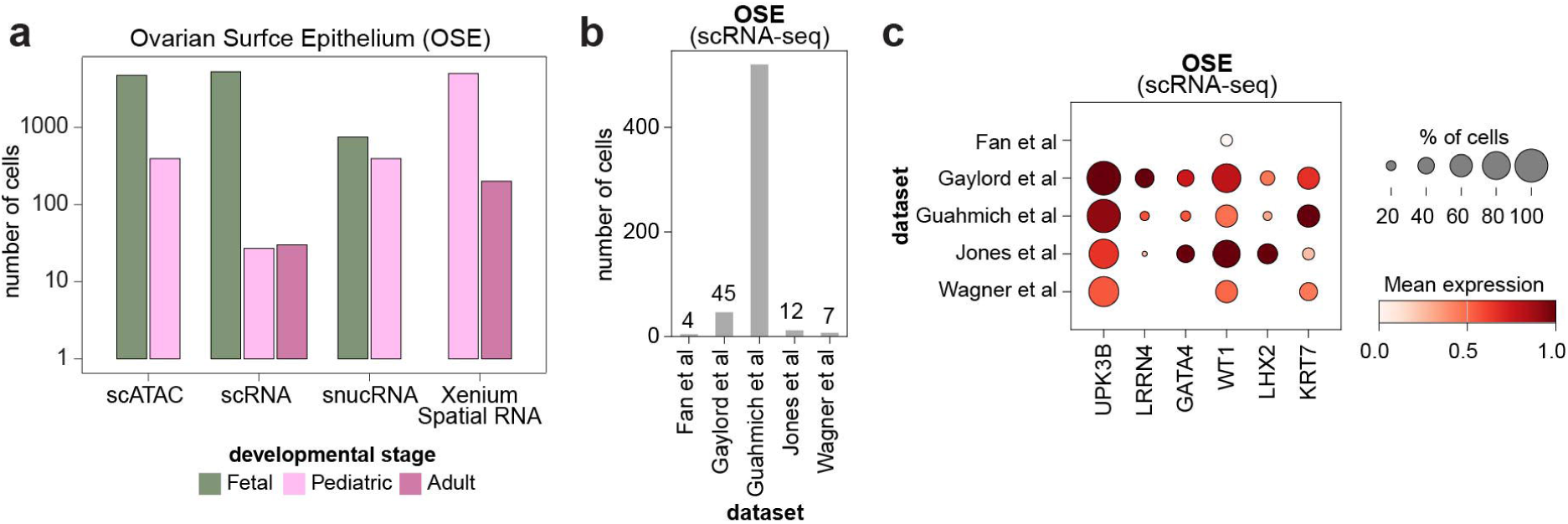
OSE and other aberrant epithelial cells in the ovary. **a.** Barplot showing the number of canonical OSE cells identified though development age and data modality. **b**. Bar plots showing the number (y-axis) of OSE cells (broad cells; annotated directly on the integrated UMAP) in each of the publicly available scRNA-seq datasets analysed (x-axis). **c.** Dotplot showing the log-transformed, min-max normalised expression of OSE markers (x-axis) along OSE cells from “b” (y-axis), in each of the publicly available scRNA-seq datasets analysed.

**Supplementary Figure 7.**
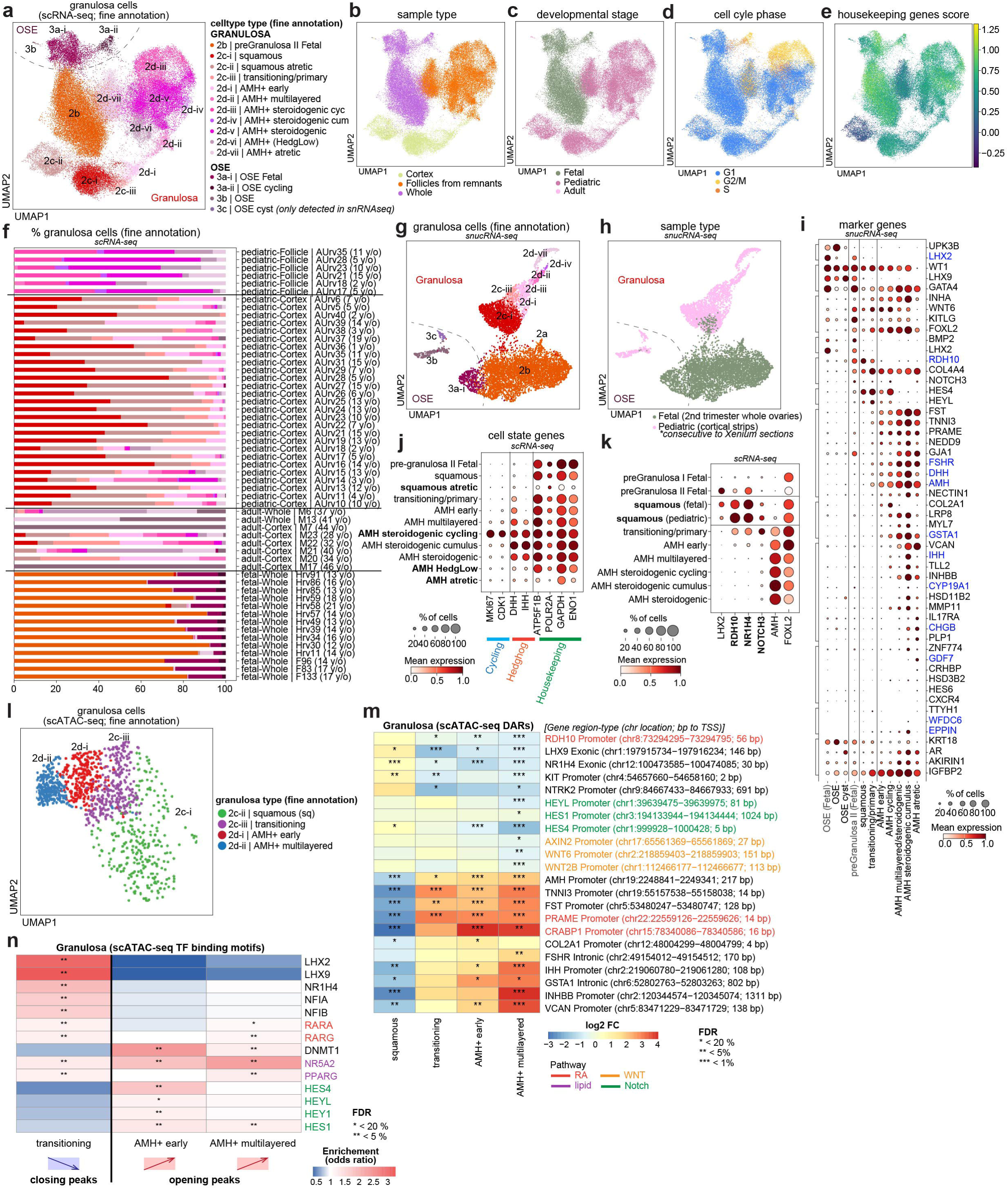
Granulosa cells characterisation. a-e. Batch-corrected Uniform Manifold Approximation and Projection (UMAP) embedding of the granulosa-annotated cells in the scRNA-seq dataset (n = 63,672 cells) coloured by fine cell type annotation (a), sample type (b), developmental stage (c), cell cycle phase (d) and average expression (score) of housekeeping genes (e). **f.** Bar plot showing the proportion of granulosa and OSE cell types per scRNA-seq library (x-axis), grouped by donor and sampling type (y-axis), and colored by their fine annotation, color legend in “a”. **g-h.** UMAP embedding of the granulosa and OSE annotated cells in the snucRNA-seq dataset (n = 8,538 nuclei) coloured by fine cell type annotation (g; color legend in “a”) and developmental stage (h). **i.** Dotplot showing the log-transformed, min-max normalised expression of selected marker genes of granulosa and OSE (x-axis) along granulosa and OSE cells (y-axis) in the snucRNA-seq dataset. **j.** Dotplot showing the log-transformed, min-max normalised expression of cell state marker genes (x-axis) along granulosa cells (y-axis) in the scRNA-seq dataset. **k.** Dotplot showing the log-transformed, min-max normalised expression of squamous granulosa marker genes (x-axis) along granulosa cells (y-axis) in the scRNA-seq dataset; with fetal and postnatal squamous granulosa separated. **l.** UMAP embedding of the granulosa cells scATA-seq profiles, coloured by fine cell type annotation, transferred from scRNA-seq dataset with ArchR. **m.** Heatmap showing the log2 fold change for selected “differentially accessible regions” (DARs) in genes relevant to early granulosa differentiation (y-axis), resulting from comparing each granulosa cell state (x-axis) against squamous granulosa cells (for non-squamous granulosa states) or against all remaining granulosa cells (for squamous granulosa). Asterisks denote FDR significance (* < 20%; ** < 5%; *** < 1%). **n.** Heatmap showing the odds ratio enrichment for selected DNA-binding protein motifs relevant to early granulosa differentiation (y-axis), in the DARs downregulated in the transitioning-vs-squamous, in the DARs upregulated in the early-vs-squamous and in the DARs upregulated in the ml-vs-squamous comparisons (x-axis). Asterisks denote FDR significance (* < 20%; ** < 5%).

**Supplementary Figure 8.**
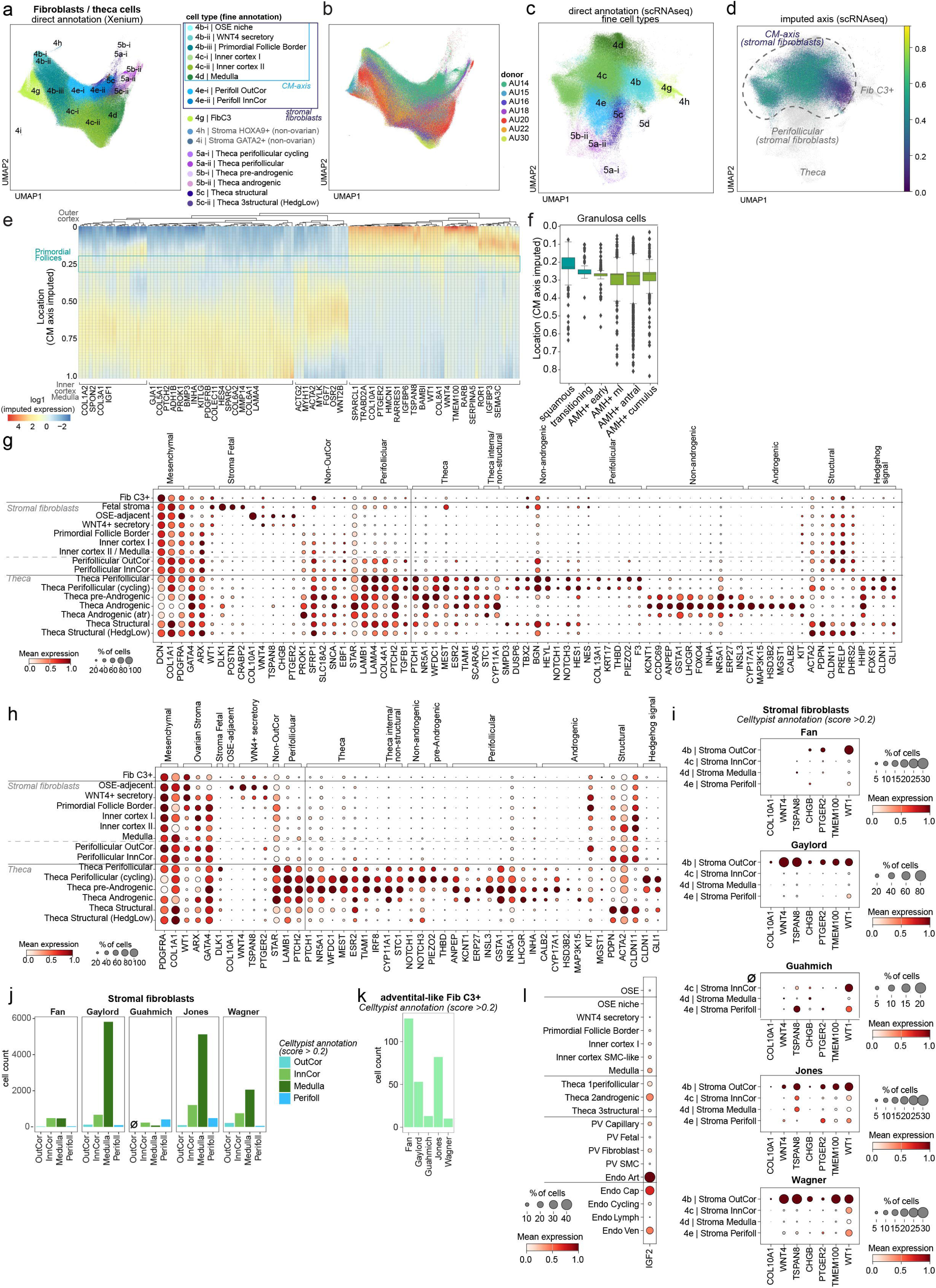
Stromal fibroblast “*Cortico-medullary spatial axis*”. a-b. Batch-corrected Uniform Manifold Approximation and Projection (UMAP) embedding of the stromal mesenchymal cells in the Xenium dataset coloured by fine cell type annotation, where stromal fibroblasts are coloured according to their assigned fibroblast strata based on their spatial coordinate in the “*Cortico-medullary axis*” (“a”) and colored by donor ID (“b”), respectively. **c-d.** UMAP embedding of the stromal mesenchymal cells in the scRNA-seq dataset coloured by fine cell type annotation (same color legend as in “a”) and their spatial coordinate along the iss-patcher imputed “*Cortico-medullary axis*”. **e.** Heatmap representing the z-score, log-transformed expression of spatially variable genes (x-axis) along the iss-patcher imputed “*Cortico-medullary axis*” (y-axis) identified in stromal fibroblast cells from the scRNA-seq dataset. **f.** Boxplots showing the iss-patcher imputed “*Cortico-medullary axis*” spatial location (y-axis) for granulosa cell subtypes (x-axis). **g-h.** Dotplots showing the log-transformed, min-max normalised expression of cell state marker genes (x-axis) along the stromal fibroblast subtypes (y-axis) in the scRNA-seq dataset (“g”) and Xenium segmented cells (“h”). **i.** Dotplots showing the log-transformed, min-max normalised expression of WNT4+ secretory fibroblast markers (x-axis) along stromal fibroblasts subtypes (y-axis; cells annotated via CellTypist label transfer from our scRNAseq dataset), in each of the publicly available scRNA-seq datasets analysed. **j.** Bar plots showing the number (y-axis) of stromal fibroblast subtypes (x-axis; cells annotated via CellTypist label transfer from our scRNAseq dataset), in each of the publicly available scRNA-seq datasets analysed. **k.** Bar plots showing the number (y-axis) of C3+ adventitial-like fibroblasts (Fib C3+; cells annotated directly on the integrated UMAP) in each of the publicly available scRNA-seq datasets analysed (x-axis). **l.** Dotplot showing the log-transformed, min-max normalised expression of *IGF2* (x-axis) along OSE, stromal fibroblast, theca, perivascular (PV) and endothelial subtypes (y-axis) in the scRNA-seq dataset.

**Supplementary Figure 9.**
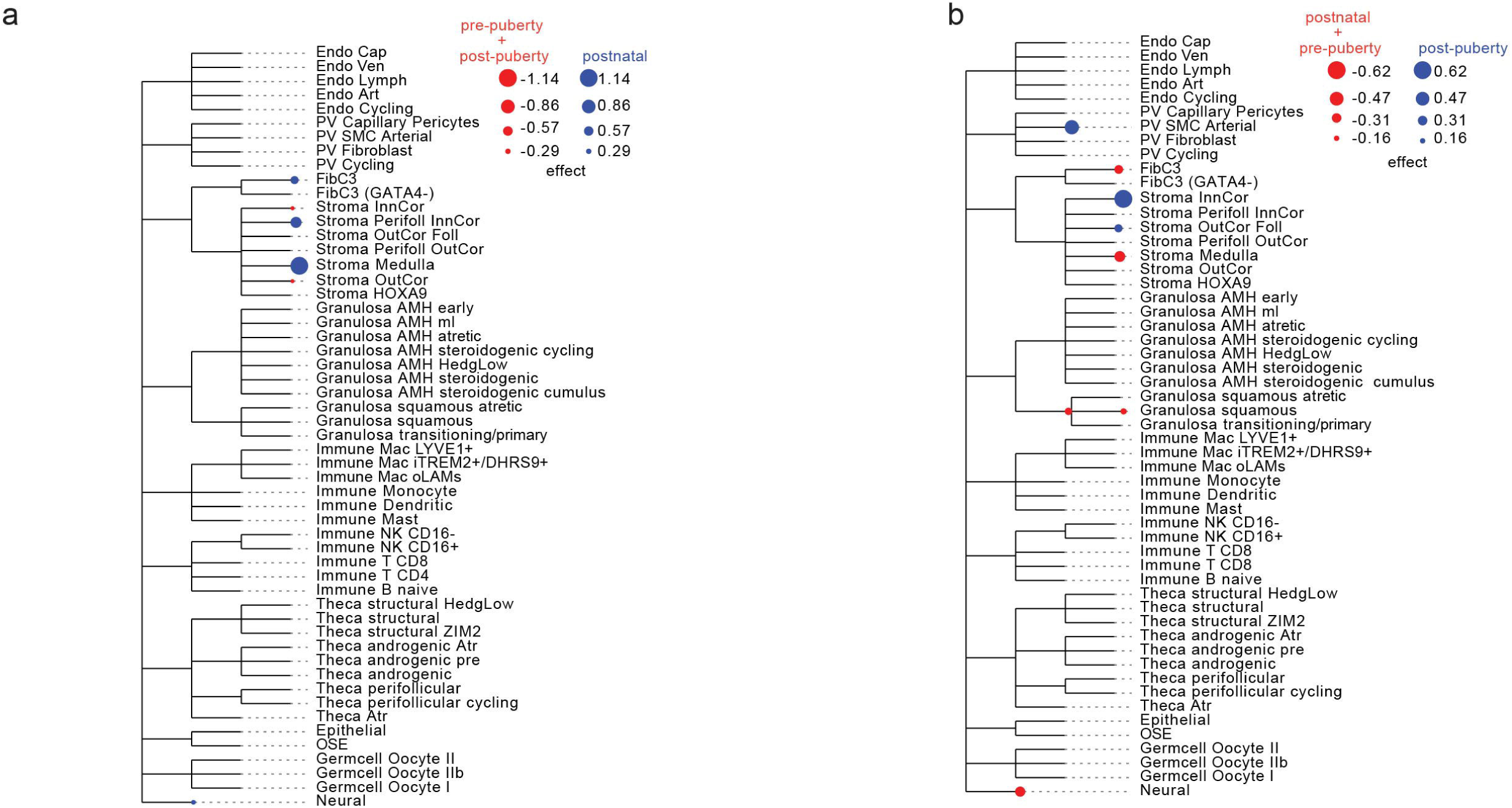
cell type enrichment across childhood stages. a-b. Cell enrichment analysis at each lifespan stage, using tascCODA with hierarchical smoothing (Φ = 5.0) in a “lifespan stage vs. the rest” design for postnatal samples vs. pre-puberty + post-puberty samples (“a”) , and post-pubertal samples vs. postnatal + pre-pubertal samples (“b”). Significant positive effect sizes indicate enrichment in the focal stage relative to both other stages. Blue circles denote positive effect sizes (enrichment) and red circles denote negative effect sizes (depletion). Circle size scales with the magnitude of the effect size.

**Supplementary Figure 10.**
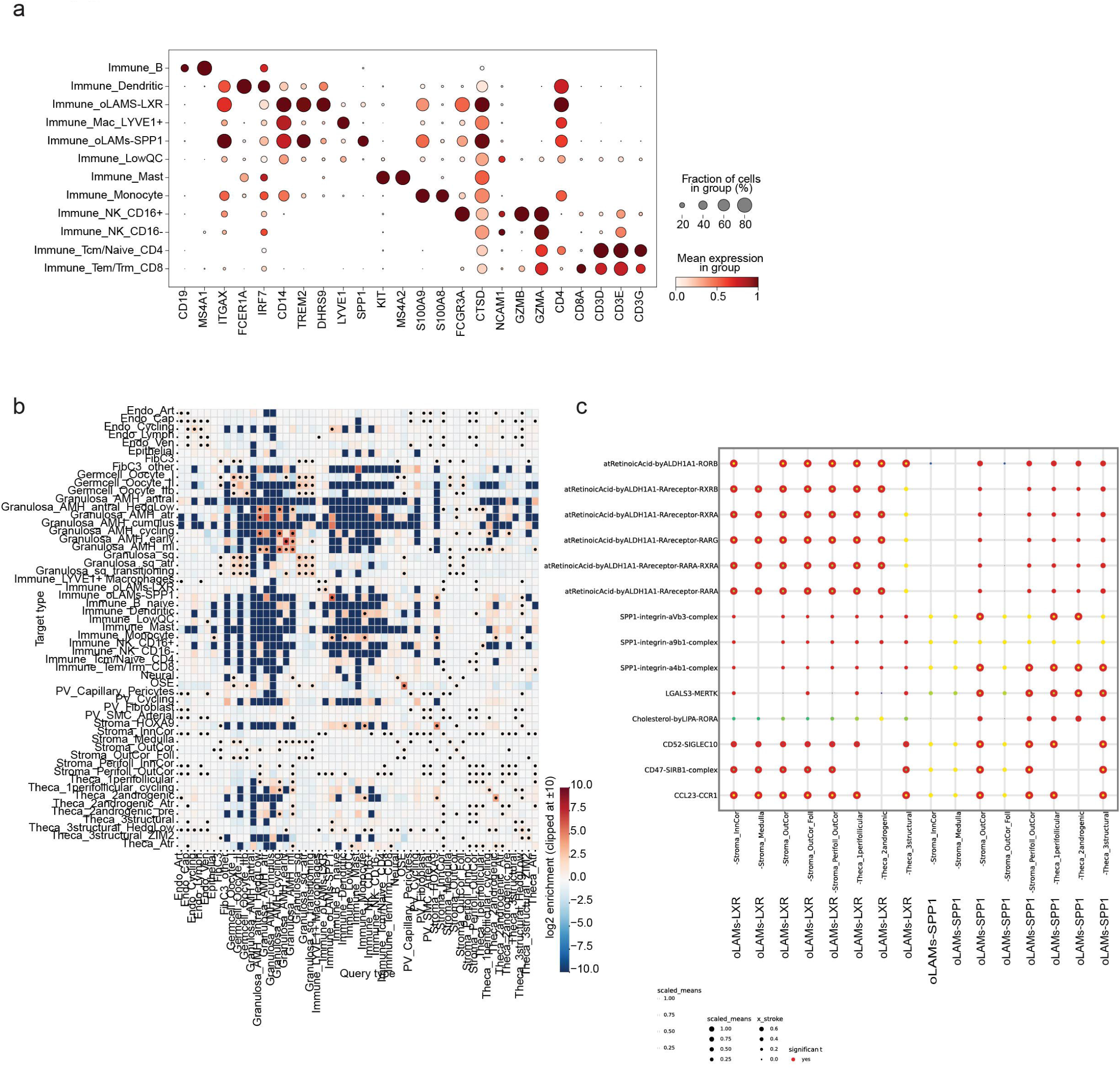
Ovarian macrophages characterisation. **a.** Dotplot showing expression of macrophage-associated markers across immune cell populations from the general UMAP. **b.** A heatmap showing the log₂ enrichment of pairwise cell-cell neighbourhoods across all ovarian cell types, calculated using Fisher’s exact test. **c.** Dotplot showing scaled mean expression of ligand-receptor interaction modules (rows) across macrophage-stromal and macrophage-thecal niche pairs (columns). Each dot corresponds to a ligand-receptor or signalling complex identified by CellPhoneDB, dot colour reflects scaled mean expression, dot size reflects relative interaction strength, and a red-outline stroke represents statistically significant interactions.

## Supplementary Table legends

**Supplementary Table 1. Metadata of donors and samples. a.** Donors information. **b-c.** 10x scRNA-seq libraries for pediatric and adult samples, respectively. **d.** 10x snucRNA-seq libraries generated from sections consecutive to those used for the sequential spatial transcriptomics experiment for pediatric samples. **e.** 10x scATAC-seq libraries for pediatric samples. **f.** 10x scRNA-seq libraries for human ovarian samples from published studies downloaded from publicly available repositories.

**Supplementary Table 2. List of genes measured in the 10x Xenium In Situ Spatial Transcriptomics. a.** List of genes measured in the Xenium Spatial Transcriptomics experiment. **b.** Donor and sample sections information used for the 10x Xenium In Situ Spatial Transcriptomics experiment.

**Supplementary Table 3. OSE characterisation across development. a-b.** Differential gene expression (t-test, one-sided) in Ovarian Surface Epithelial (OSE) cells, comparing fetal samples to pediatric/adult samples in the scRNA-seq and snucRNA-seq data modalities, respectively.

**Supplementary Table 4. Granulosa markers and differentiation dynamics.** a. Differential gene expression (t-test, one-sided) for each granulosa cell type (fine annotations) in the pediatric-adult samples. Only genes expressed in >10% of cells per group are reported. **b**. Marker genes computed with TF-IDF for each granulosa cell type (fine annotations) in the pediatric-adult samples. Only genes expressed in >10% of cells per group are reported **c.** Differentially Accessible Regions/Peaks (DARs) identified using ArchR getMarkerFeatures() function comparing Granulosa_sq versus all other granulosa states combined and each granulosa state vs Granulosa_sq (reporting only DARs with at least one FDR < 20% in one test). **d.** TF-binding motif enrichment within the DARs that were up- or down-regulated in ’c’, computed using a hypergeometric test (reporting only DARs with at least one FDR < 20% in one test).

**Supplementary Table 5. Spatial axes of stroma fibroblasts: Cortico-medullary, Growing Follicular and Atretic Follicular axes. A-c.** Differential Gene Expression Along Spatial Trajectories. Results from tradeSeq::associationTest(), which tests whether the gene expression (modeled by smoothed splines in scRNAseq stromal fibroblasts) is significantly associated with spatial location along the three defined axes: “*Cortico-medullary”* (‘a’), “*Growing Follicle*” (‘b’), and “*Atretic Follicle*” (‘c’) axes, respectively. These spatial trajectories were derived from Xenium data and transferred to scRNA-seq via iss-patcher. **d.** Differential gene expression (one-sided t-test) comparing scRNAseq theca interna cells (pediatric-adult samples) located within 20µm of growing follicles versus those within 20µm of atretic follicles. The 20µm proximity was inferred using the spatial growing and atretic axes transferred from Xenium onto scRNA-seq via iss-patcher. Only genes expressed in >10% of cells per group are reported.

**Supplementary Table 6. Ovarian macrophage characterisation. a.** Differential expression analysis (Wilcoxon rank-sum test) comparing each macrophage subtype against the other. **b.** Fisher’s exact test of macrophage spatial cell type neighbourhood enrichment (knn=3) **c.** Relevant interactions of CellPhoneDB macrophage analysis. Differentially expressed genes were filtered using thresholds (adjusted p-value < 0.001 and log₂FC > 2.0), and only interactions involving these DEGs were retained. For each cell type pair, CellPhoneDB computed mean ligand and receptor expression and performed permutation testing with a 10% expression threshold. CellPhoneDB outputs were further constrained to those ligand–receptor pairs occurring between cell type pairs that showed significant spatial proximity in Fisher’s exact test kNN3 neighbourhood analysis (p-value < 0.05). **D.** scCellFie group-wise comparisons of metabolic task activity scores and Benjamini Hochberg FDR corrected p values between macrophage subtypes to identify significantly altered metabolic programmes.

## Supplementary material

Supplementary Notes File.

