## Supplementary Notes for "Spatial atlas of the ovary identifies molecular events in primordial follicle activation in humans"

**Supplementary Notes.** 2
Supplementary Note 1. H&E-guided annotation, segmentation and registration of Xenium dataset. 2
Supplementary Note 2. Annotation of ovarian cell types across data modalities. 3
2.1. Single-cell and single-nuclei RNA-seq datasets. 3
2.2. Xenium dataset. 4
2.3. scATAC-seq dataset. 5
Supplementary Note 3. Characteristics of ovarian cell types. 6
3.1. Granulosa cells. 6
3.2. Germ cells. 7
3.3. Theca cells. 7
3.4. Mesenchymal stroma cells and fibroblasts. 8
3.5. Ovarian Surface Epithelium and other Epithelial. 10
3.6. Immune. 11
3.7. Endothelial and perivascular. 12
3.8. Neural. 13
Supplementary Note 4. Definition of spatial ovarian axes. 14
4.1. Cortico-Medullary Axis. 14
4.2. Follicular Axes. 15
Supplementary Note 5. Comparison to published single-cell ovarian transcriptomes. 15
**Supplementary References.** 17

### Supplementary Notes.

#### Supplementary Note 1. H&E-guided annotation, segmentation and registration of Xenium dataset.

H&E sections of ovarian tissue were imaged at high resolution and uploaded to the OMERO image management platform. Follicle annotation was performed manually using OMERO's inbuilt annotation tools to identify and classify follicles based on established morphological criteria. Primordial follicles were defined by a single layer of flattened granulosa cells, the primary follicles were characterised by a single layer of cuboidal granulosa cells, while the ones having a mix of flat and round were called transitioning. Secondary follicles exhibited two or more layers of granulosa cells. For more detailed classification of growing follicles, early multilayered follicles were defined as those with a minimum of two and a maximum of three granulosa cell layers. Late multilayered follicles were defined as those with more than three granulosa cell layers but no visible antral space. Antral follicles were identified by the presence of any fluid-filled space indicative of antrum formation. Each follicle was evaluated for structural integrity and classified as atretic if it exhibited characteristic signs of degeneration, including pyknotic nuclei, cytoplasmic shrinkage or fragmentation of the oocyte, and disorganisation or loss of granulosa cell architecture. Annotations were performed by NK and independently reviewed by CTC to ensure consistency, with any discrepancies resolved by consensus. This approach enabled accurate spatial tracking and quantification of follicle stages across patients.

Histology images were registered to xenium images using a nextflow pipeline defined in github.com/uhlmanngroup/ovarian_atlas_xenium_analysis, enabling reproducible data preprocessing and registration that is controlled by a single config file per experiment. Briefly, this pipeline identifies tissue masks for tissue sections in an image, of which there are typically 4-6 sections per image. These masks are then used to crop each section into an independent image (one section per image). This is associated with its respective Xenium experiment as defined in the configuration file and both rigid and non-rigid registrations are performed using Valis[1]. The computed alignment matrix is used to transform omero annotations to the xenium target image. OMERO annotation names are related to Xenium cell segmentation based on if the OMERO annotation overlaps with at least 80% of the Xenium cell. If available, sequential histology sections were also aligned to the xenium experiment. Xenium data and all registered histology were stored within the SpatialData format.

### Supplementary Note 2. Annotation of ovarian cell types across data modalities.

#### 2.1. Single-cell and single-nuclei RNA-seq datasets.

We assembled a comprehensive, across-lifespan single-cell (scRNA-seq) and single-nuclei (snucRNA-seq) atlas of the human ovary using a unified dataset generated at the Wellcome Sanger Institute. Detailed donor metadata, including age and clinical characteristics, are provided in **Supplementary Table** 1. The atlas integrates several developmental windows: (i) second-trimester fetal ovaries (Garcia-Alonso et al.[2]); (ii) pediatric ovarian cortical strips and isolated follicles; and (iii) adult samples obtained via cortical biopsies or whole-organ removal.

To assess the impact of biological and technical variables, we first analysed each library separately and confirmed that all major lineages could be consistently resolved. In particular, we verified that embeddings were not dominated by technical artefacts and ensured that the cell states occupied distinct manifold spaces rather than forming technical continua driven by ambient mRNA or multilineage markers, artifacts typical of low-complexity or technically compromised libraries.

To validate the scRNA-seq findings, we generated an independent snucRNA-seq dataset from OCT-embedded samples. These samples were obtained from consecutive tissue sections matched to those used in the Xenium spatial transcriptomics experiments. Compared with scRNA-seq, snucRNA-seq captures predominantly nuclear RNA displaying differences with the scRNA-seq, yet the majority of marker genes are still specifically detected. A key advantage of snucRNA-seq here is that it enables profiling of tissue sections closely matched to those used for Xenium, supporting cross-modality validation.

Technical details of our computational strategy are provided in the "Analysis of scRNA-seq and snucRNA-seq data" section. Briefly, scRNA-seq and snucRNA-seq atlases were integrated separately using single-cell variational inference (scVI) v0.6.8. In all integrations, donor identity was modelled as the batch variable. scVI learns a batch-corrected latent representation from raw count data, which was used for neighbourhood graph construction and UMAP visualisation. Importantly, batch correction is applied in latent space rather than at the gene level; therefore, downstream gene-level analyses were performed on uncorrected expression values, and gene expression shown in dot plots throughout the manuscript is not batch-corrected.

Cell-type annotation followed a hierarchical strategy. First, all cells were analysed jointly to identify major lineages. Next, we performed lineage-specific scVI re-integration and clustering to derive finer-grained annotations, integrating fetal, paediatric and adult cells within each lineage. For lineages with pronounced fetal vs postnatal differences (for example, granulosa and stromal fibroblast/theca), we additionally repeated the lineage-specific analyses restricted to paediatric and adult cells to ensure that subtler postnatal differences were not masked by the inclusion of transcriptionally distant fetal populations. scVI integration and clustering parameters were considered satisfactory when well-established cell types from different datasets aligned in the embedded space. Cell states were annotated using known marker genes (for previously described populations) and newly identified markers (for previously unreported populations), and marker profiles were checked for consistency across donors to minimise dataset-specific artefacts (see Methods, “Annotation of cell types”). The resulting annotation is represented in the **Supplementary Figure 1c**.

The reported cell types were validated at the transcriptomic level through:

- Individual and integrated analyses of each scRNA-seq dataset, where detection of the same subsets in per-donor and integrated manifolds supported robustness across datasets.
- Analysis of the independent snucRNA-seq dataset, which does not include between-laboratory variation in dissociation protocols; most cell states identified in scRNA-seq were also recovered in snucRNA-seq.
- Integration with spatial transcriptomics (Xenium and Visium CytAssist), enabling in situ localisation of the identified cell states using whole-transcriptome mapping rather than reliance on single marker genes. This is particularly relevant for transitional states defined by coordinated changes in gene expression rather than exclusive markers.

#### 2.2. Xenium dataset.

Xenium In Situ Spatial Transcriptomics data were processed and analysed using a unified spatial framework. Raw Xenium outputs were converted into the SpatialData (0.3.0) open file format using the spatialdata_io (0.1.7) Xenium reader. Registered histology, described above, and associated annotations were then added to this SpatialData object. A NextFlow pipeline then coordinated the merging of cell type annotations and computation of spatial axes for each SpatialData object. These fully annotated SpatialData objects with adjacent histology were uploaded to WebAtlas for interactive visualisation and to the BioImage Archive for download.

To mitigate donor-associated batch effects and obtain a corrected low-dimensional representation of Xenium gene expression, we applied ResolVI, using donor identity as the batch variable. Clustering resolution was adjusted to align with well-established cell types identified in the scRNA-seq and snucRNA-seq datasets.

Annotation of Xenium-derived clusters followed a hierarchical strategy analogous to that used for the dissociated datasets, but with a specific modification to address the technical constraints of 2D spatial segmentation. Since transcript-to-cell assignment can be confounded by the lack of 3D-based segmentation (leading to potential misassignment between neighboring cells) we performed re-analysis and re-clustering based on anatomical and structural proximity rather than global lineage identity alone. Re-analysis was conducted within three spatially defined compartments: (i) vascular structures (endothelial and perivascular cells), (ii) follicular and interstitial compartments (theca, granulosa, oocytes, and stromal fibroblasts), and (iii) immune niches (myeloid and lymphoid populations). In each instance, latent spaces were evaluated using per-sample annotations to ensure the preservation of lineage identity while mitigating donor-specific batch effects. This nested integration strategy allowed us to robustly define major lineages, cell types (broad_annotation) and high-resolution cell states (fine_annotation).

Given the nature of Xenium cell-segmented profiles due to segmentation limitations, we identified marker genes via differential gene expression analysis, and we compared top upregulated genes with marker genes defined in the scRNA-seq and snRNA-seq analyses to guide annotation at both broad and fine levels. Clusters exhibiting mixed lineage signatures, characterised by co-expression of markers from multiple lineages and lacking a coherent identity, were annotated as *Mixed*, consistent with doublet-like or improperly segmented cells.

The reported cell types were validated at the transcriptomic level through:

- Analysis of the consecutive sections of the snucRNA-seq dataset, ensuring the same cell states were consistently identified, were also recovered in snucRNA-seq.
- Indirect annotations using transfer from the scRNAseq with **DOT**.

#### 2.3. scATAC-seq dataset.

scATAC-seq cell type annotation was performed through a multi-step supervised label transfer from the scRNA-seq atlas using the ArchR framework. We followed again a hierarchical approach: first, lineage-level labels were assigned using the addGeneIntegrationMatrix function and refined through cluster-level majority voting; secondly, we repeated this process in a per-lineage manner to project 'broad_annotation' labels onto the ATAC manifold. Rare populations such as OSE were manually refined at high clustering resolution based on specific GeneScoreMatrix markers and populations validated by GeneScore values of marker genes.

The reported cell types were validated at the ATAC level by evaluating the accessibility of bona fide marker genes exclusive to the population.

### Supplementary Note 3. Characteristics of ovarian cell types.

#### 3.1. Granulosa cells.

The granulosa compartment displayed greater heterogeneity and offered a more detailed view of early granulosa differentiation than previous single-cell studies. We list below granulosa states identified and their distinctive markers:

| Cell type - fine annotation | Distinctive markers within lineage | Location | Comments |
|---|---|---|---|
| Granulosa **sq (squamous)** | RDH10+, WNT6+<br>AMH- | Outer cortex<br><br>Primordial follicles | |
| Granulosa **AMH+ transitioning** | RDH10+, WNT6+<br>**AMH+, DHH+** | Outer cortex<br><br>Transitional follicles | |
| Granulosa **AMH+ early** | RDH10-,WNT6+<br>AMH+, DHH+<br>**GSTA1+** | Inner cortex<br><br>Primary follicles | |
| Granulosa **AMH+ ml (multilayered)** | RDH10-,WNT6-<br>AMH+, DHH+,<br>GSTA1+, **IHH+, CYP19A1+** | Inner cortex<br><br>Multilayered/preantral follicles | |
| Granulosa **AMH+ antral** | RDH10-,WNT6-<br>AMH+, DHH+,<br>GSTA1+, IHH+, CYP19A1+<br>**GDF7+, HSD3B2+** | Inner cortex<br><br>Small antral follicles | |
| Granulosa **AMH+ cumulus** | RDH10-,WNT6-<br>AMH+, DHH+,<br>GSTA1+, IHH+, CYP19A1+<br>GDF7+, HSD3B2+<br>**EPPIN+*, WFDC6+*** | Inner cortex<br><br>Antral follicles - cumulus | ***EPPIN & WFDC6** reported to be upregulated in antral follicles by Zhang et al 2021 (Supplementary Figure S5) |
| Granulosa **sq (squamous) atretic** | RDH10+, WNT6+<br>AMH-<br>downregulation of metabolic genes | Outer cortex<br><br>Primordial follicles | Downregulation of housekeeping markers |
| Granulosa **AMH+ cycling** | RDH10-,WNT6-<br>AMH+, DHH+,<br>GSTA1+, IHH+, CYP19A1+<br>**MKI67+** | Inner cortex<br><br>Growing follicles | Upregulation of cycling genes |
| Granulosa **AMH+ atretic** | RDH10-,WNT6-<br>AMH+, DHH+,<br>GSTA1+<br>**downregulation of housekeeping genes** | Inner cortex<br><br>Corpus Atreticum | Downregulation of housekeeping markers |

**Supplementary Note Table 3.1.** Table listing and describing the main characteristics of the identified granulosa cell states (fine annotation).

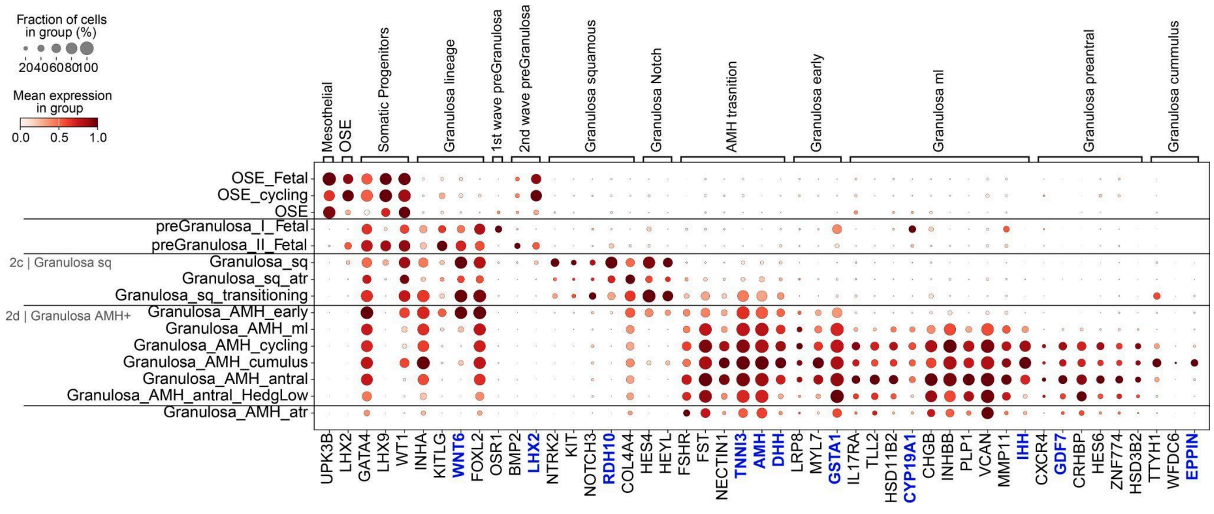


**Supplementary Note Figure 3.1.** Dot plot showing the variance-scaled, log-transformed expression of marker genes (x-axis) characteristic of the granulosa cell states (y-axis; fine annotation) across lifespan in the scRNA-seq dataset. Top-layer groups marker genes by fine annotation.

#### 3.2. Germ cells.

Oocytes (ZP3+/FIGLA+) displayed a relatively homogeneous transcriptome, with variation driven mainly by differing levels of oocyte marker expression and upregulation of primary follicle oocytes signature (Zhang et al), like ZP4. The absence of BMP15+ oocytes supports that these are predominantly from primordial and primary follicles.

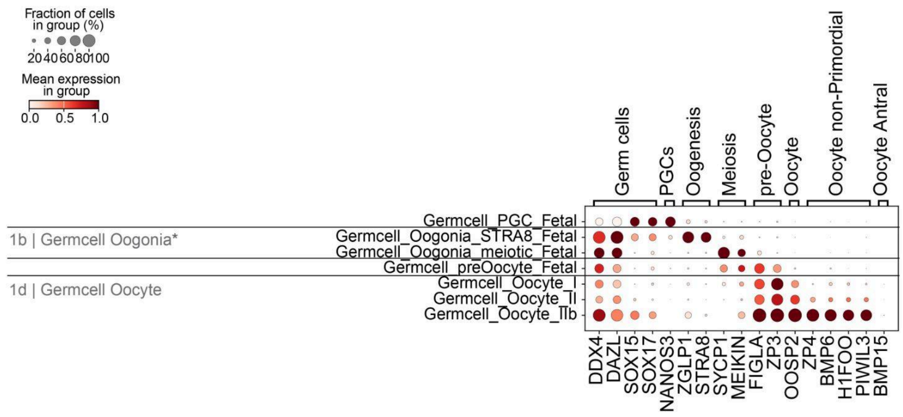


**Supplementary Note Figure 3.2.** Dot plot showing the variance-scaled, log-transformed expression of marker genes (x-axis) characteristic of the germ cell states (y-axis; fine annotation) across lifespan in the scRNA-seq dataset. Top-layer groups marker genes by fine annotation.

#### 3.3. Theca cells.

Theca cell characterisation revealed three major subtypes, which aligned with the ones described by Guahmich et al: *perifollicular theca interna* (HHIP+/THBD+) cells in direct contact with granulosa cells expressing the mechanosensitive gene PIEZO2, likely in response to compression from expanding adjacent granulosa and theca cells; *structural theca externa* (PTCH1+/ACTA2$^{high}$) cells located in the outer layers from the prenatal follicle stage; and *androgenic/pre-androgenic theca* (CYP17A1+/LHCGR+) cells forming aggregates in between externa/interna layers. Integration with Guahmich et al dataset, which profiled theca cells from manually isolated antral follicles, confirmed that our cortical strip theca cells represent the same populations (**Supplementary Figure 5c**).

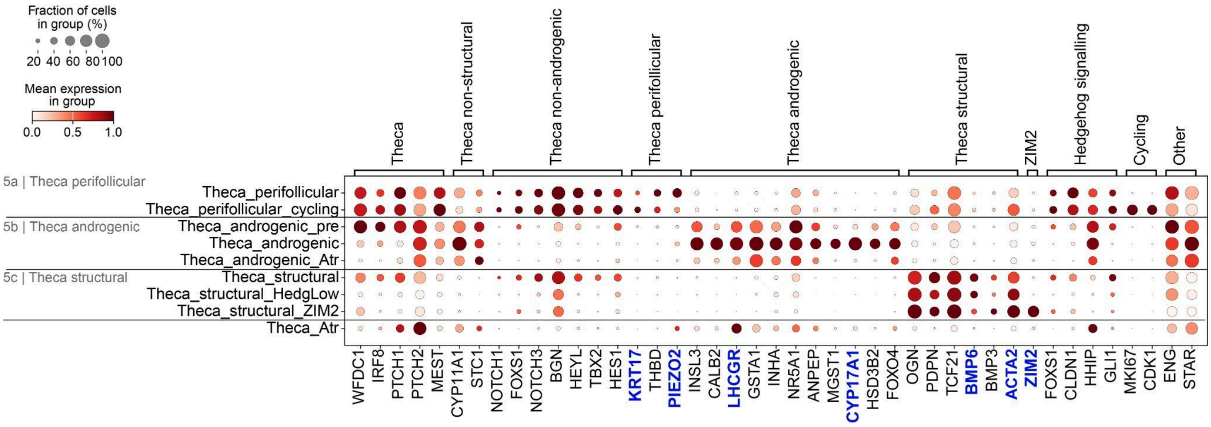


**Supplementary Note Figure 3.3.** Dot plot showing the variance-scaled, log-transformed expression of marker genes (x-axis) characteristic of the theca cell states (y-axis; fine annotation) in postnatal samples in the scRNA-seq dataset. Top-layer groups marker genes by fine annotation.

#### 3.4. Mesenchymal stroma cells and fibroblasts.

We annotated stromal fibroblast using a two-step approach:

(1) First, we derived an initial fine annotation using a conventional scVI-based integration and clustering workflow applied to mesenchymal cells, performed in parallel for both the scRNA-seq (**Supplementary Note Figure 3.5.4**) and Xenium datasets.

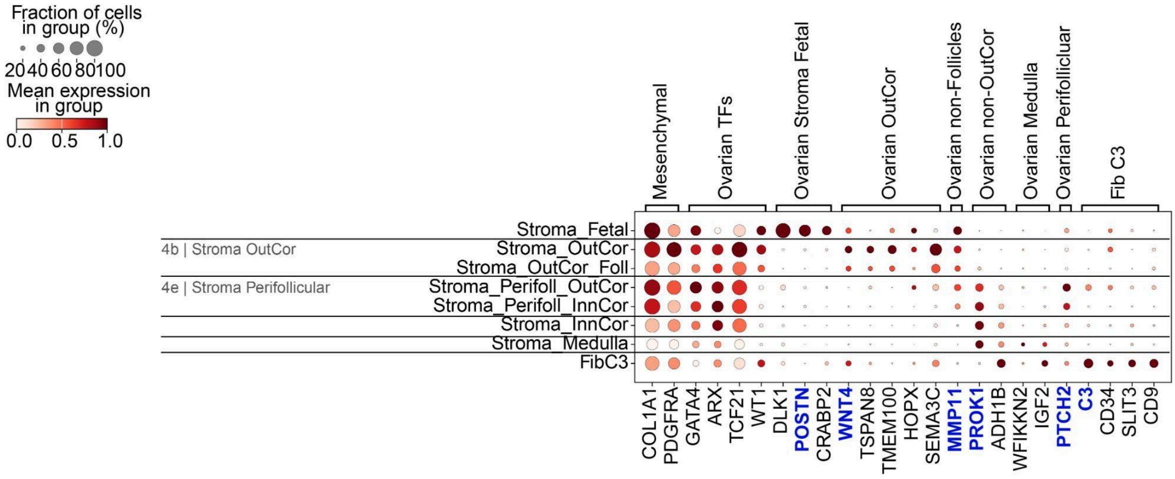


**Supplementary Note Figure 3.4.** Dot plot showing the variance-scaled, log-transformed expression of marker genes (x-axis) characteristic of the stromal fibroblasts (y-axis; fine annotation), directly annotated by classical scVI integration and clustering of the scRNA-seq dataset. Top-layer groups marker genes by fine annotation.

(2) Second, because fibroblast heterogeneity appeared largely continuous and was more readily resolved in Xenium than in dissociated scRNA-seq, we implemented an additional

spatially guided annotation scheme based on each cell's position along the cortico-medullary (CM) axis (**Supplementary Note 4.1**). Briefly, fibroblasts from each tissue section (2–28 years) were mapped onto a CM axis measuring distance from the outermost cortex toward the medulla. This axis uses the fibroblast fine_annotation from point "1" as conserved spatial landmarks/reference points. Then we applied an Organ_Axis[3] inspired approach to propagate a CM axis that standardises spatial distances (i.e. cortical depth) across donors despite differences in ovarian size, sampling region and section orientation. We then used the continuous CM axis to discretise fibroblasts (broad_annotation="Stroma_OutCor', 'Stroma_InnCor', 'Stroma_Medulla') into strata by combining (i) inflection points where broad transcriptomic programmes shift along the gradient (by modelling gene expression along the axis with TradeSeq) and (ii) proximity to primordial follicle granulosa (**Supplementary Note 4.1)**, yielding the fibroblast strata described in the table below (marker visualisation in Supplementary Figure 8g-h, for scRNAseq and Xenium respectively):

| **Fine_annotation (CM-axis-refined)** | Location | Gene expression |
|---|---|---|
| OSE-adjacent fibroblasts | Fibroblasts located immediately beneath the ovarian surface epithelium (OSE).<br><br>Shared transcriptomic programme with WNT4+ secretory fibroblasts. | COL10A1, HOXP<br><br>WNT4, TSPAN8, IGFBP6, TMEM100, SEMA3C |
| WNT4$^{+}$ secretory fibroblasts | Dominant fibroblast population of the outer cortex forming a distinct cortical stratum. They coincide spatially with the dense outer cortex in contact juxtaposed to the primordial follicles. | WNT4, TSPAN8, IGFBP6, TMEM100, SEMA3C |
| Primordial follicle border fibroblasts | A transitional fibroblast state located at the inner boundary of the WNT4$^{+}$ region, flanking primordial follicles. | - KIT expression peaks.<br><br>- Downregulation of WNT4, TSPAN8, IGFBP6, TMEM100, SEMA3C |
| Inner cortical fibroblasts | Fibroblasts located deeper in the cortex. | - Lack expression of WNT4, TSPAN8, IGFBP6, TMEM100, SEMA3C<br><br>- Express SFRP1, LTBP1, EBF1, CXCL12, IGF1/2, PROK1 |
| SMCs-like Inner/ Medullary fibroblasts | Fibroblasts located deeper in the cortex or the medulla, with a SMC-like flavour. | - Lack expression of WNT4, TSPAN8, IGFBP6, TMEM100, SEMA3C<br><br>- Express SFRP1, LTBP1, EBF1, CXCL12, IGF1/2, PROK1<br><br>- Upregulate SMC genes like MYH11, ACT2 |
| Medullary fibroblasts | Fibroblasts located in the medullary region. Fibrotic and with a more stress-like signature in dissociated datasets. | Similar to Inner cortical fibroblasts |

**Supplementary Note Table 3.4.** Table listing and describing the main characteristics of the identified stromal fibroblasts cell states (fine annotation CM-refined).

#### 3.5. Ovarian Surface Epithelium and other Epithelial.

Beyond canonical OSE, we identified additional mesothelial and epithelial-like cells lining cortical cysts. Each subtype was observed in only one or two donors, consistent with donor-specific occurrences. These populations were detected exclusively in the Xenium dataset and were not recovered in the scRNA-seq or snRNA-seq data owing to their low abundance (3,348 of 3,973,414 cells; 0.08% of QC-passing cells). We expect their true frequency to be even lower, as these epithelial populations are cortical, whereas most low-quality cells with compromised QC were medullary. In one donor, we identified mesothelial-lined cysts (UPK3B+/EPCAM-) that expressed gonadal origin markers (WT1, GATA4, LHX9) but lacked LHX2, and showed a transcriptomic signature distinct from the eutopic OSE. Other two donors presented ovarian cortical cysts lined by PAX8+ epithelial cells (EPCAM+), and with a transcription factor profile suggesting a müllerian origin (EMX2+/GATA4-), not an ovarian origin (GATA4+/LHX9+). The prevalence and broader relevance of these cysts remain to be determined in larger cohorts (**Supplementary Note Figure 3.5.2**).

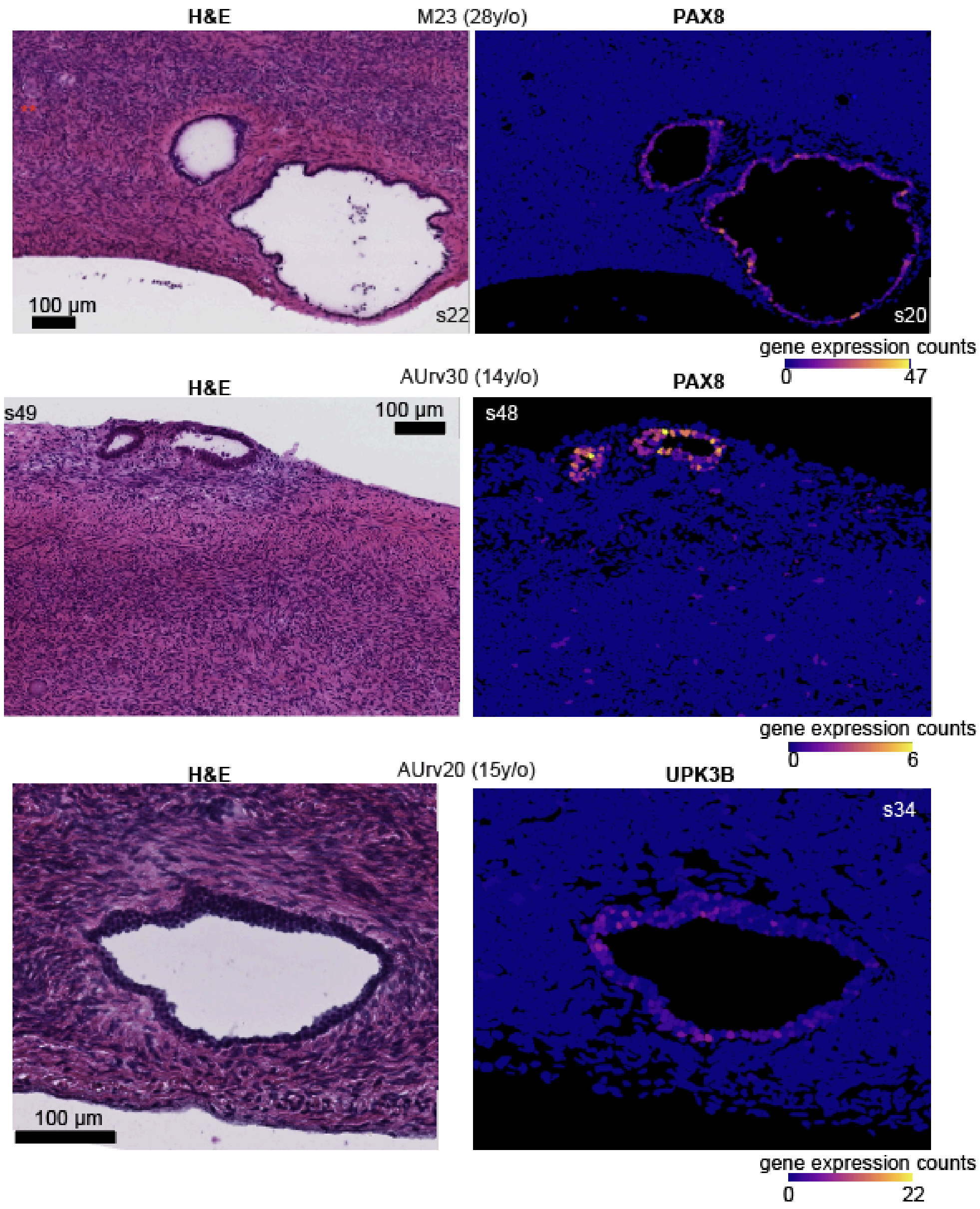


**Supplementary Note Figure 3.5.2.** Hematoxylin and eosin (H&E) staining (left) and gene expression for genes characteristic of the reproductive tract epithelium *PAX8* or mesothelium *UPKB3* (right) marking nonOSE epithelial structures identified in the cortical biopsies of three donors (M23 s20 28y/o; AU30 s48; 14y/o; and AU20 s34 15 y/o) profiled by Xenium spatial transcriptomics.

#### 3.6. Immune.

Immune cell characterisation revealed a structured immune landscape within the ovarian cortex. Three major macrophage subtypes were identified: LYVE1 tissue-resident macrophages (LYVE1+/STAB1+/F13A1+), SPP1+ ovarian lipid-associated macrophages (oLAMS; SPP1+/CD9+), and oLAMs LXR+ macrophages (TREM2+/DHRS9+/CCL18+). Additional myeloid lineages included monocytes (S100A8+/S100A9+/FCN1+) and dendritic cells (CD1C+/FCER1A+/CLEC10A+). Major lymphoid populations comprised B cells (CD79A+/CD79B+) and T/NK lineages, including CD4 T cells (IL7R+/TCF7+), CD8 T cells (CD8A+/CD8B+), and NK CD16–positive and –negative subsets (FCGR3Ahigh or low/KLRD1+/GZMB+/PRF1+). Together these immune populations outline a conserved cortical immune niche involved in surveillance, extracellular matrix turnover, and follicular remodelling (Supplementary Figure).

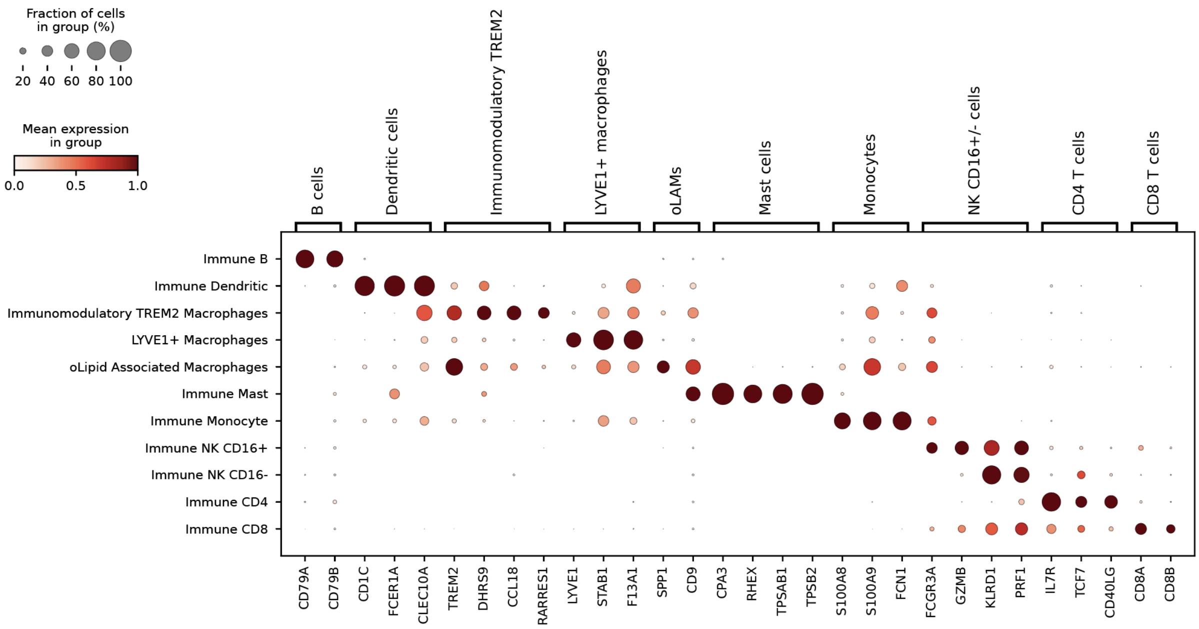


**Supplementary Note Figure 3.6.** Dot plot showing the variance-scaled, log-transformed expression of marker genes (x-axis) characteristic of the immune cells (y-axis; fine annotation)in the scRNA-seq dataset. Top-layer groups marker genes by fine annotation.

Macrophage populations correspond to clusters described in the adult ovary. Briefly, LYVE1$^+$ macrophages corresponds to the previously defined C1QC$^+$ ovarian macrophage cluster, oLAMs-SPP1 corresponds to the SPP1$^+$ macrophage cluster and oLAMs-LXR aligning with the previously defined HLA-DQA1$^+$ ovarian macrophage population.

#### 3.7. Endothelial and perivascular.

The ovarian vasculature was composed of diverse endothelial and supporting mural populations spanning arterial, venous, capillary, and lymphatic networks. Arterial endothelial cells (SEMA3G+/GJA5+/HEY1+) and capillary endothelial cells (VWF+/RGCC+/CA4+) mapped to small vascular structures coursing through the cortex and medulla, while venous endothelial cells (POSTN+/VCAM1+/PLVAP+) delineated broader luminal vessels. Lymphatic endothelial cells (PROX1+/PDPN+/CCL21+) formed a discrete lymphatic network. The perivascular compartment was represented by pericytes (RGS5+/PDGFRB+/TAGLN+) and arterial smooth muscle cells (MYH11+/RERGL+/CNN1+), outlining the mural layers surrounding arterioles and venules.

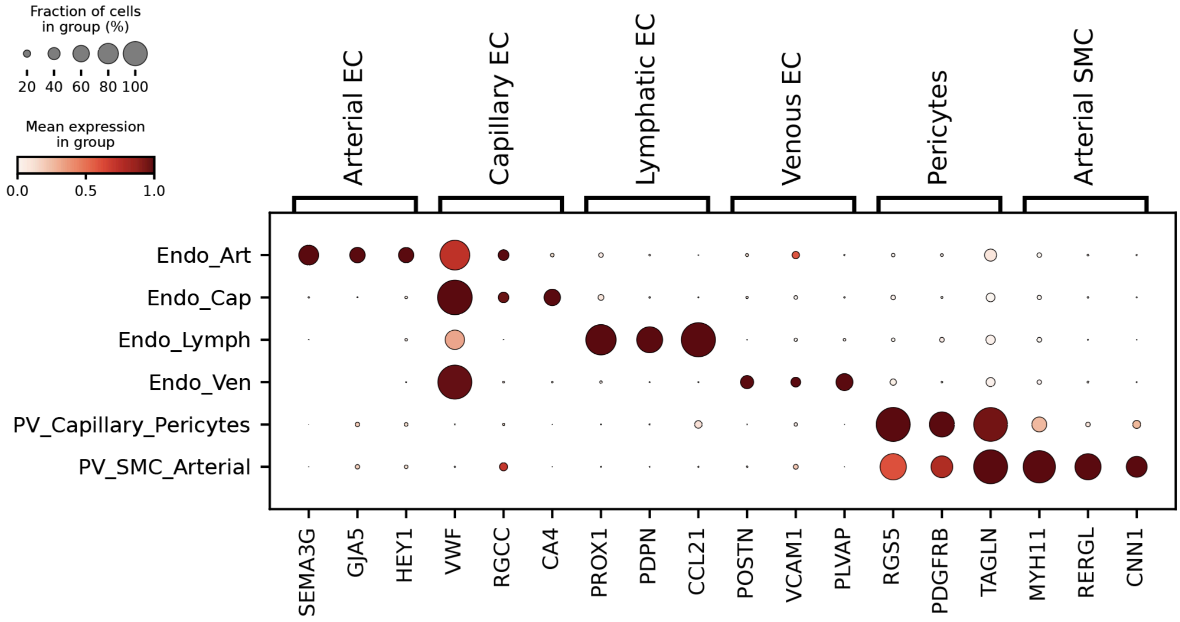


**Supplementary Note Figure 3.7.** Dot plot showing the variance-scaled, log-transformed expression of marker genes (x-axis) characteristic of the endothelial cells (y-axis; broad annotation)in the scRNA-seq dataset. Top-layer groups marker genes by broad annotation.

#### 3.8. Neural.

Neural profiling identified a single glial population corresponding to ovarian Schwann cells, marked by expression of SOX10+, MPZ+, ERBB3+, and NGFR+.

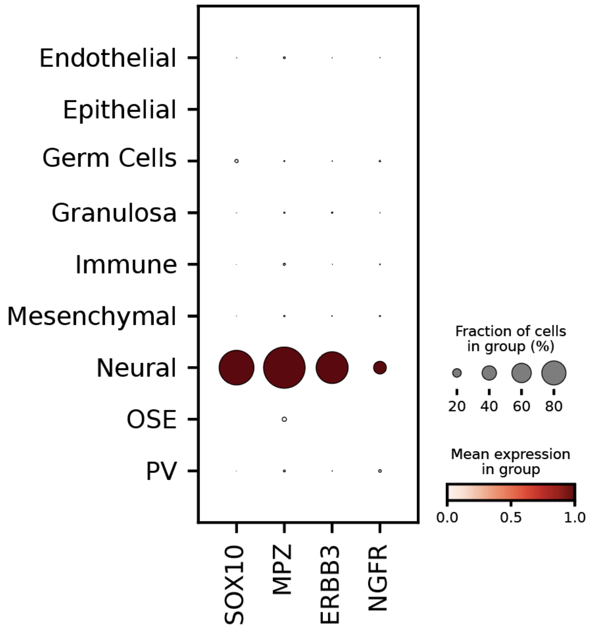


**Supplementary Note Figure 3.8.** Dot plot showing the variance-scaled, log-transformed expression of marker genes (x-axis) characteristic of the neural cells (y-axis; lineage annotation)in the scRNA-seq dataset.

#### Supplementary Note 4. Definition of spatial ovarian axes.

We constructed a common coordinate framework for ovarian tissue using spatial_axis (https://github.com/callum-jpg/spatial_axis), which is conceptually similar to the method described in Yayon et al 2024[3]. A notable distinction in our method is that we derive spatial tissue annotations (i.e. morphological landmarks) from transcriptomic cell type annotation, rather than annotations created manually or from a pixel classifier. Additionally, our method supports any number of classes, allowing us to construct a continuous axis across multiple morphological landmarks.

Briefly, spatial_axis defines for each cell a value in respect to morphological landmarks (for example, stromal layers). For each cell, we calculate the distance to K=15 nearest neighbours for each morphological landmark. These distances are then mean-aggregated, and used to compute the normalised distance for each adjacent morphological landmark. The final spatial axis is then min-max scaled between 0 and 1 for each donor. Before computing any spatial axis, we discarded cells with less than 50 transcripts or annotated as lowQC.

For computation of the Corpus Atreticum and Growing Follicular spatial axes, we only consider K=1 nearest neighbour, providing us with the Euclidean distance to the nearest atretic or growing landmark for every cell within the sample. Prior to finding the distance to these landmarks, we apply a cell filter which only considers cells that are within 25 microns of another cell of the same type. This ensures that atretic or growing follicles, which are composed of multiple landmark cell types, are considered for the axis, rather than sparse single cells.

Spatial axis gradients were first computed in the Xenium space, followed by value transfer to the single-cell space using iss_patcher (https://github.com/Teichlab/iss_patcher). Due to the size of our dataset (>5 million cells), we take a stratified subsample of 0.8 of the lineage cell types for the stromal spatial axis to alleviate memory constraints. Xenium cells with less than 50 transcripts are also dropped prior to running iss_patcher. After transferring the values onto the single-cell space, all growing_follicle_spatial_axis values exceeding 150 μm were masked (set to NA).

##### 4.1. Cortico-Medullary Axis.

For Cortico-Medullary (CM) axis computation, the cell annotation order used was as follows: 'Stroma_OutCor_OSE', 'Stroma_OutCor', 'Stroma_OutCor_Foll', 'Stroma_InnCor', 'Stroma_Medulla'.

After transferring the CM axis onto the cells in the scRNAseq dataset using iss_patcher, we model gene expression dynamics in the stromal fibroblasts along this axis with Tradeseq. We defined genes as spatially variable along the CM axis if they had a Wald statistics > 100, mean log fold change larger than 0.25 and FDR < 0.05. We then used these spatial expression patterns and primordial follicle distribution (i.e. CM axis values in granulosa squamous cells) to annotate fibroblasts niches:

| Fine annotation refined | Description | CM axis range |
|---|---|---|

| | | |
|---|---|---|
| OSE-contact fibroblasts (outer cortex) | Adjacent to the OSE (cortical extreme) | Cells in direct contact with the OSE, receiving an CM axis == 0 (the minimum). |
| WNT4$^{+}$ secretory fibroblasts (outer cortex) | Between OSE and the primordial follicle border. It is situated above the majority (95%) of primordial follicles, meaning it contains up to 5% of them. | This region starts right after the OSE-contact fibroblasts and stops at the point where the CM axis reaches the **5th percentile value** in squamous granulosa cells. CM axis = [0-0.12]. |
| Primordial follicle border fibroblasts (outer cortex) | Niche that hosts the majority (95%) of primordial follicles. | Star and end are calculated as the **5th and 95th percentile values** of the CM axis in squamous granulosa cells, respectively. CM axis = [0.12-0.28] |
| Inner cortex/medulla sub-niches | Remaining portion of the axis toward the Medulla end, subdivided based on gene expression patterns, and displaying a peak in perivascular signature | |
| Inner cortex/medulla SMC-like | **high** expression of smooth muscle / mural markers such as TAGLN, ACTA2, and MYH11 | CM axis = [0.28- 0.5] |
| Inner cortex/medulla | **low** expression of TAGLN, ACTA2, and MYH11 | CM axis = [0.5-0.75] |

**Supplementary Note Table 4.1.** Table listing and describing the main characteristics of the identified stromal fibroblasts cell states (fine annotation CM-refined) and the CM thresholds used for the discretisation.

##### 4.2. Follicular Axes.

Annotations from single-cell RNA-seq were transferred to Xenium cells using DOT (https://github.com/saezlab/DOT). For "healthy growing follicles axis", the cells predicted to be "Theca_1perifollicular_cycling" by DOT were used as histological landmarks to construct a 1D axis. For atretic follicles, the cells annotated as "Theca_atretic" were used as histological landmarks to construct the atretic axis. For each axis, only KNN=1 was considered, providing for each cell in the sample the distance in microns to a growing or atretic follicle. The axis can be translated into microns by multiplying the value per 0.2125 to convert the value into microns.

#### Supplementary Note 5. Comparison to published single-cell ovarian transcriptomes.

| **Reference** | **Cells/tissue profiled** | **Technology** | **Cohort** |
|---|---|---|---|
| Zhang et al 2018 (GSE107746)[4] | Oocyte and granulosa cells enzymatically or mechanically digested from follicles -primordial to preovulatory. | RNA-Seq (pseudo bulk; 10 granulosa cells per reaction). | Reproductive age adults (24-32 years).<br><br>Ovariectomy following sex reassignment surgery or fertility preservation prior to gonadotoxic treatment. |

| | | | |
|---|---|---|---|
| | Follicles were manually isolated from fresh ovarian cortical samples. | | |
| Fan et al 2019 (GSE118127)[5] | Fresh ovarian tissue from inner cortex remnants.<br><br>Follicles manually isolated from inner cortex remnants. | 10X Chromium scRNA-seq | Reproductive age adults.<br><br>Fertility preservation. |
| Wagner et al 2020 (E-MTAb−8381)[6] | Cryopreserved & thawed human ovarian cortical tissue. | 10X Chromium scRNA-seq | Reproductive age adults.<br><br>Ovariectomy following sex reassignment surgery or c-section. |
| Roos 2022 (PRJEB50778)[7] | Preovulatory follicular cells from fresh follicular fluid - IVF | 10X Chromium scRNA-seq | Reproductive age adults undergoing IVF. |
| Guahmich et al 2023 (GSE192722)[8] | Antral follicles resected from the cortex of whole ovaries or from discard material from fertility preservation patients.<br><br>Only follicles deemed to be healthy by morphological appearance. | 10X Chromium scRNA-seq | Reproductive age adults (tissue donors) or fertility preservation adults. |
| Choi 2023 (GSE206143)[9,10] | Follicular aspiration - IVF | 10X Chromium scRNA-seq | Reproductive age adults undergoing IVF. |
| Jones et al 2024 (sc) (GSE230685)[9] | Fresh ovarian tissue (cortex and medulla). | 10X Chromium scRNA-seq | Deceased adult donors of reproductive age. |
| Jones et al 2024 (nanostring)[9] | Ovarian Regions of Interests (ROIs) - cortex, medulla and follicles - from tissue in paraffin. | NanoString GeoMx Spatial transcriptomics. | |

**Supplementary Note Table 5.** Table listing and describing existing adult single cell or spatial datasets profiling adult ovarian tissue and their characteristics.

### Supplementary References.
