## Supplementary material for "Spatial atlas of the ovary identifies molecular events in primordial follicle activation in humans": Figures and legends

#### Contents

**Figures and Legends** 1
**Supplementary Figures Legends** 7

### Figures and Legends

Figure 1

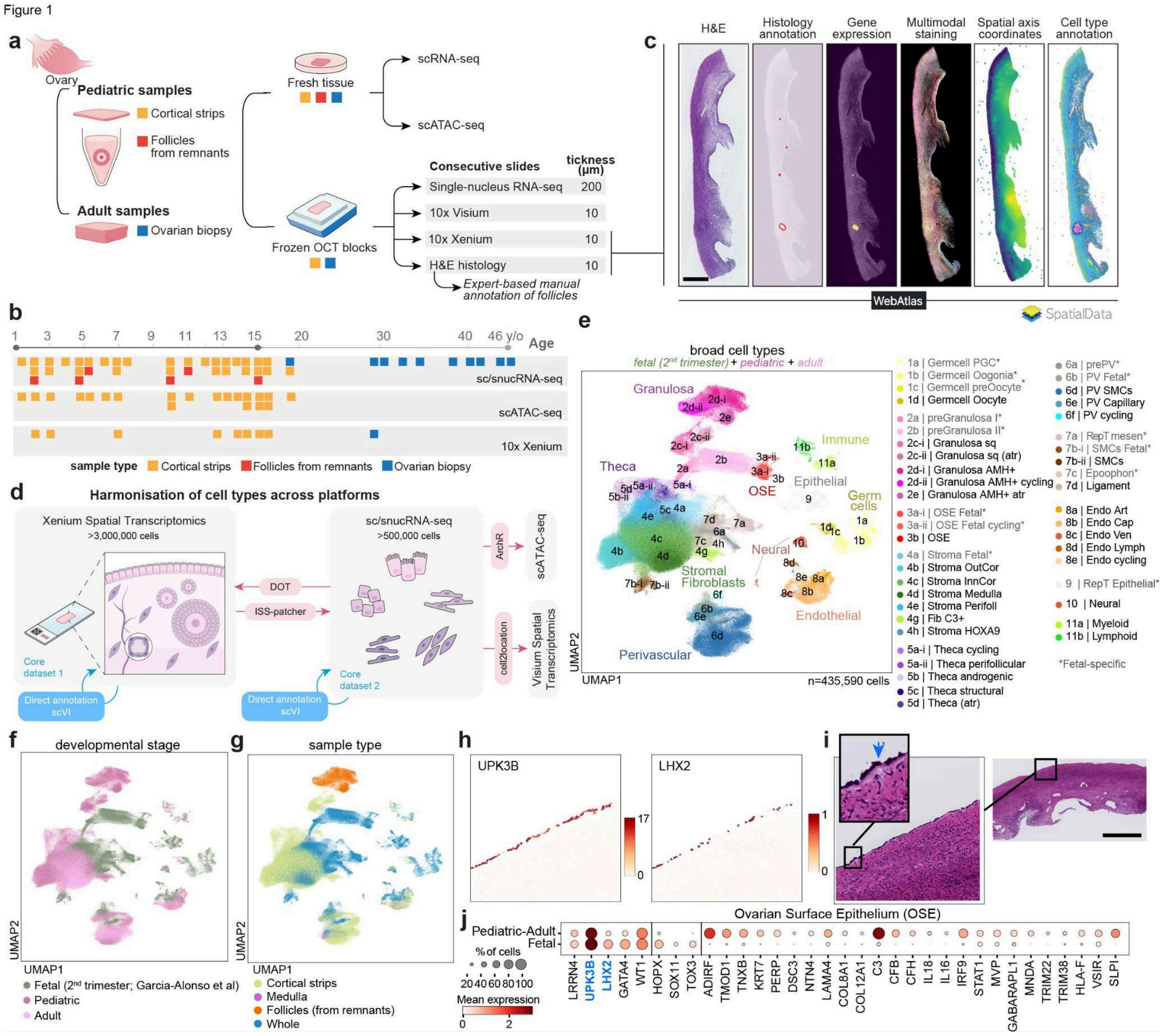


**Figure 1. Study overview. a.** Schematic illustration of samples processing workflow. Color indicates sample type ("orange" for cortical strips, "red" for follicles from the inner cortex remnants, and "blue" for ovarian biopsies). **b.** Diagram summarising the age (x-axis) and sample type composition (color) of our donor cohort along with the technologies used to characterise the donors. **c.** Example Xenium section (AU14; 3 y/o) showing registered cell images and molecular data modalities, available through the WebAtlas web portal and as a SpatialData object. Scale bar = 1mm. **d.** Schematic representation of the pipeline to define and harmonise cell type annotations across data modalities. **e-g.** Batch-corrected Uniform Manifold Approximation and Projection (UMAP) embedding of the scRNA-seq dataset (n = 435,590 cells; n=50 donors) coloured by broad cell type annotation ("e"), developmental

stage (“f”), and sample type (“g”). **h.** Measured gene expression of *UPK3B* and *LHX2* in a representative cortical strip sample (AU30; 14y/o) containing Ovarian Surface Epithelium (OSE) cells. **i.** Hematoxylin and eosin (H&E) staining of the section adjacent to panel “h”, shown at both high and low magnification. Scale bar in low magnification = 1,000μm. **j.** Dotplot showing the log-transformed, min-max normalised expression of selected marker and differentially expressed genes (x-axis) between prenatal and pediatric+adult OSE cells (y-axis).

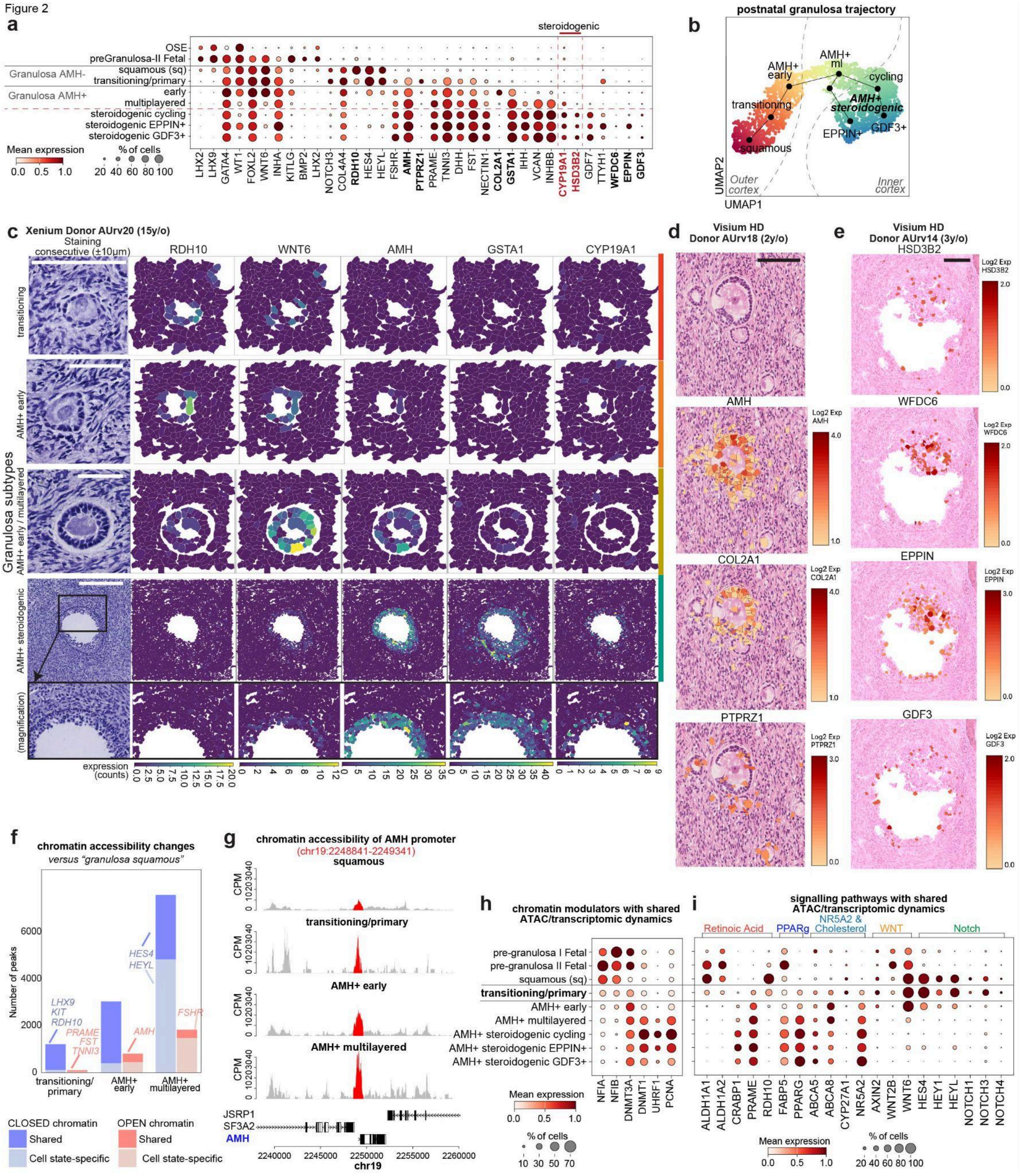


**Figure 2. Molecular changes along early granulosa differentiation. a.** Dotplot showing the log-transformed, min-max normalised expression of selected marker genes (x-axis) for prenatal and pediatric-adult granulosa cell states (y-axis; fine-level annotation). Blue gene names denote literature marker genes. **b.** Batch corrected force directed graph (FDG)

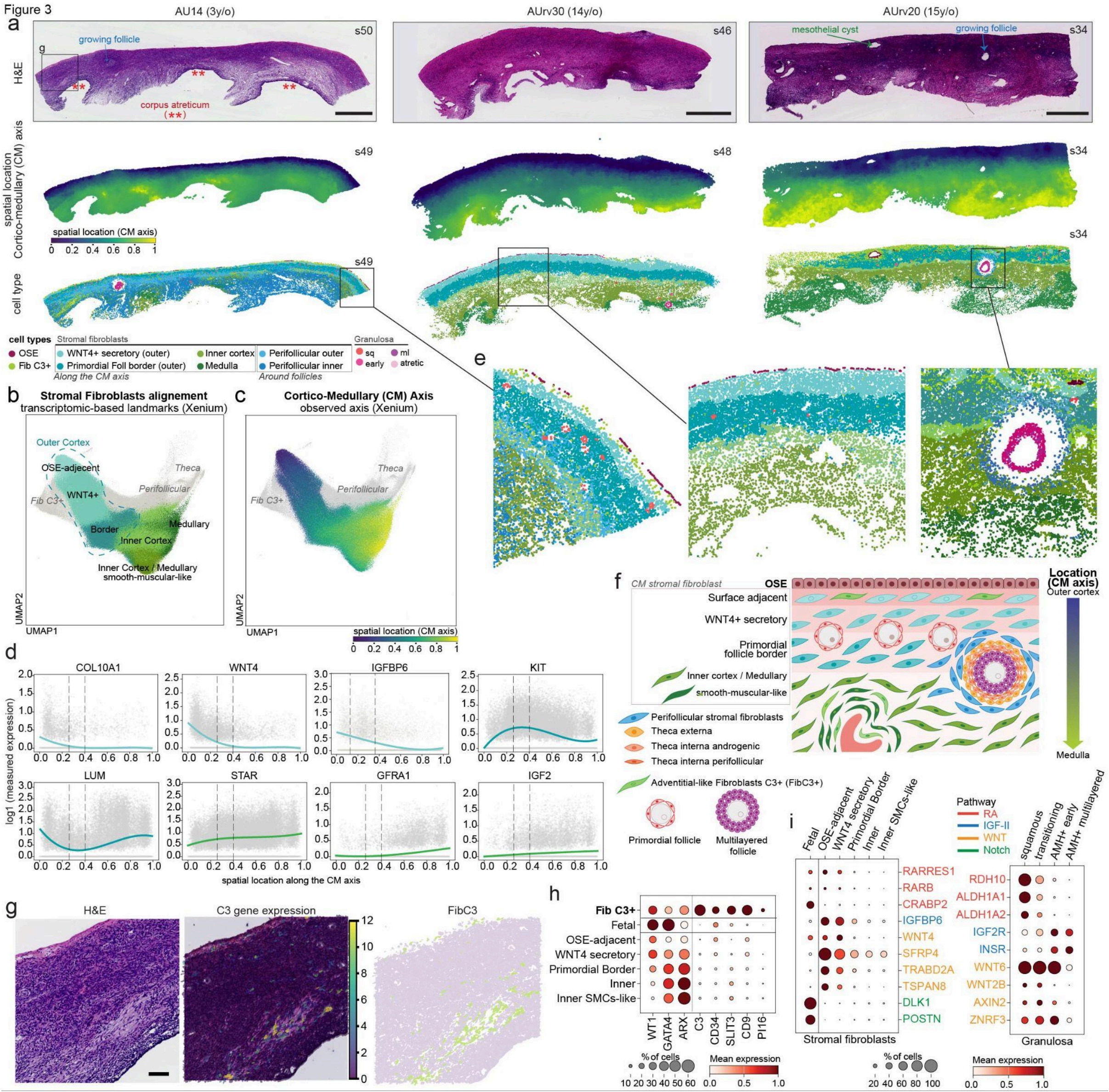


**Figure 3. Fibroblast architecture of the ovarian outer cortex. a.** Three representative ovarian cortical strip sections from pediatric donors (AU14 3y/o; AU30 14y/o and AU20 15y/o) profiled by Xenium, showing adjacent Hematoxylin and eosin (H&E) staining histology (top row), "*Cortico-Medullary axis"* location inferred from the *OrganAxis*-inspired approach (middle row), and cell type annotations for stromal fibroblasts and ovarian surface epithelium (OSE). Scale bar = 1mm **b-c.** Batch-corrected Uniform Manifold Approximation and Projection (UMAP) embedding of the Xenium stromal fibroblast and theca cells coloured by broad cell type annotation, and *"Cortico-medullary Axis"* location inferred from *OrganAxis*-inspired approach (n = cells). **d.** Smoothed splines of Xenium-measured expression (log-transformed) of selected genes spatially dynamic along the "*Cortico-medullary Axis*" in stromal fibroblasts cells. **e.** Higher magnification of selected regions from the sections represented in "a". **f.** Schematic illustration summarising the location of the different stromal fibroblast subtypes along the "*Cortico-medullary Axis*" together with other ovarian cell types. **g.** Hematoxylin and eosin (H&E) staining (left), gene expression for gene *C3* (middle) and cell type annotation of C3+ Fibroblasts (Fib C3+; right) of a representative pediatric cortical strip section (AU14; 3y/o, low magnification in panel "a" left column sample) profiled by Xenium spatial transcriptomics. Scale bar = 100μm. **h.**

Dotplot showing the log-transformed, min-max normalised expression of selected Fib C3+ marker genes (x-axis) in prenatal and pediatric+adult ovarian stromal fibroblasts (y-axis). **i.** Dotplots showing the log-transformed, min-max normalised expression of selected genes (y-axis) in prenatal and pediatric+adult fibroblasts and granulosa cells (x-axis). Genes are colored by their associated signalling pathway.

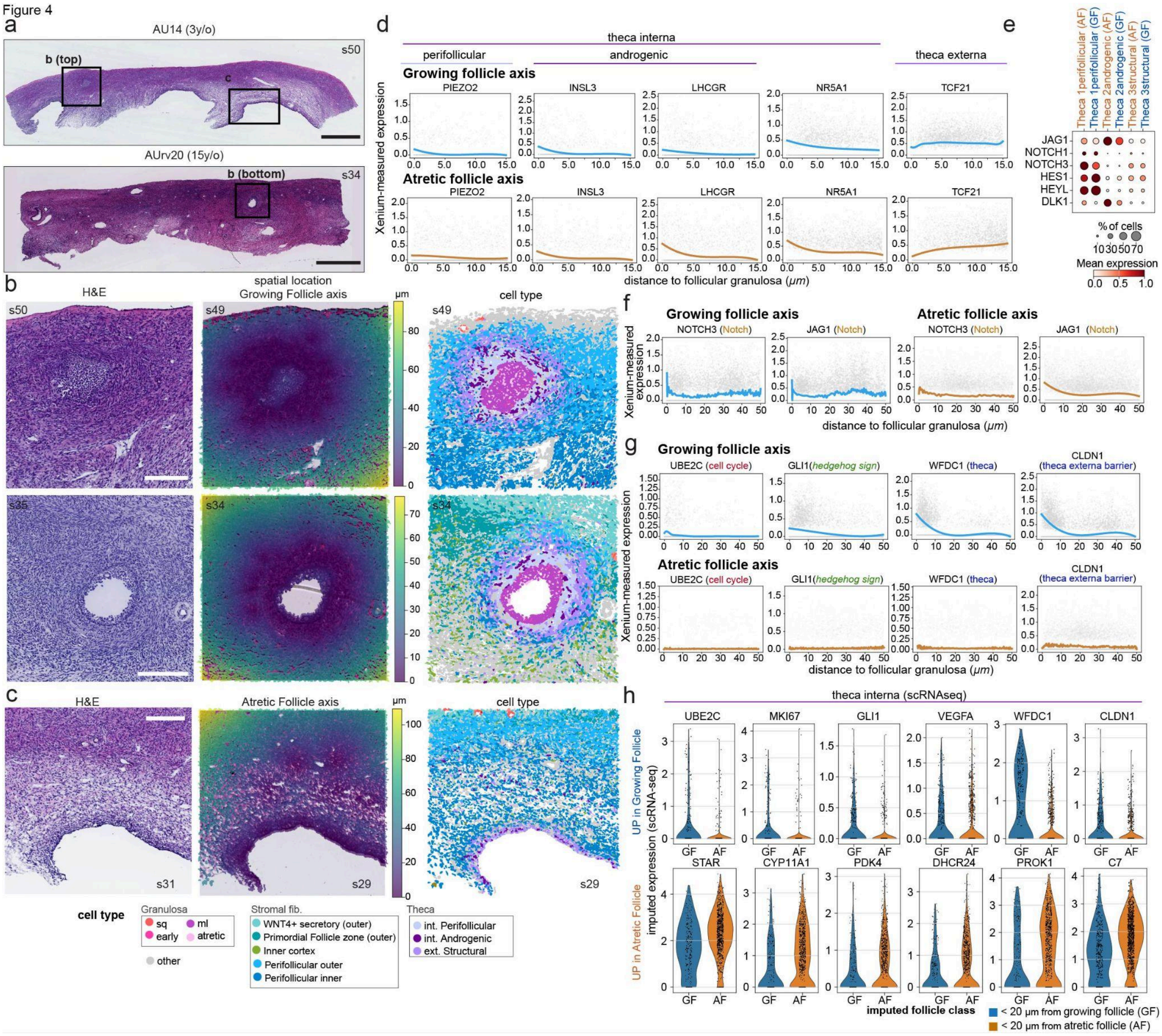


**Figure 4. Perifollicular niche characterisation during growth and atresia. a.** Hematoxylin and eosin (H&E) staining of two representative ovarian cortical strip sections from pediatric donors (AU14 3y/o and AU20 15y/o) containing growing antral and atretic follicles profiled by Xenium. Scale bar = 1mm. **b.** Magnified H&E images (left), inferred growing-follicle axis positions using our common coordinate framework approach (middle), and cell type annotations for fibroblasts, theca and granulosa cells (right) for the two pediatric sections shown in panel "a" (AU14, 3 y/o; AU20, 15 y/o), each containing a growing antral follicle. Scale bar = 200μm. **c.** Magnified H&E image (left), inferred atretic-follicle axis positions using our common coordinate framework approach (middle), and cell type annotations for fibroblasts, theca and granulosa cells (right) for the two pediatric sections shown in panel "a" (AU14, 3 y/o; AU20, 15 y/o), each containing a growing antral follicle. Scale bar = 200μm. **d.** Smoothed splines of Xenium-measured expression (log-transformed) of theca markers. **e.** Dotplot showing the log-transformed, min-max normalised scRNAseq expression of selected NOTCH signalling genes (x-axis) in prenatal and pediatric+adult

theca cells assigned to growing or atretic follicles (y-axis). **f-g.** Smoothed splines of Xenium-measured expression (log-transformed) of selected NOTCH signalling genes ("f") and selected genes with different spatial dynamics between the growing and the atretic follicular axes ("g"). **h.** Boxplots showing the iss-patcher imputed scRNA-seq expression (log-transformed) of differentially expressed genes (y-axis; one-sided t-test; FDR < 0.01) in theca cells classified according to their predicted proximity (< 20 um) to growing or atretic follicles (x-axis).

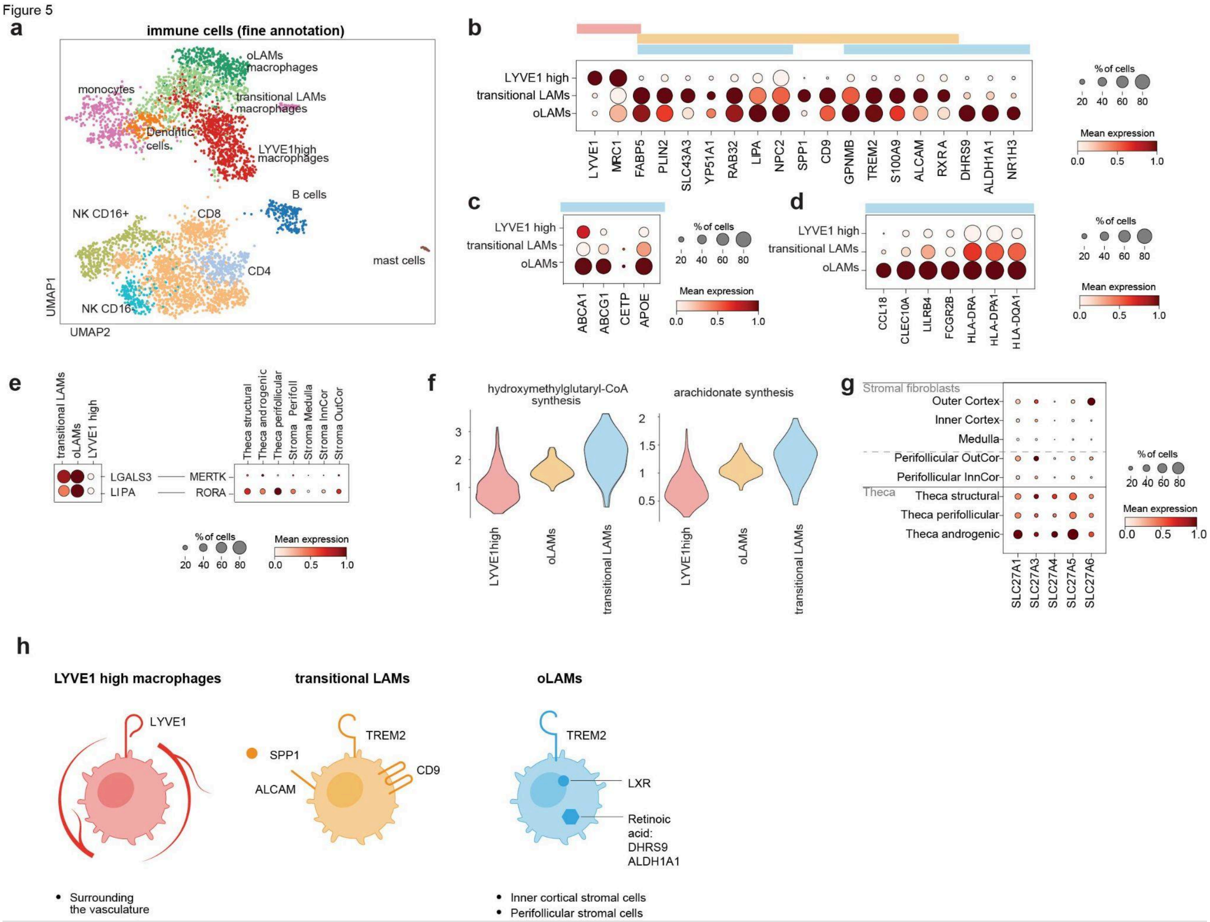


**Figure 5. Ovarian macrophages. a.** Batch-corrected Uniform Manifold Approximation and Projection (UMAP) embedding of the scRNA-seq immune cells from the pediatric-adult ovaries (n = 4,819). **b.** Dotplot showing the log-transformed, min-max normalised expression of macrophage marker genes (x-axis) across the three ovarian macrophage subtypes (y-axis). **c.** Dotplot showing log-transformed, min-max normalised expression of LXR target cholesterol efflux transporters and lipid transport genes (x-axis) across the three ovarian macrophage subtypes (y-axis). **d.** Dotplot showing log-transformed, min-max normalised expression of anti-inflammatory genes and antigen-presentation genes (x-axis) across the three ovarian macrophage subtypes (y-axis)**. e.** Dotplot showing the log-transformed, min-max normalised expression of selected macrophage genes and stromal- and thecal-associated signalling genes (x-axis) across macrophage subsets (y-axis). **f.** Violin plots showing the distribution of scCellFie metabolic scores for key lipid-associated pathways across ovarian macrophage subsets. **g.** Dotplot showing the log-transformed, min-max normalised expression of fatty-acid transporters of the SLC27A family genes (x-axis) across stromal fibroblast and theca cells (y-axis). **h.** Diagram of macrophage subtypes.

### Supplementary Figures Legends

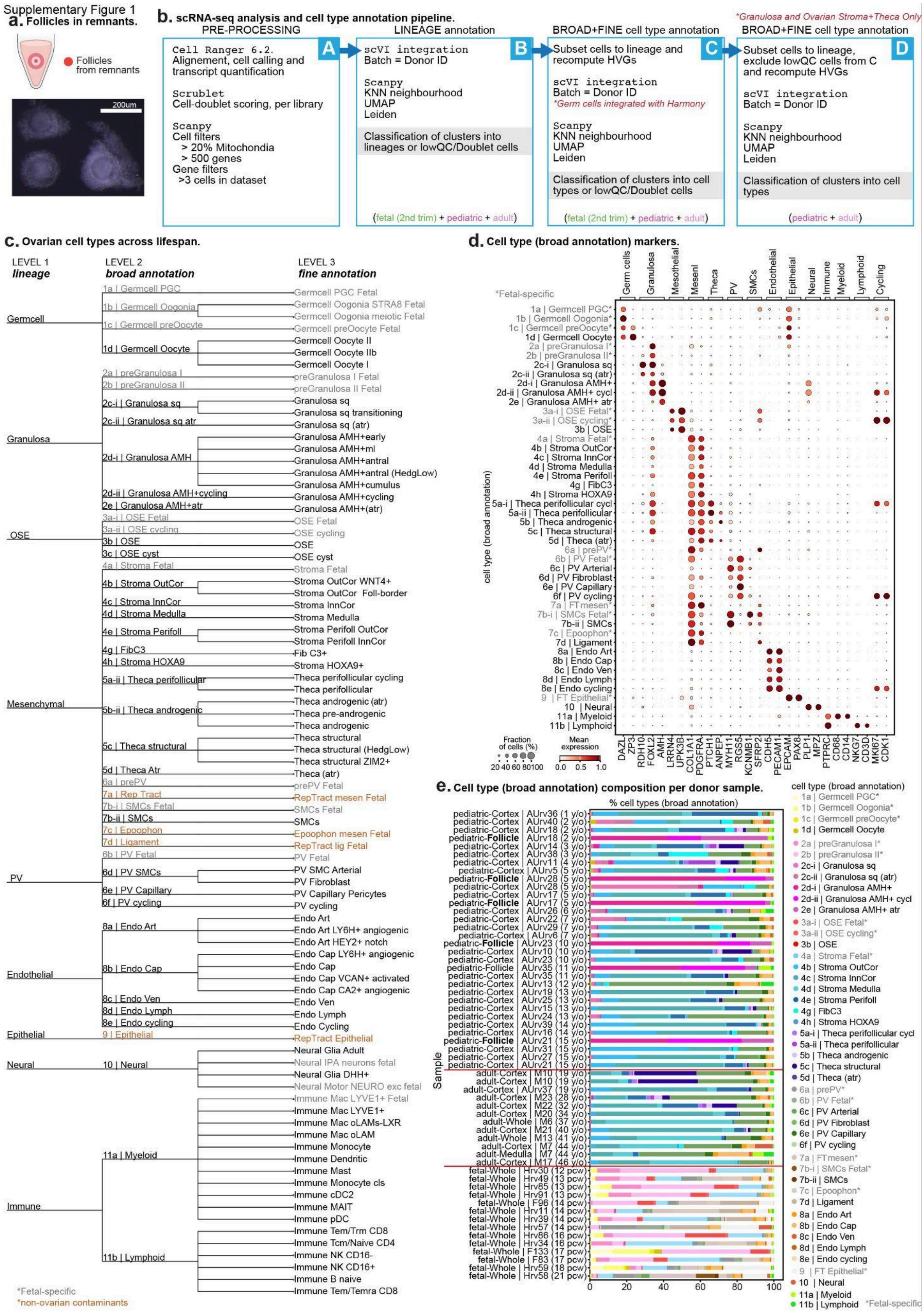


**Supplementary Figure 1. scRNAseq analysis overview and ovarian cell type**

**nomenclature. a.** Representative light microscopy images of intact follicles isolated from remnant fragments from the cortical strip dissection. Scale bar = 200µm. **b.** Schematic representation of the computational workflow used to analyse scRNA-seq data. **c.** Schematic representation of ovarian cell type hierarchy and nomenclature. The tree illustrates the taxonomic organization across three levels of increasing granularity from major lineage to broad and fine-resolution annotations. Tree built with *itol.embl.de*. **d.** Dot plot showing the variance-scaled, log-transformed expression of marker genes (x-axis) characteristic of the annotated cell types (y-axis; broad annotation). Top-layer groups marker genes by major lineage. **e.** Bar plot showing the proportion of cell types per scRNA-seq libraries, grouped by donor and sampling type, and colored by their broad annotation.

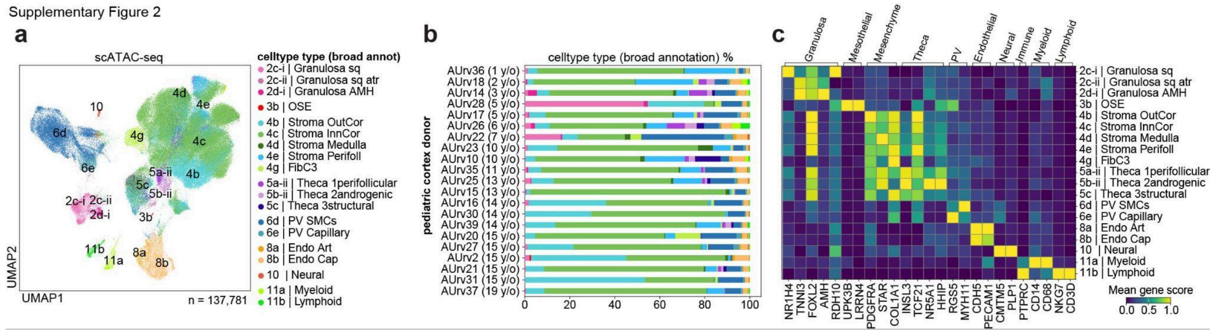


**Supplementary Figure 2. scATACseq analysis overview. a.** Batch-corrected Uniform Manifold Approximation and Projection (UMAP) embedding of the scATAC-seq dataset (n = 137,781 cells) coloured by broad cell type annotation, transferred from scRNA-seq dataset with ArchR. **b.** Bar plot showing the proportion of cell types in the scATAC-seq libraries per donor, colored by their broad annotation. **c.** Heatmap plot showing the variance-scaled, accessibility scores of marker genes (x-axis) characteristic of the annotated cell types (y-axis; broad annotation). Top-layer groups marker genes by major lineage.

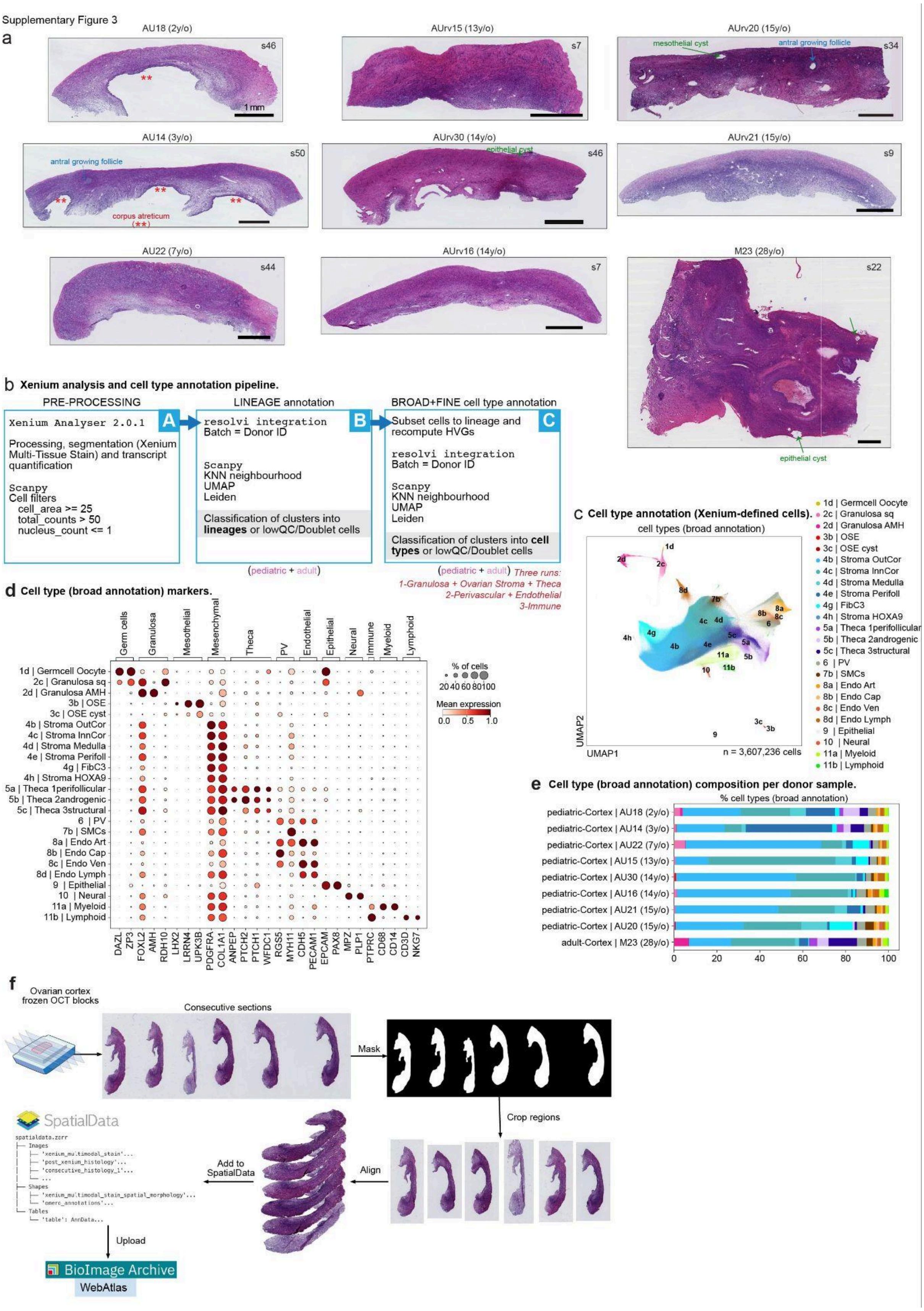


**Supplementary Figure 3. Xenium and imaging analysis overview. a.** Representative Hematoxylin and eosin (H&E) stained sections from each of the eight ovarian cortical strips from pediatric donors and the cortical biopsy from an adult donor profiled by 10x Xenium In

Situ Spatial Transcriptomics; donor ids and age indicated at the title. Scale bar = 1mm. **b.** Schematic representation of the computational workflow used to analyse the transcriptomic profiles from Xenium segmented cells for direct cell type annotation. **c.** Batch-corrected Uniform Manifold Approximation and Projection (UMAP) embedding of the Xenium segmented cells (n = 3,607,236 cells) coloured by broad cell type annotation, annotated based on marker genes. **d.** Dot plot showing the variance-scaled, log-transformed expression of marker genes (x-axis) characteristic of the annotated cell types (y-axis; broad annotation). Top-layer groups marker genes by major lineage. **e.** Bar plot showing the proportion of cell types per donor cortical biopsy, colored by their broad annotation. **f.** Schematic representation of the pipeline for aligning H&E images from consecutive sections and registering multiple spatial modalities (H&E, histological annotation, gene expression, multimodal staining, spatial axis coordinates and cell type annotation) into a SpatialData object, made available through WebAtlas and the BioImage Archive.

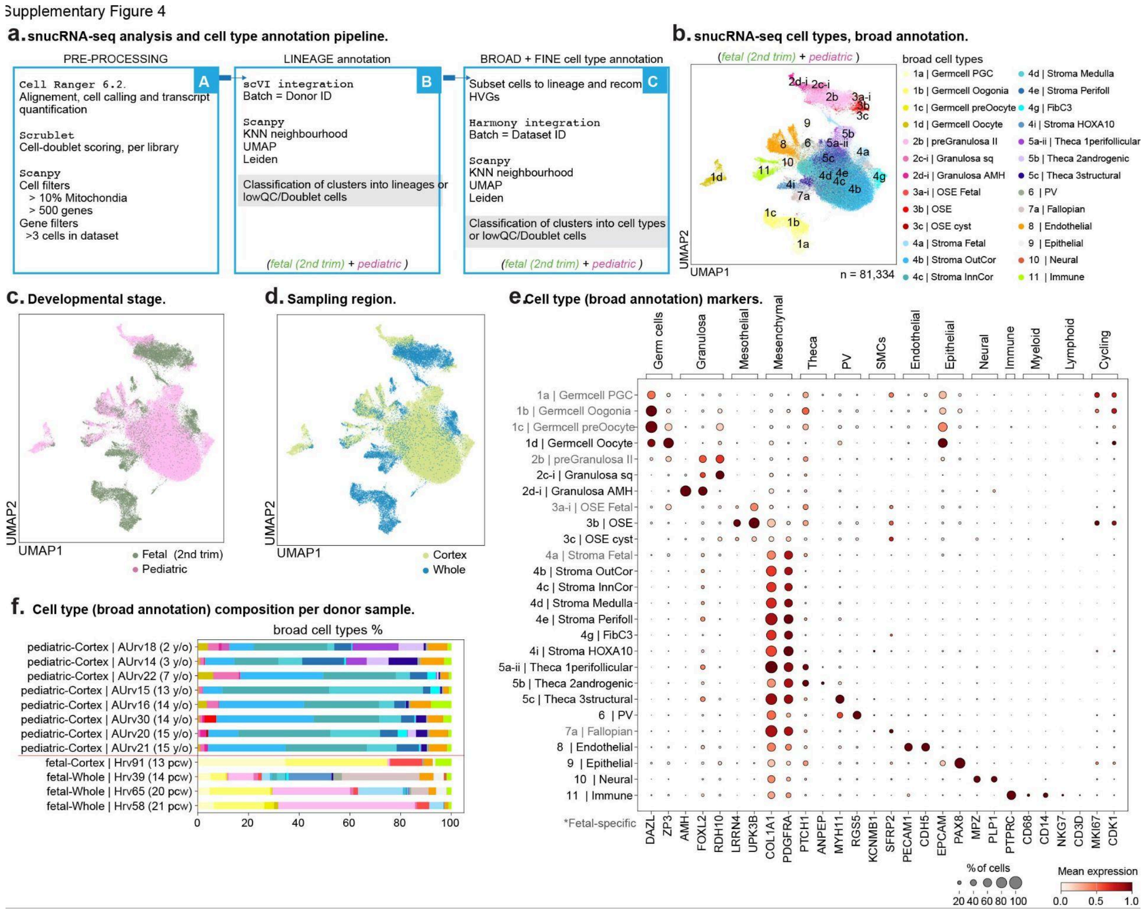


**Supplementary Figure 4. snucRNAseq analysis overview. a.** Schematic representation of the computational workflow used to analyse snucRNA-seq data. **b-d.** Batch-corrected Uniform Manifold Approximation and Projection (UMAP) embedding of the snucRNA-seq dataset (n = 81,334 cells) coloured by broad cell type annotation, annotated based on marker genes ("b"), developmental stage ("c") and "sampling region" ("d"). **e.** Dot plot showing the variance-scaled, log-transformed expression of marker genes (x-axis) characteristic of the annotated cell types (y-axis; broad annotation). Top-layer groups marker genes by major lineage. **f.** Bar plot showing the proportion of cell types in the snucRNA-seq libraries per donor, colored by their broad annotation.

Supplementary Figure 5

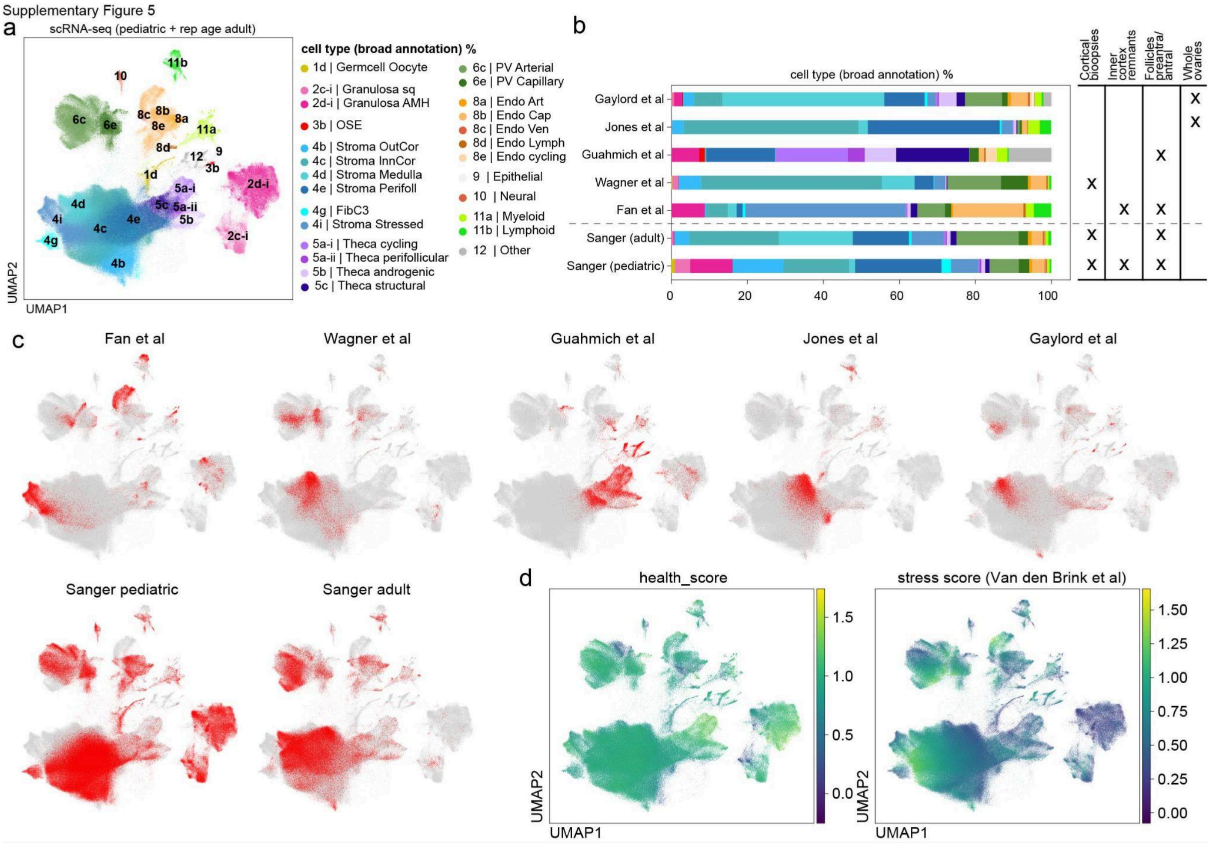


**Supplementary Figure 5. Integration and comparison with existing scRNAseq data from human adult ovaries. a.** Batch-corrected Uniform Manifold Approximation and Projection (UMAP) embedding of our and existing scRNA-seq datasets combined (n = 675,119 cells) coloured by broad cell type annotation, annotated based on marker genes. **b.** Bar plot showing the proportion of cell types in each scRNA-seq dataset, colored by broad annotation (left); and table indicating the biopsy type profiled by each dataset (right). **c.** UMAPs, as in (a), coloured by each of the datasets used in the integration. **d.** UMAPs, as in ("a"), coloured by health score (left) and stress score (right).

Supplementary Figure 6

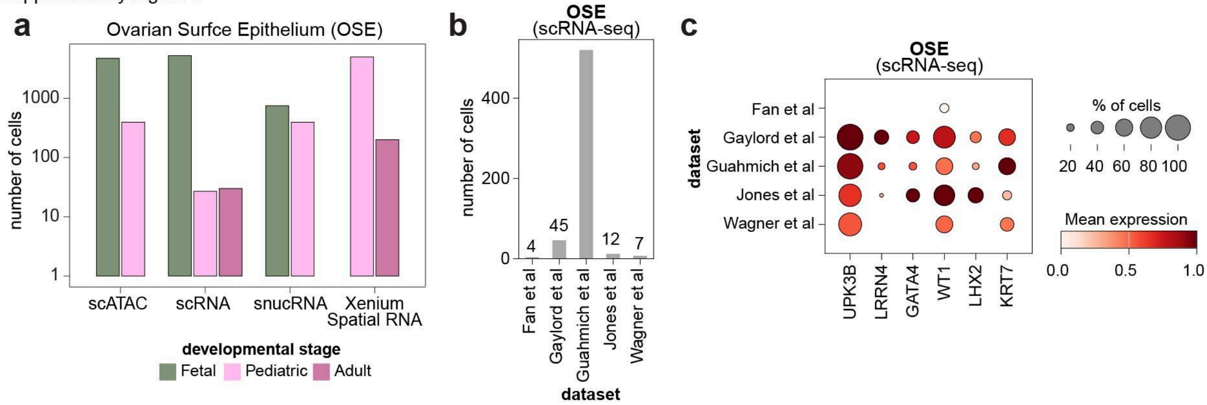


**Supplementary Figure 6. OSE and other aberrant epithelial cells in the ovary. a.** Barplot showing the number of canonical OSE cells identified though development age and data modality. **b**. Bar plots showing the number (y-axis) of OSE cells (broad cells; annotated directly on the integrated UMAP) in each of the publicly available scRNA-seq datasets analysed (x-axis). **c.** Dotplot showing the log-transformed, min-max normalised expression of OSE markers (x-axis) along OSE cells from "b" (y-axis), in each of the publicly available scRNA-seq datasets analysed.

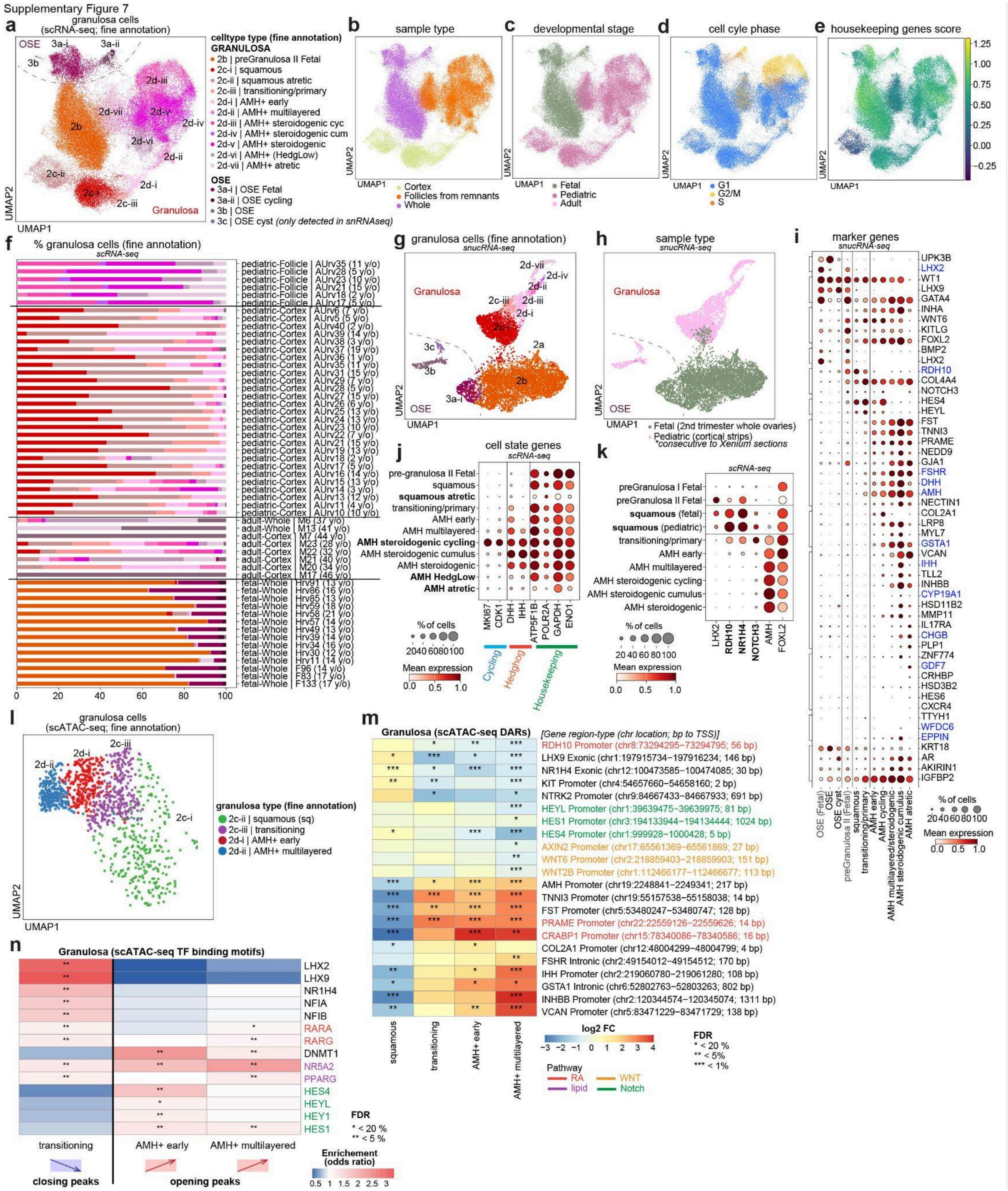


**Supplementary Figure 7. Granulosa cells characterisation. a-e.** Batch-corrected Uniform Manifold Approximation and Projection (UMAP) embedding of the granulosa-annotated cells in the scRNA-seq dataset (n = 63,672 cells) coloured by fine cell type annotation (a), sample type (b), developmental stage (c), cell cycle phase (d) and average expression (score) of housekeeping genes (e). **f.** Bar plot showing the proportion of granulosa and OSE cell types per scRNA-seq library (x-axis), grouped by donor and sampling type (y-axis), and colored by their fine annotation, color legend in "a". **g-h.** UMAP embedding of the granulosa and OSE annotated cells in the snucRNA-seq dataset (n = 8,538 nuclei) coloured by fine cell type annotation (g; color legend in "a") and developmental stage (h). **i.** Dotplot showing the log-transformed, min-max normalised expression of selected marker genes of granulosa and OSE (x-axis) along granulosa and OSE cells (y-axis) in the snucRNA-seq dataset. **j.** Dotplot showing the log-transformed, min-max normalised expression of cell state marker genes (x-axis) along granulosa cells (y-axis) in the scRNA-seq dataset. **k.** Dotplot showing the log-transformed, min-max normalised expression of squamous granulosa marker genes (x-axis) along granulosa cells (y-axis) in the scRNA-seq dataset; with fetal and postnatal squamous granulosa separated. **l.** UMAP embedding of the granulosa cells scATA-seq profiles, coloured by fine cell type annotation, transferred from scRNA-seq dataset with ArchR. **m.** Heatmap showing the log2 fold change for selected "differentially accessible regions" (DARs) in genes relevant to early granulosa differentiation (y-axis), resulting from comparing each granulosa cell state (x-axis) against squamous granulosa cells (for non-squamous granulosa states) or against all remaining granulosa cells (for squamous granulosa). Asterisks denote FDR significance (* < 20%; ** < 5%; *** < 1%). **n.** Heatmap showing the odds ratio enrichment for selected DNA-binding protein motifs relevant to early granulosa differentiation (y-axis), in the DARs downregulated in the transitioning-vs-squamous, in the DARs upregulated in the early-vs-squamous and in the DARs upregulated in the ml-vs-squamous comparisons (x-axis). Asterisks denote FDR significance (* < 20%; ** < 5%).

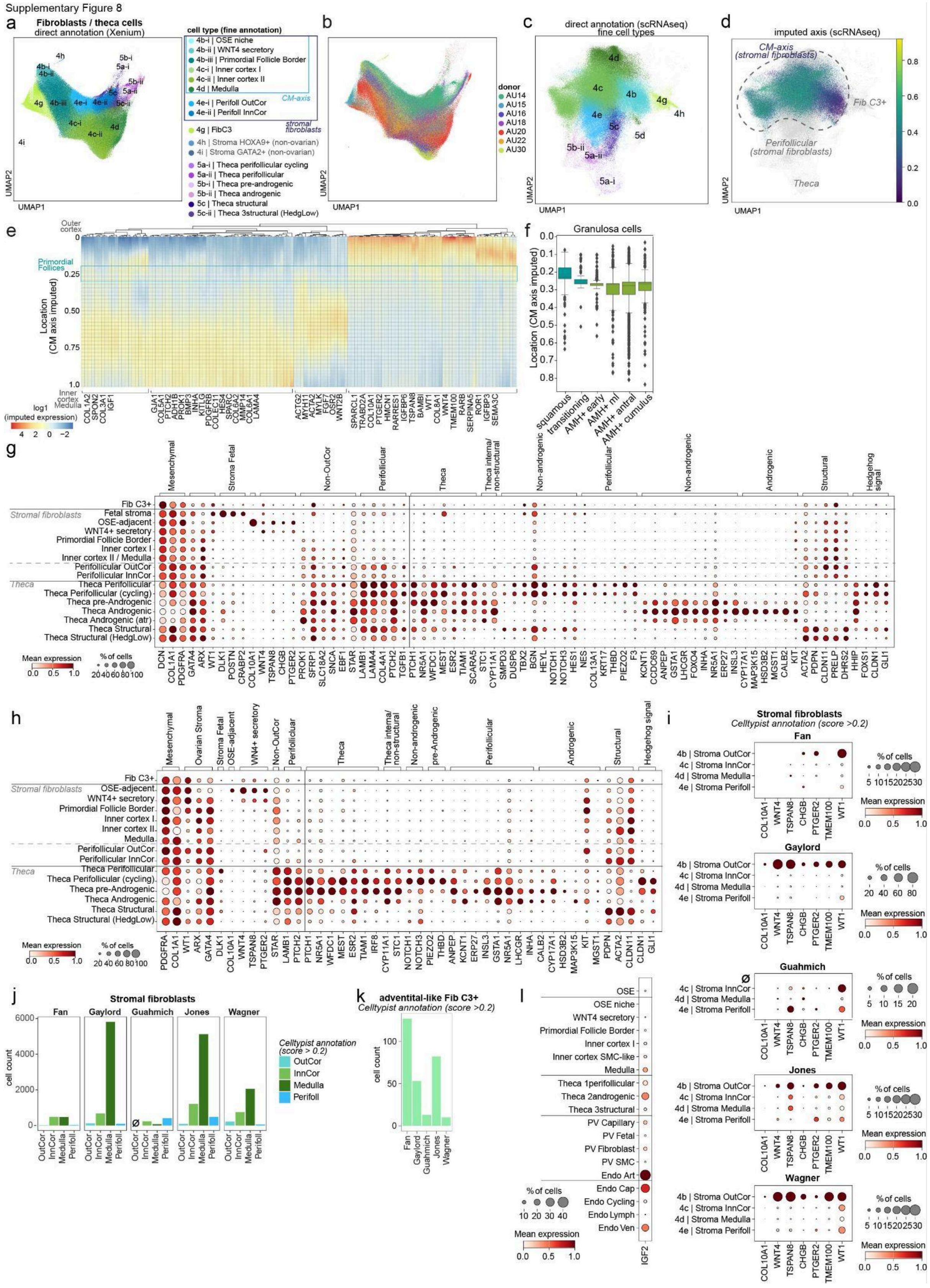


**Supplementary Figure 8. Stromal fibroblast “*Cortico-medullary spatial axis*”. a-b.** Batch-corrected Uniform Manifold Approximation and Projection (UMAP) embedding of the stromal mesenchymal cells in the Xenium dataset coloured by fine cell type annotation, where stromal fibroblasts are coloured according to their assigned fibroblast strata based on their spatial coordinate in the "*Cortico-medullary axis*" ("a") and colored by donor ID ("b"), respectively. **c-d.** UMAP embedding of the stromal mesenchymal cells in the scRNA-seq dataset coloured by fine cell type annotation (same color legend as in "a") and their spatial coordinate along the iss-patcher imputed "*Cortico-medullary axis*". **e.** Heatmap representing the z-score, log-transformed expression of spatially variable genes (x-axis) along the iss-patcher imputed "*Cortico-medullary axis*" (y-axis) identified in stromal fibroblast cells from the scRNA-seq dataset. **f.** Boxplots showing the iss-patcher imputed "*Cortico-medullary axis*" spatial location (y-axis) for granulosa cell subtypes (x-axis). **g-h.** Dotplots showing the log-transformed, min-max normalised expression of cell state marker genes (x-axis) along the stromal fibroblast subtypes (y-axis) in the scRNA-seq dataset ("g") and Xenium segmented cells ("h"). **i.** Dotplots showing the log-transformed, min-max normalised expression of WNT4+ secretory fibroblast markers (x-axis) along stromal fibroblasts subtypes (y-axis; cells annotated via CellTypist label transfer from our scRNAseq dataset), in each of the publicly available scRNA-seq datasets analysed. **j.** Bar plots showing the number (y-axis) of stromal fibroblast subtypes (x-axis; cells annotated via CellTypist label transfer from our scRNAseq dataset), in each of the publicly available scRNA-seq datasets analysed. **k.** Bar plots showing the number (y-axis) of C3+ adventitial-like fibroblasts (Fib C3+; cells annotated directly on the integrated UMAP) in each of the publicly available scRNA-seq datasets analysed (x-axis). **l.** Dotplot showing the log-transformed, min-max normalised expression of *IGF2* (x-axis) along OSE, stromal fibroblast, theca, perivascular (PV) and endothelial subtypes (y-axis) in the scRNA-seq dataset.

Supplementary Figure 9

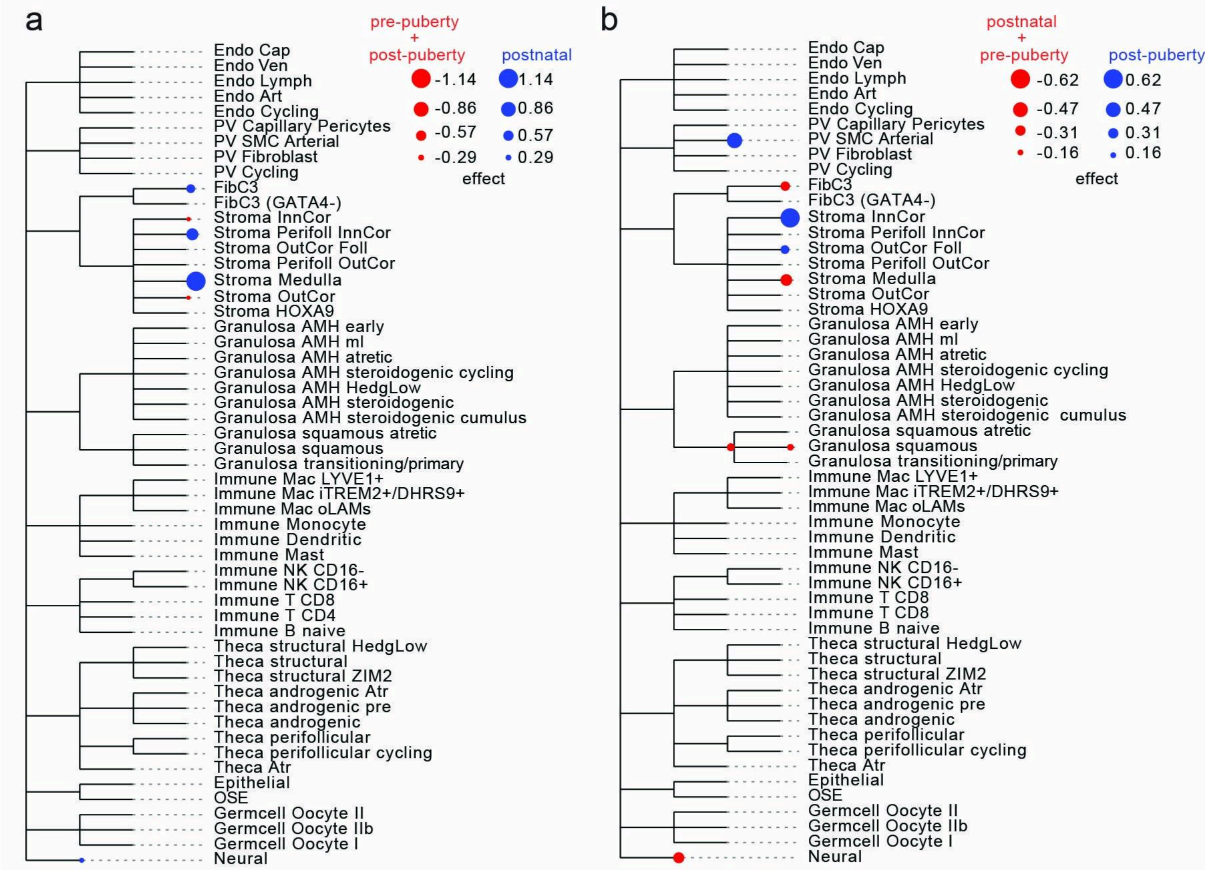


**Supplementary Figure 9. cell type enrichment across childhood stages. a-b.** Cell enrichment analysis at each lifespan stage, using tascCODA with hierarchical smoothing (Φ

= 5.0) in a “lifespan stage vs. the rest” design for postnatal samples vs. pre-puberty + post-puberty samples (“a”) , and post-pubertal samples vs. postnatal + pre-pubertal samples (“b”). Significant positive effect sizes indicate enrichment in the focal stage relative to both other stages. Blue circles denote positive effect sizes (enrichment) and red circles denote negative effect sizes (depletion). Circle size scales with the magnitude of the effect size.

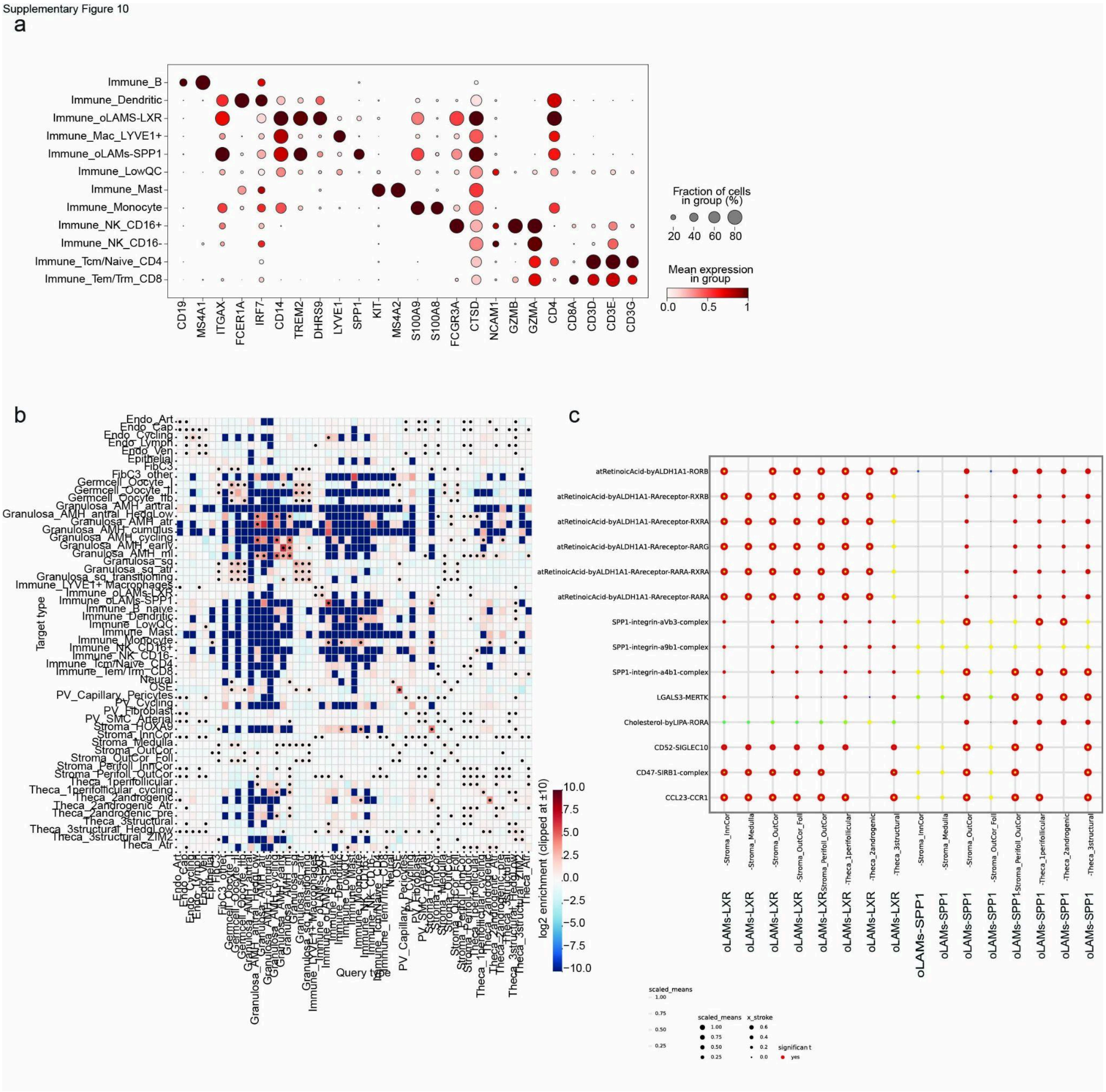


**Supplementary Figure 10. Ovarian macrophages characterisation. a.** Dotplot showing expression of macrophage-associated markers across immune cell populations from the general UMAP. **b.** A heatmap showing the $\log_2$ enrichment of pairwise cell-cell neighbourhoods across all ovarian cell types, calculated using Fisher’s exact test. **c.** Dotplot showing scaled mean expression of ligand-receptor interaction modules (rows) across macrophage-stromal and macrophage-thecal niche pairs (columns). Each dot corresponds to a ligand-receptor or signalling complex identified by CellPhoneDB, dot colour reflects scaled mean expression, dot size reflects relative interaction strength, and a red-outline stroke represents statistically significant interactions.
